# Population genetics meets ecology: a guide to individual-based simulations in continuous landscapes

**DOI:** 10.1101/2024.07.24.604988

**Authors:** Elizabeth T. Chevy, Jiseon Min, Victoria Caudill, Samuel E. Champer, Benjamin C. Haller, Clara T. Rehmann, Chris C. R. Smith, Silas Tittes, Philipp W. Messer, Andrew D. Kern, Sohini Ramachandran, Peter L. Ralph

**Author notes:** These authors contributed equally to the paper (Elizabeth T. Chevy, Jiseon Min).

## Abstract

Individual-based simulation has become an increasingly crucial tool for many fields of population biology. However, continuous geography is important to many applications, and implementing realistic and stable simulations in continuous space presents a variety of difficulties, from modeling choices to computational efficiency. This paper aims to be a practical guide to spatial simulation, helping researchers to implement individual-based simulations and avoid common pitfalls. To do this, we delve into mechanisms of mating, reproduction, density-dependent feedback, and dispersal, all of which may vary across the landscape, discuss how these affect population dynamics, and describe how to parameterize simulations in convenient ways (for instance, to achieve a desired population density). We also demonstrate how to implement these models using the current version of the individual-based simulator, SLiM. We additionally discuss natural selection – in particular, how genetic variation can affect demographic processes. Finally, we provide four short vignettes: simulations of pikas that shift their range up a mountain as temperatures rise; mosquitoes that live in rivers as juveniles and experience seasonally changing habitat; cane toads that expand across Australia, reaching 120 million individuals; and monarch butterflies whose populations are regulated by an explicitly modeled resource (milkweed).

## 1 Introduction

Explicit spatial models are indispensable for understanding how species live, interact, and evolve across geographic landscapes. However, formulating sensible models of demography in continuous space is fraught with pitfalls and choices unfamiliar to many researchers interested in spatial modeling. For instance, in the commonly-used, nonspatial Wright–Fisher model the population size is directly specified. However, in spatial models with locally-defined dynamics the number of individuals is a stochastic, emergent property. It takes some expertise to coerce the model to produce a desired equilibrium size. Even fundamental population genetic concepts such as selection coefficients cease to have a single obvious interpretation in a spatial context. Implementing an individual-based simulation requires great specificity – choices must be made regarding many mechanisms and parameters. Simulated organisms must separately give birth and die, unlike in more abstract theory where often only the net effects of birth minus death enter (*e.g*., Cantrell and Cosner, 2004). Here we present modeling strategies for implementing individual-based simulations in explicit geographic space. These include spatial movement, such as dispersal, as well as spatial interactions, such as the feedback between local population density and net reproductive rate that is necessary to avoid unbounded growth.

Why individual-based simulations, and why in continuous space? Since real individuals are discrete, and live in continuous space, such simulations can in principle more accurately model many real-world situations. Some phenomena simply require individual-based simulations (*e.g*., locally adaptive genotypes).

Non-individual-based models can be more computationally efficient, but at an often unknown cost to accuracy (Stillman et al., 2015). Similarly, discretized spatial landscapes are usually associated with model assumptions that do not provide a consistent approximation to continuous-space dynamics (e.g., Barton et al., 2002). For instance, Battey et al. (2020) showed that some aspects of genetic variation in fine grids of randomly-mating demes of fixed size were irreconcilably different from continuous-space models: in fact, discretization error increased with finer grids. Given these effects of discretization, it is in our experience simpler to move directly to continuous space. Indeed, individual-based simulations in continuous space may even require *less* modeling expertise than other strategies, because it is usually relatively easy to come up with order-of-magnitude estimates of aspects of an organism’s life cycle from natural history data which can be used to parameterize an individual-based simulation. More abstracted modeling frameworks can involve analytical approximations, compound parameters, and other technicalities that require careful checking and mathematical experience. For instance, it is more difficult to translate “offspring disperse around 100m” to a migration rate between adjacent grid cells than to directly input a mean dispersal parameter of 100m. Although abstract models with fewer choices might feel more general for theoretical work or methods development, this can be a false generality, as more work is required after the fact to determine to which real-world organisms a given result or method is applicable.

Explicit individual-based population models are not new to ecology (DeAngelis and Yurek, 2017; Grimm, 1999). A great deal of ecological work has sought to quantify the effects of density-dependent demographic feedback. For a single species, negative feedback between population density and growth rate is necessary to avoid the population growing without bound, although there are a great many ways to set this up in practice (De Wit, 1960; Beverton and Holt, 1957; Ellner et al., 2016). Demographers have a deep understanding of how to describe and parameterize the statistical properties of birth and death, and what the emergent consequences are for population growth, lifespan, age distribution, and long-term fitness (Tuljapurkar, 2013). Although temporal stochasticity is relatively well-understood in demography (Tuljapurkar, 1989), the consequences of spatial heterogeneity – particularly outside of metapopulation models (Hanski, 1997) – have received less attention.

Geography can have strong effects on patterns of genetic variation (Wright, 1943; Malécot, 1969; Rousset, 2000; Charlesworth et al., 2003; Battey et al., 2020; Min et al., 2022) and on evolutionary processes (Felsenstein, 1976; Uecker et al., 2014; Savolainen et al., 2007). Genetic differentiation is shaped by the movement of individuals, and hence distance, geographical features, and the spatio-temporal history of the species (Hewitt, 2011; Rosenberg et al., 2005; Ramachandran et al., 2005). Geography is therefore not only important, but also a relatively untapped source of information to inform inference (Bradburd and Ralph, 2019). However, it is difficult to obtain analytical predictions from spatial population genetics models (Felsenstein, 1975; Barton et al., 2002). Most spatial work in population genetics uses partial differential equations that do not represent genetic differentiation (Slatkin, 1973; Barton, 1979; Sedghifar et al., 2016; Etheridge et al., 2024), or specifically look at the fronts of population expansions (e.g., Barton et al., 2013; Paulose et al., 2019; Nullmeier and Hallatschek, 2013; Etheridge and Penington, 2022).

Simulation has long been a useful tool in the study of populations (Grimm, 1999; DeAngelis and Yurek, 2017), particularly for the purposes of prediction (for example, for population viability analysis, see Dunning Jr et al., 1995) – even predating common usage of digital computers (Pearson, 1960). Simulations are also useful for inference, ranging from exploratory studies to training for deep learning. Their use depends on their computational cost: many modern machine learning methods are only feasible with relatively fast simulations. Introducing geographic space increases computational complexity, making the task of producing training data more difficult (but see Smith et al., 2023; Champer et al., 2021). Today there are several sophisticated software suites targeted at just the sort of landscape-scale simulations we discuss here, with both discretized spatial models (Landguth et al., 2017; Bocedi et al., 2021; Landguth and Cushman, 2010; Schumaker and Brookes, 2018; Neuenschwander et al., 2018; Rebaudo et al., 2013) and continuous spatial coordinates (Haller and Messer, 2023).

### 1.1 How to use this paper

This paper is intended as a guide to the territory of spatial modeling, with a focus on individual-based simulations in discrete time on continuous geographic space.

The paper has two broad parts: The first part (Sections 2 through 8) describes modeling strategies to capture ecological and geographic processes in individual-based models. The second part, **??**, provides concrete examples of how to combine these modeling strategies to simulate a biological system.

First we tackle population regulation (Section 2), as this is the topic which in our experience is the greatest barrier to new modelers. We then describe our strategy for parameterizing and analyzing spatial simulations (Section 3). Next, we cover spatial movement and mate choice (Sections 4 and 5 respectively), highlighting some interesting challenges. Subsequently, we discuss how to implement spatial heterogeneity, and further concepts in population regulation, including stochasticity (Section 6), and finally natural selection (Section 8).

We hope this material will be useful to researchers who wish to produce simulations that are – at least roughly – modeled on concrete empirical systems. This includes: empirical researchers wanting to explore plausibility of hypotheses or power of study designs, methods developers who want to test their methods on realistic spatial models, theoreticians wishing to explore spatial models, and managers wishing to explore alternative scenarios. So, we aim to make it easy to build a model starting from those quantities that we generally have good estimates of for particular organisms. This differs from the modeling philosophy in much theoretical work, which often begins from a “simplest possible” model.

To give practitioners a head-start on simulating the models we describe, the main text is accompanied by Boxes containing code that implements each modeling concept. The code may be run with the program SLiM (Haller and Messer, 2023), a flexible and powerful individual-based eco-evolutionary simulator that, as of the newly-released version 4.2, includes a full set of tools for modeling interactions between multiple species across geographic landscapes. We recommend that readers open the minimal SLiM template (provided in Appendix A and at https://github.com/kr-colab/spatial_sims_standard) in SLiM’s GUI to experiment with while reading. The online code repository also contains SLiM scripts for each of the Case Studies.

## 2 Population dynamics and density

We start by describing how to maintain a stable population. Consider an extremely simple spatial simulation: organisms are asexual, and do not move during their lifespan. Each time step, each adult gives birth to a Poisson(*f*) number of offspring, that each disperse to a random location, whose displacement from their parent’s location is drawn from a Normal distribution with mean 0 and standard deviation *σ*_*D*_. Then, each individual dies with a probability *µ*.

Simulating this model, its flaw is immediately apparent as it runs: either all individuals rapidly disappear (if *f*≤ *µ*), or the computer grinds to a halt as the number of individuals explodes (if *f > µ*). We need some kind of *density-dependent feedback* to maintain a stable population size – in other words, we need the net population growth rate to change from positive to negative as the population density grows past some point. When population density in an area is high, either birth rates need to decrease or death rates need to increase.

To provide density-dependent feedback, we first need a notion of “population density” at a point in space, a measure of the number of individuals nearby per unit area. Let *K* denote the desired equilibrium population density (in individuals per unit area). A general way to define *n*(***x***), the population density around a location ***x***, is to specify an *interaction kernel, ρ*(***x***), which is a nonnegative function with **∫** *ρ*(***x***)***dx*** = 1; an *interaction scale σ*_*X*_ ; and then if the two dimensional locations of the individuals in the population are ***x***_**1**_, …, ***x***_***n***_, define *n*(***x***) by

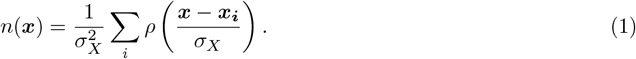

Since 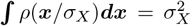,the value of *n*(***x***) is in units of individuals per unit area. A common choice for *ρ*(***x***) is the Gaussian density function. One concrete interpretation is that if *ρ*((***x*** − ***y***)*/σ*_*X*_) gives the proportion of time that an individual at ***y*** spends near ***x***, then *n*(***x***) is proportional to the time spent by *all* individuals near ***x***. (More concretely, ∫_*A*_ *n*(***x***)***dx*** is the total amount of time spent by all individuals in the region *A*.)

Now suppose that in each time step of the model, each individual has a chance to produce offspring. The number of offspring depends on the individual’s location. An individual at ***x*** produces a random number of offspring with a mean of *f* (*n*(***x***)*/K*), where *f* is the *birth rate* or *fecundity*. Each juvenile then disperses to a nearby location whose displacement from ***x*** is chosen from a given probability distribution. Then, all individuals (including those just born) survive to the next time step with probability 1 − *µ*(*n*(***x***)*/K*), where *µ*(*u*) is the *mortality rate* at scaled density *u*. This way, birth and death rates depend on an individual’s location ***x*** via the smoothed population density *n*(***x***), scaled by a parameter *K* that controls the equilibrium density. We build on this general form throughout the paper to describe how life history, geography, and selection make individual fecundity and mortality rates more complex.

### 2.1 Equilibrium population density

Will this model stabilize, and if so, to what density? We expect an equilibrium when births balance deaths. If we define the local *per capita* net reproductive rate (the expected increase due to birth minus the decrease due to death, per individual) to be

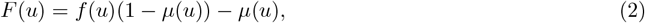

then we expect an equilibrium at a density of *n*_*_, solving *F* (*n*_*_*/K*) = 0. Note that newborns are subjected to the same mortality rate *µ*(*u*) as the rest of the population, so the expected increase is *f* (*u*)(1 − *µ*(*u*)) rather than *f* (*u*) in equation (2). For this reason, it is convenient to choose the functional forms of *f* and *µ* so that *F* (1) = 0, in which case we expect the equilibrium *n*_*_ to be roughly equal to *K*. (However, we will see in Sections 6, 7.1, and AppendixB.4 that often *K* is not *exactly* the equilibrium density.) In addition to *F* (1) = 0, for the population to be stable we also need *F* ^*′*^(1) *<* 0 so that the net reproductive rate decreases with density near to the equilibrium. The argument *u* in *f* (*u*), *µ*(*u*), and *F* (*u*) is the *scaled* population density *u* = *n*(***x***)*/K*, written this way so that it is easy to control the equilibrium population density by simply changing *K*, independently of other factors.

A brief note on what we have done here: it might seem most natural to set up a simulation using fecundity and mortality rates based on empirical observation. If so, then mean population density would be an emergent property of the simulation – in other words, our script would not have a parameter *K* that we could directly adjust. However, to do this we need empirical estimates of how fecundity and mortality depend on local density, which are very difficult to obtain. A much more common situation is to have estimates of fecundity and mortality rates *and population density* at equilibrium. The parameterization we outline here, in which *K* is a directly tunable rather than emergent parameter, is designed to make this use case natural.

### 2.1.1 Functional forms

The next question is: what forms should we use for the birth and death rate functions *f* (*u*) and *µ*(*u*)? There is surprisingly little guidance from the theoretical or empirical literature. Population models mostly come in two flavors: “phenomenological” (or “top-down”) models that only consider the net reproductive rate, *F* (*u*); and “mechanistic” (or “bottom-up”) models that explicitly consider birth and death separately (Geritz and Kisdi, 2012). We need a mechanistic model, since our approach here is individual-based, but most literature on density dependence uses phenomenological models (for notable exceptions, see Coulson et al. (2008), or integral projection models Ellner et al. (2016)). Matrix population models (Caswell, 2000) are (mostly) mechanistic and widely used for management, but rarely incorporate density dependence. Estimating the functional forms of density dependence from empirical data is a logistically and statistically daunting task, even without considering environmental variation. As we intend to simulate from our model, we will simply choose a mathematically convenient form, reducing the problem to estimating parameters given that functional form. Perhaps the most obvious strategy is to multiply the base fecundity or survival rate by a function of the local density that decreases when density is high. Appendix C works through examples using common functional forms for *F* (*u*).

### 2.2 Regulation by mortality

Suppose we would like to use density-dependent feedback only on mortality. In this case, the average number of offspring is constant: we may say *f* (*u*) = *f* . Rearranging equation (2), we see the survival probability in a location experiencing a scaled population density *u* is

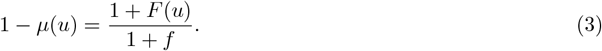

The Beverton–Holt form for the net effects of density (Beverton and Holt, 1957) would have that *F* (*u*) is proportional to (1 + *a*)*/*(1 + *au*) − 1, for some constant *a* that controls the strength of the density-dependent feedback. So, to set up the model to have “Beverton–Holt” feedback, we plug in *F* (*u*) = (1 + *a*)*/*(1 + *au*) − 1 and obtain that

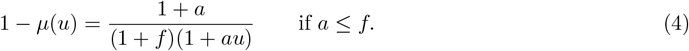

The leftmost plot in Figure 1 visualizes equations (3) and (4); the others are discussed in Section 7. Mortality regulation is also demonstrated in Box 1, and most of the examples in this paper use this model with *a* = *f* . In practice, an empirical estimate of the survival probability at low density (the limit as *u* → 0) *s*_0_ can be incorporated so that *a* = *s*_0_(1 + *f*) −1.

**Figure 1:**
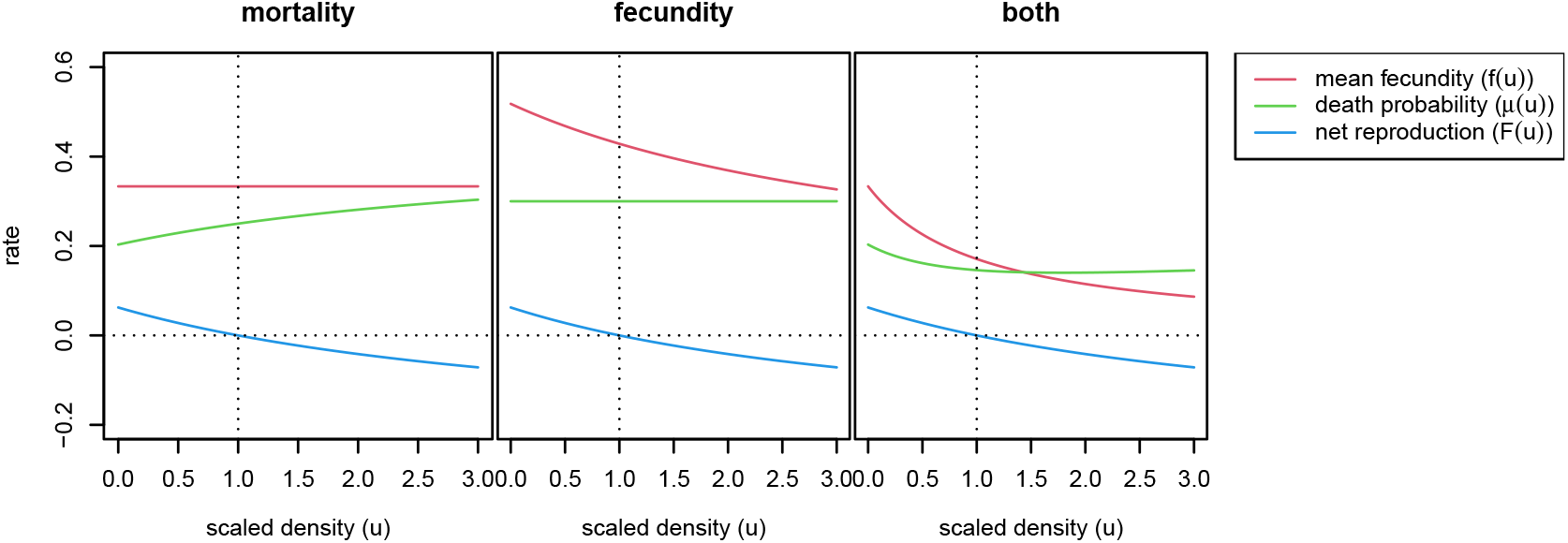
Three example models with “Beverton–Holt” regulation of population density. The relationship between scaled local population density and vital rates: mean fecundity *f* (*u*), probability of death *µ*(*u*), and net reproduction rate *F* (*u*) = *f* (*u*) − *µ*(*u*). In each, the horizontal axis is shown in units of *K*, the parameter controlling equilibrium density (“carrying capacity”). The models are: **mortality** from equation (4), **fecundity** from equation (5), and **both** from equation (6).

The condition that *a* ≤ *f* is so that this gives us a valid probability: otherwise, the survival probability can be negative. Note that the strategy used here cannot work with the discrete logistic: since *F* (*u*) = 1− *u* can get arbitrarily negative, eventually random density fluctuations will lead to negative probabilities, and errors in the simulation.

There are many other possible ways to set up *f* (*u*) and *µ*(*u*) that result in the same net form of density dependence. (See Section 7.1 and Appendix C for how stochasticity alters the realized equilibrium density and alternative functional forms.) The general forms of *f* (*u*) and *µ*(*u*) may also need to be modified depending on the organism. If there are two sexes and only one sex can bear offspring, then *f* should not be the total number of offspring, but rather the number of offspring *of the offspring-bearing sex*. More generally, if vital rates depend on any aspect of the individual (for example, only adults can reproduce, or mortality is age-dependent), then the equivalent calculations must be done with a matrix population model (Caswell, 2000) or an integral projection model (Ellner et al., 2016). Our examples mostly ignore such complications, although several models have distinct life stages (Case Studies 9.2 and 9.4).

#### Box 1

**Regulating mortality and fecundity in SLiM**

The Wright–Fisher model is a population model with a *fixed* population size. A Wright–Fisher model in continous space, therefore, has *global* population regulation – individuals are affected by others arbitrarily far away, resulting in counter-intuitive, unrealistic consequences (Felsenstein, 1975). SLiM’s default model is the Wright–Fisher model (“WF”). In this study, our goal is to model particular species in realistic ways, so we use only SLiM’s non-Wright–Fisher (“nonWF”) model, which requires explicitly choosing how births and deaths occur.

In a SLiM nonWF model, the “fitness” attribute of an individual is the probability of survival until the next time step. So, controlling mortality in SLiM is simply a matter of setting individuals’ fitness. To compute local density, we use the `localPopulationDensity()` function, which computes density just as described in equation (1) (with options for the choice of kernel). To do this we need to first (during setup) define an `InteractionType` object, here using a Normal kernel with a scale of `SX` (as in, *σ*_*X*_) and a maximum distance of `SX * 3`):

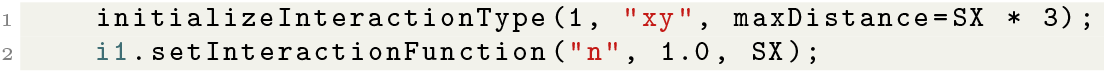

Now, to use Beverton–Holt regulation on mortality as described around equation (4) (setting *a* = 1 in equation (4) and with *K* defined elsewhere), in each time step, we use this interaction type to set survival probabilities:

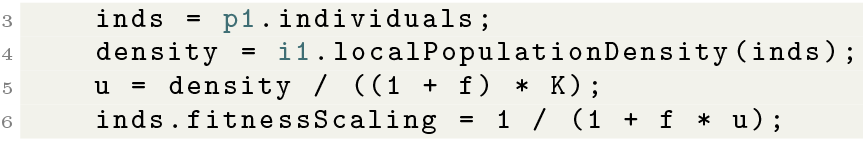

Note that the local population density is measured after reproduction and before death (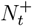in Appendix B), so we divide by 1 + *f* as well as *K* when converting it to scaled population density *u* from the equation (1), so it reflects density *before* reproduction.

On the other hand, offspring are produced by a “`reproduction()` callback,” a chunk of code that is executed each time step for each individual, and produces any desired new offspring for the focal individual. For instance, if we would like to use Ricker regulation on fecundity (see equation (19) with *α* = 1 and *β* = 0), we might for efficiency pre-compute each individual’s number of offspring:

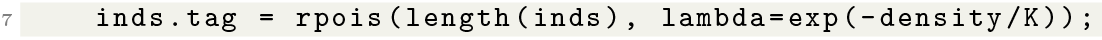

and then in the `reproduction()` callback produce the offspring:

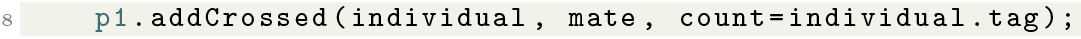

## 3 Spatial scales and neighborhood sizes

Now that we understand how to obtain stable simulations, we can move on to a more spatial topic: that of spatial scale. It is convenient to use spatial scale parameters to describe distance within the model; some are depicted in Figure 2: (*i*) interaction scale *σ*_*X*_, the typical distance over which individuals affect each other ecologically, (*ii*) dispersal scale *σ*_*D*_, the typical distance between parent and offspring, (*iii*) mate choice scale, *σ*_*M*_, the typical distance between mates, and (*iv*) movement scale *σ*_*V*_, the typical displacement of an individual each time step. (*σ*_*D*_ is often called “dispersal distance,” but here we use “dispersal scale” for consistency.) We met *σ*_*X*_ in equation (1), where it determined which individuals were close enough to each other to affect local density.

**Figure 2:**
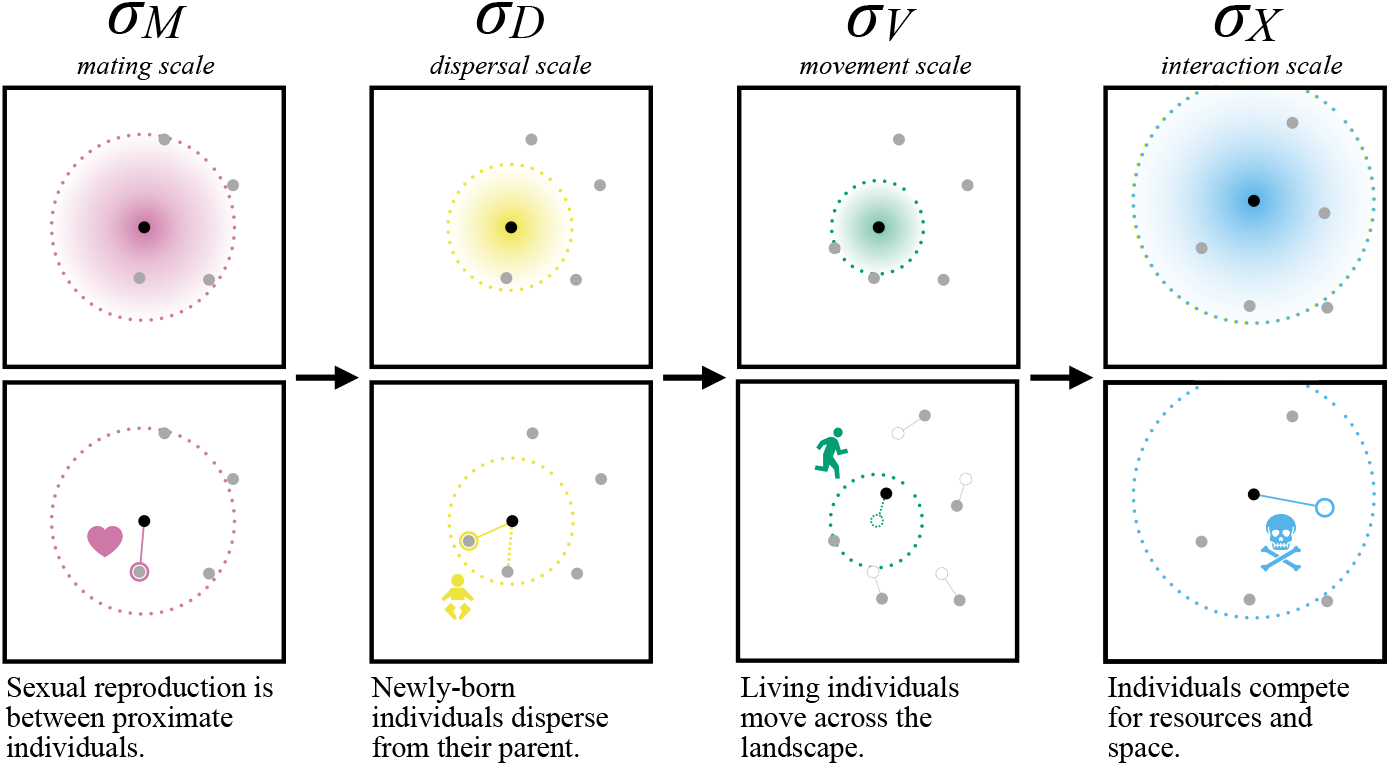
Spatial processes act over spatial scales defined by the *σ* parameters (radii of dotted circles). In one simulation time step, mate choice, offspring dispersal, adult movement and inter-individual interaction (columns) all take place. (The order of these events is shown here as performed in SLiM.) The scale over which these processes occur are parameterized by *σ*_*M*_, *σ*_*D*_, *σ*_*V*_, and *σ*_*X*_, respectively; they can be used to calculate neighborhood sizes, to define the scale of the kernels used to draw movement or dispersal vectors (see Section 4) and to estimate local population density (see Section 5).

Each spatial scale parameter determines how many other individuals typically exist in an individual’s “neighborhood.” First, the *mating neighborhood size* 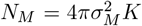 and the *interaction neighborhood size* 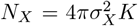,measure, respectively, the typical number of other individuals in a circle of radius 2*σ*_*M*_ or 2*σ*_*X*_ at equilibrium.

In practice, these measures are only intended to be order-of-magnitude diagnostics. For example, *N*_*M*_ should be the same order of magnitude as the number of potential mates; if it is small then mate limitation may be a problem. That said, depending on the mating kernel (*i.e*., shape of the distribution that determines how “attractive” nearby mates are), individuals as far away as 3*σ*_*M*_ are probably also available to mate. (See Section 5.)

Less obvious but equally important is *N*_*X*_, which measures the typical number of other individuals that “count toward” the local density of a given individual, and hence affect its demographic rates (such as survival probability, as in Box 1). *N*_*X*_ also provides a measure of individual-to-individual variability in local density. Roughly speaking, a larger *N*_*X*_ means that local density is obtained by averaging over an area with more individuals, meaning that individuals across the landscape experience more similar local densities. When *N*_*X*_ is small, densities across the landscape can be more noisy – with some individuals experiencing no competition from neighbors, and some experiencing high density. Further discussion of how *N*_*X*_ and *N*_*M*_ can help diagnose odd model behavior is given in Appendices B.2 and B.3.

Another “neighborhood size” is a classical one: Wright’s neighborhood size 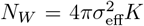,where 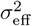 is the (squared) *effective dispersal distance*, the variance of the displacement between parent and offspring along any axis looking back along a lineage (chain of parent–offspring relationships). This quantity appears frequently in work on continuous spatial models in population genetics (e.g., Wright, 1943, 1946; Barton et al., 2002; Rousset, 1997; Robledo-Arnuncio and Rousset, 2010), and is clearly affected by *σ*_*D*_, *σ*_*V*_, and *σ*_*M*_, as well as the mean generation time if adults move, but no explicit expression for *σ*_eff_ from these parameters is known. *N*_*W*_ gives, roughly, the number of “potential parents” of a given individual, and so is a measure of the rate of local genetic drift. If *N*_*W*_ is small, then local inbreeding (and spatial structure more generally) will be stronger.

Although we encourage basing modeling decisions on empirical understanding, it is not always feasible. For instance, suppose one is simulating a species of fairly common shrub that lives widely across a landscape.

In practice, its local density is determined by microhabitat and complex interactions with other species. Any concrete estimate of the interaction scale for a plant is probably quite small – the competitive effect of one shrub on another more than a few meters away is (in the short term at least) usually quite small. However, implementing a spatial model with an interaction scale of only a few meters (and no other species) will likely lead to a population size that is much too large. One option is to somehow include other species and fine-scale habitat suitability, but doing this in a realistic and efficient way can be a major challenge. A simpler option is to set the interaction scale to be on the order of the mean inter-individual spacing, and adjust the form of density dependence to roughly match the observed population density. The simulated population will probably be more evenly spread out across space than in reality, but it is hopefully at least a better approximation than a nonspatial model. More work is needed to develop appropriate modeling strategies for such situations, and to understand their consequences.

### Box 2

**Summarizing the state of the population**

SLiM’s GUI lets the user visualize the state of the simulation as it unfolds. We can customize the display to see areas of higher fecundity, differences in age structure, or even local adaptation.

One strategy is to set the color of each individual: by default, individuals are colored by survival probability, but this code snippet will set the color to reflect the sex of each individual:

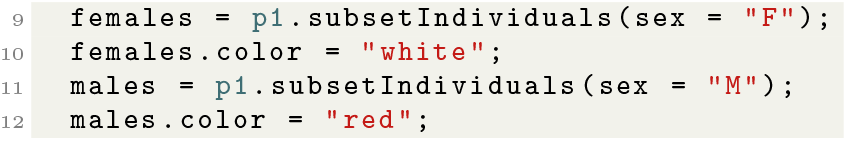

Another strategy is to summarize the state of the population as a map. This can be done with the `summarizeIndividuals()` function, which creates a rasterized map for which the value of each pixel is some summary of the individuals within that pixel.

For instance, the map of density shown as the background was made using the following code:

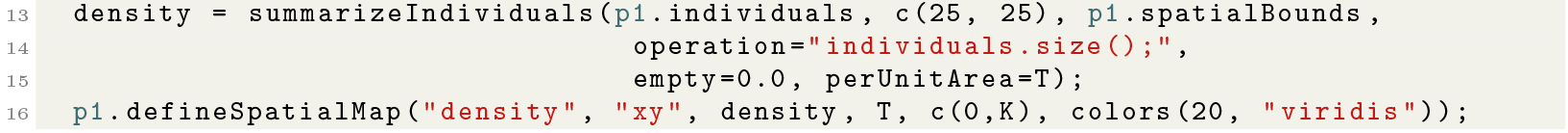

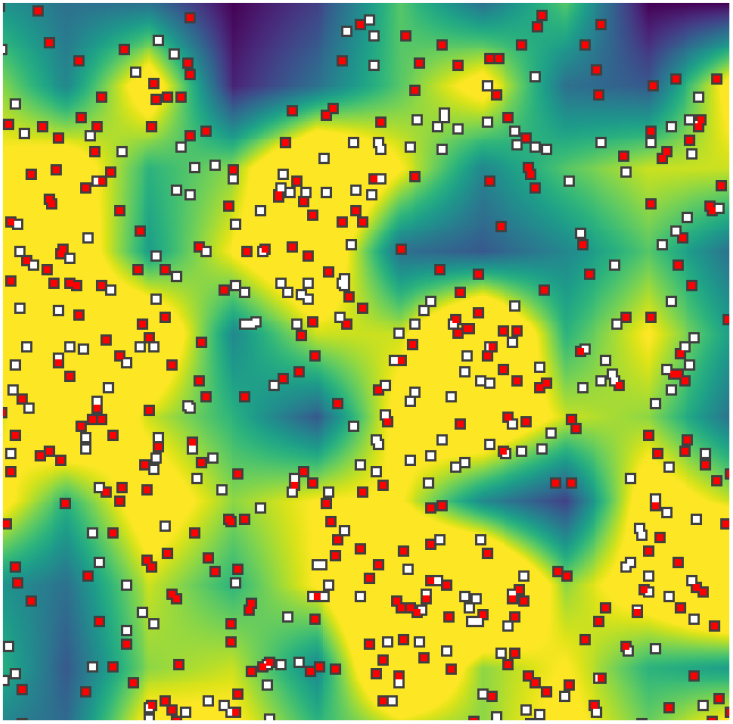

In this close-up screenshot of a dioecious simulation, there tend to be more females (white points) where the local density is higher (yellow background); and there tend to be more males (red points) where the density is lower (blue background). We discuss the heterogeneous spatial patterns generated in dioecious simulations in Section 5. See Box 5 for more use of `defineSpatialMap()`.

## 4 Movement and dispersal

How individuals move across the landscape influences both their spatial distribution and how related they will be to nearby individuals. We consider parent–offspring dispersal and the movement of organisms during their lifetimes as different processes, but they can be implemented in similar ways. To model dispersal and movement, we need to consider both the overall scales of movement (*σ*_*D*_ and *σ*_*V*_ ; introduced in Section 3), as well as the shape of the dispersal distribution, or *kernel*.

The easiest way to implement dispersal in two dimensions is simply to say that an individual at location ***x*** = (*x, y*) will produce an offspring that lives at (*x* + *σ*_*D*_*X, y* + *σ*_*D*_*Y*), where *X* and *Y* are independent draws from a Gaussian. Most simulations in this paper were done in this way, for familiarity rather than any particular reason.

### Box 3

**Random dispersal and displacement**

Perturbing a spatial location by adding a displacement drawn from a given kernel is a common operation in simulations – for instance, to choose offspring locations. To do this, SLiM provides the `pointDeviated()` method, which also needs to know the shape of the kernel, the type of boundary condition, and other parameters. For instance, the following code:

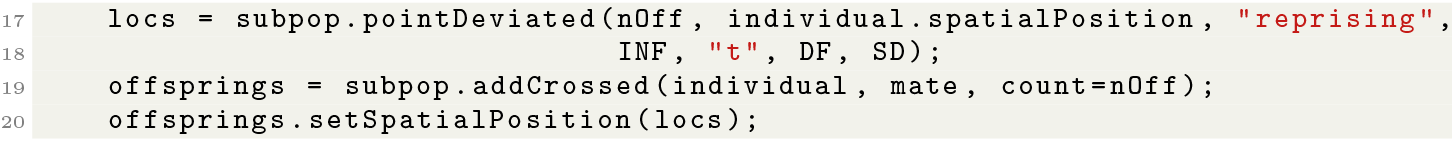

creates `nOff` offspring, and sets each offspring’s position to a randomly sampled location near the location of the parent (`individual`). The random displacement is drawn with density *f* (*x*) ∝ (1 + *x/*SD ^2^*/*DF)^−(DF+1)*/*2^, *i.e*., from the kernel whose density is formed by rotating the *t* distribution with scale `SD` (as in *σ*_*D*_) and `DF` degrees of freedom about the origin. (See Appendix D for other strategies.) The `reprising` argument conditions the result on falling within the spatial bounds of the simulation – other options include `stopping` and `reflecting`. (Or for an absorbing boundary, `none`, with offspring falling outside the area removed.)

Although the scale of movement most strongly affects spatial patterns, the shape of the kernel is also important, and can have surprising effects: even very rare long-range movement can have strong effects on the speed of a range expansion (Mollison, 1972; Paulose et al., 2019) or the relationship between genetic and geographic distances (Smith and Weissman, 2023). To see the effects of rare, long-range movement, a convenient “fat-tailed” kernel is the Student’s *t*: the smaller the degrees of freedom parameter, the more likely are extremely long movements.

How to draw from a different dispersal kernel? The first guess – choose *X* and *Y* from a different distribution – does *not* work: the result will not be rotationally symmetric, and dispersal will tend to fall along the *x* or *y* directions. Options are to either move a random distance at a uniformly chosen angle (which is conceptually simpler), or to multiply a bivariate Gaussian by a random scaling factor (which has other advantages). What to call a given two-dimensional kernel is not standard – for instance, would a “Student’s *t* kernel” have a *t*-distributed distance? Or, a *t*-shaped cross-section? In simulations below we use the latter convention, as described in Box 3, and discuss these choices more in Appendix D.

The behavior of individuals near boundaries must of course be specified. Common choices include “reflecting” and “absorbing”. These differ substantially: imagine the new individual as taking a straight-line path from (*x, y*) in the direction of (*X, Y*) and encountering a boundary either bounce off (reflecting) or die (absorbing). The latter clearly reduces the effective fecundity of individuals near the boundary, and so reduces mean density up to a few multiples of *σ*_*D*_ away. Another choice is “reprising”, for which the random draw (*X, Y*) is chosen conditionally so that (*x* + *σX, y* + *σY*) stays within the range. (For more details of the consequences of boundary conditions, see Mazzucco et al. (2018).)

Movement in practice often depends on the environment, of course: organisms tend to move within particular habitats, and barriers are ubiquitous on all scales. Small-scale heterogeneity may be averaged out across the time scale simulated, and so incorporated (implicitly) in the movement kernel. However, large-scale heterogeneity can be important. Movement on a heterogeneous landscape still relies on some way of randomly choosing nearby points, and hence a movement kernel. One way to incorporate this is discussed in Box 5.

### 4.1 Clumping

In many species reproduction is local, and so tends to produce clumps which are spread out by movement and dispersal. Such patterns can be intriguing or puzzling, and have real consequences for demography and genetic variation. Spatial clumpiness is sometimes visually obvious (as in Figure 3B), but more generally the clumping tendency of individuals can be measured by spatial correlations.

**Figure 3:**
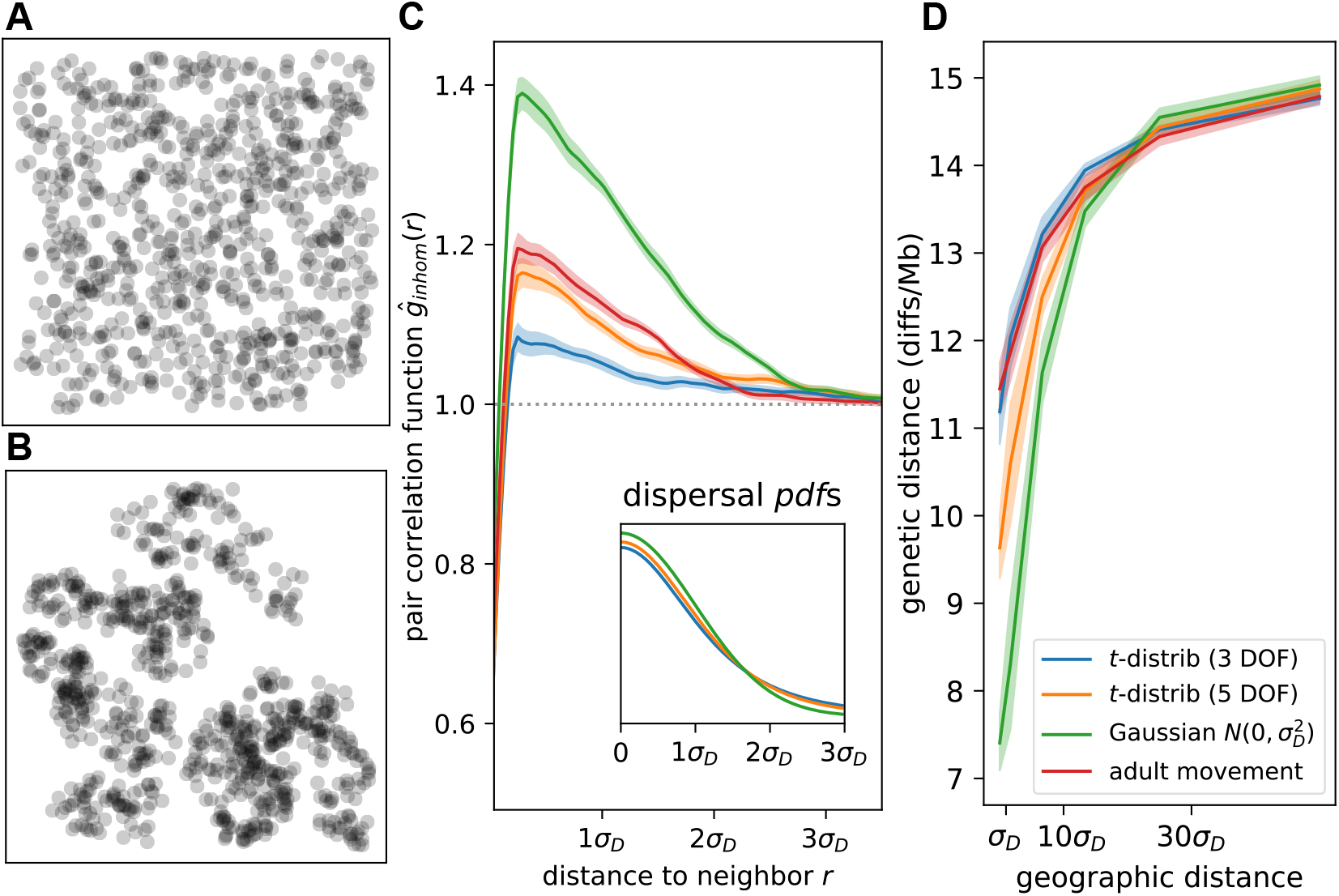
**(A, B)** Examples of simulations showing clumping (B) and not (A). **(C)** Dispersal kernel and movement type affects magnitude and spatial scale of clumping. Pair correlation functions (main panel) show density of pairs of individuals found a particular distance apart, relative to distances expected under a Poisson process (1.0; grey dotted line). Curves show the average across 50 independent time steps. Probability density functions (inset) for dispersal scales under the three dispersal kernels used: *t*-distribution with 3 degrees of freedom, *t*-distribution with 5 degrees of freedom, and Gaussian. Scale of *t*-distributions is equal to the standard deviation of the normal distribution (*σ*_*D*_ = 0.3). “Adult movement” scenario uses the Gaussian for dispersal as well as for movement at each time step. **(D)** Fat-tailed dispersal and adult movement flatten genetic isolation-by-distance. Plots show mean genetic distance between pairs of individuals at increasing geographic distance, averaged across ten independent replicates. Figure S3 visualizes clumping quantified in (C, D); Figure S4 quantifies clumping visualized in (A, B).

One informative measure of clumping is the *pair correlation function*, which shows for each distance *x* how likely an individual is to have a neighbor at that distance relative to the average density. (See Baddeley et al. (2015) other useful descriptors of point data.) Concretely, the pair correlation function estimates the mean density at distance *x* away from an individual divided by the overall mean density, averaged across individuals; if the points are independently placed, it is constant at 1.0. Figure 3C shows pair correlation functions for simulations in which adults do not move and dispersal follows either a *t* or a Gaussian distribution, or in which both dispersal and adult movement are Gaussian (with *σ*_*D*_ = *σ*_*V*_).

Individuals are more likely to be around a distance of *σ*_*D*_ from each other than otherwise, but this tendency is reduced with more long-distance dispersal (*t* dispersal with lower degrees of freedom; Figure 3C). In other words, a little long-range dispersal reduces clumping. Note, however, that in these examples the scale of clumping is quite narrow: correlations only extend out to 2 or 3 multiples of *σ*_*D*_. For another visualization of this relatively subtle clumping, see Figure S3. Unsurprisingly, adding adult movement (*σ*_*V*_ ≠ 0) to a dispersal-only model reduces correlations as well.

In spatial models, neighbors tend to be more related to each other than to distant individuals, a pattern known as “isolation by distance” (Wright, 1943). The scale at which this correlation appears is determined by how far individuals move, and is also affected by the shape of the dispersal kernel: Figures 3C and D show that simulations with more long-range movement – but with comparable mean dispersal scale – tend to have a weaker relationship between geographic and genetic distance. (See also Smith and Weissman (2023).)

## 5 Mating and other pairwise interactions

Mating is a crucial interaction for biological simulations, and there are numerous aspects and choices to consider. We probably don’t want to simulate the detailed movement of individual pollen grains or the meanderings of a male moth seeking a female, and instead would like to skip to the realized outcome, *i.e*., “choose a mate nearby.” (We follow the literature on mating systems in calling this “mating,” even when referring to plants or broadcast spawners.)

As with dispersal (Section 4), it is easiest to specify what “nearby” means with a kernel: roughly, an individual can be chosen with probability proportional to the kernel. More concretely, if the kernel is *ρ* and the mating scale is *σ*_*M*_, then an individual at ***x*** would assign weight *w*_*i*_ = *ρ*((***x***_*i*_ − ***x***)*/σ*_*M*_) to another individual at location ***x***_*i*_. The probability individual *i* is chosen is equal to *w*_*i*_ divided by the sum of weights across all nearby individuals. This same method can be used for other individual-to-individual interactions, such as predators choosing prey individuals.

We can adjust the typical distance between mates with the mating scale, *σ*_*M*_, which is directly analogous to parameters used to describe interaction (*σ*_*X*_) and dispersal (*σ*_*D*_) scales (Section 3), although each uses an underlying kernel in slightly different ways. Of course, the criteria determining potential mates for a given individual differ widely among species (Shuster, 2003). Relevant questions about the mating system include: How often does selfing occur, and under what circumstances? Are sexes separate (dioecy/gonochory) or not (monoecy/hermaphrodity)? Are there distinct mating types or self-incompatibility systems? An important note is that for dioecious species, calculations to determine stability of population density are easiest if done using only the reproducing sex.

### Box 4

**Interactions, and mating**

The mechanism that SLiM uses to mediate most effects that some individuals have on others is called an “interaction type” (see Box 1). We used a symmetric interaction type in Box 1 to compute local density: every individual affected every other. Some interactions are not symmetric: we might, for instance, want each female to be able to find nearby males. To do this, we first set up a sex-specific interaction (again using a Gaussian kernel, with standard deviation `SM` as in *sigma*_*M*_):

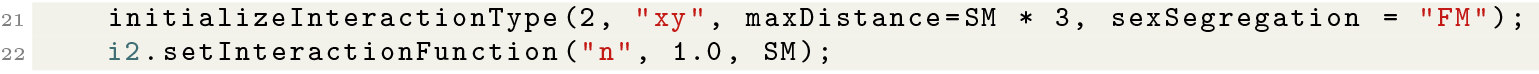

The `sexSegregation` parameter value of `“FM”` means that females will receive the interaction and males will exert the interaction; it is asymmetric. Then, we can use the interaction in a `reproduction(NULL, “F”)` block as follows (the `NULL` and `‘‘F’’` arguments imply that it applies to all females) to produce a single offspring:

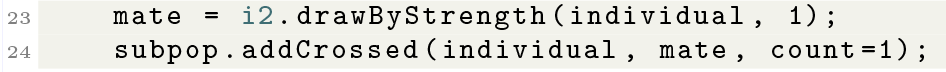

This chooses a male `mate` from among the neighbors of the focal female, `individual`, with probability proportional to a Gaussian kernel with standard deviation `SM`. The interaction type itself guarantees that the chosen mate will be male. It is possible to set up other constraints on interaction types as well, such as a minimum or maximum ages, to represent other constraints on the reproductive eligibility of individuals.

### 5.1 Sex-specific spatial structure

The combination of density dependence and mating system can have surprising consequences: for instance, in dioecious simulations, clustering is dependent on sex. To illustrate this, Figure 4 shows the *mark connection function* (Baddeley et al., 2015, § 14.6.4.2) for female/male, female/female, and male/male neighbors, which shows the proportion of pairs of points at distance *r* that are one female and one male, two females, or two males, respectively. Curiously, we see that the probability that an individual’s neighbor is of the other sex does not depend on that neighbor’s distance. However, the probability of having a *same*-sex neighbor *does* change with distance. As shown in Figure 4, within 3*σ*_*D*_ neighbors of a female are more likely than expected to be female, and nearby neighbors of a male are less likely to be male than expected.

**Figure 4:**
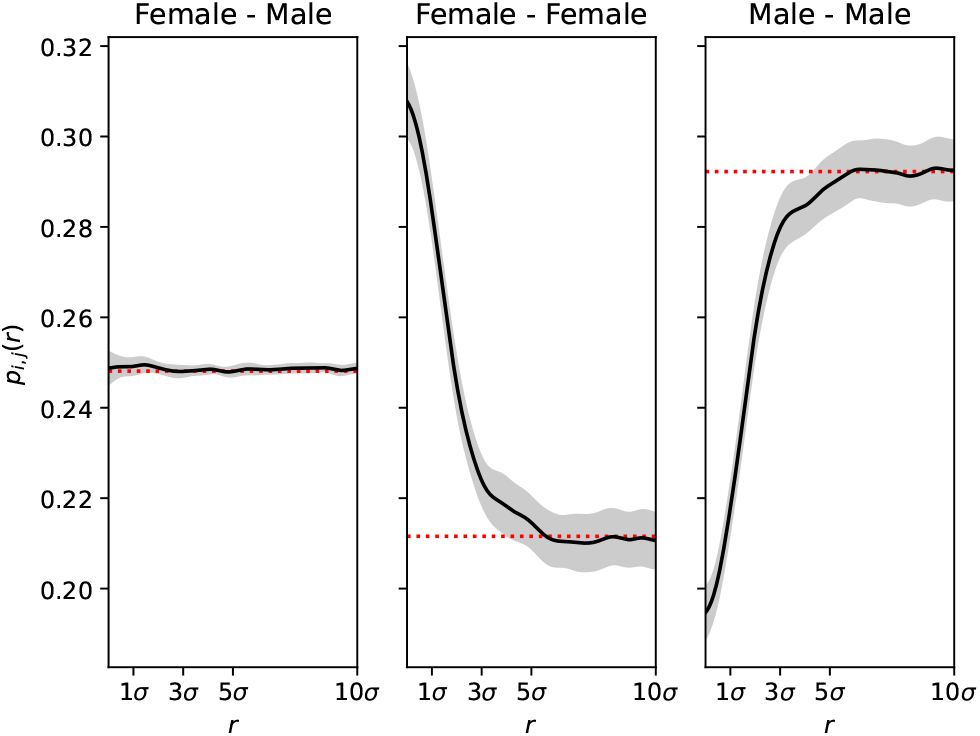
Dioecy generates female underdispersion, male overdispersion. Each panel shows the proportion of pairs of individuals at the distance shown on the horizontal axis that are either **(left)** a female and a male **(center)** both female, or **(right)** both male. Average and standard deviation of 50 independent simulation ticks are shown. Red dashed lines indicate expectation if individuals’ locations were chosen uniformly. Note that the “Male–Female” proportion is be identical to the “Female–Male” proportion, so that twice the left panel plus the other two panels is equal to 1.

Why does this happen? A simple reason is that we are modeling a dioecious scenario with no adult movement and where offspring are only generated by females and placed nearby. Local density-dependent mortality means that all individuals tend to kill their neighbors, but only females can replace them. Correspondingly, the spatial range of a dioecious system where offspring disperse from a particular sex is determined by the range of that sex – others on the periphery cannot extend the range because offspring do not disperse from them. This is a disadvantage for colonizing new areas (Obbard et al., 2006), and may explain the spatial distribution of dioecious individuals away from a range front (Mirski et al., 2017).

Indeed, Shuster (2003) showed that female aggregation (as observed in Figure 4) is a universal consequence in mating systems with female choice (Shuster, 2003, ch. 2). Other aspects of the simulation may differ by sex: for instance, sex-biased dispersal is pervasive (Trochet et al., 2016) and generates detectable spatial patterns of genetic relatedness between sexes (Aguillon et al., 2017; Broquet and Petit, 2009; Laporte and Charlesworth, 2002). Social mating structures can induce sex-biased dispersal and thereby create similar patterns (Pusey and Packer, 1987; Hammond et al., 2005).

## 6 Maps: spatial heterogeneity

### Box 5

**Defining and manipulating maps**

SLiM provides support for defining and manipulating spatial maps. We can read the values for a spatial map (say, of elevation) from a `.csv` file containing a rectangular grid of values using the code below:

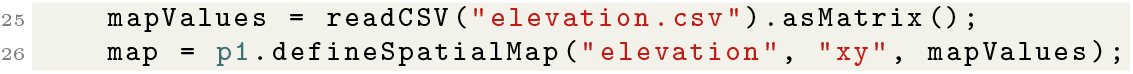

(Images can also be read in as `.png` files.) If this is a low-resolution raster then we may wish to smoothly interpolate it to higher resolution, done below with bicubic interpolation:

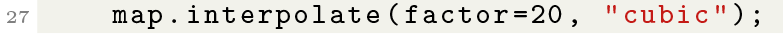

Once we have this map, we can extract the values of the map at arbitrary locations – for instance, the elevations at which individuals live:

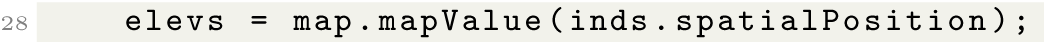

Further operations are available, including blurring and algebraic manipulations of the values. The dispersal method in Box 3 allowed individuals to move equally well in any direction. To guide movement with a map – for instance, to induce a preference for moving uphill, for this example where map values indicate elevation – we can use the `map.sampleNearbyPoint()` method:

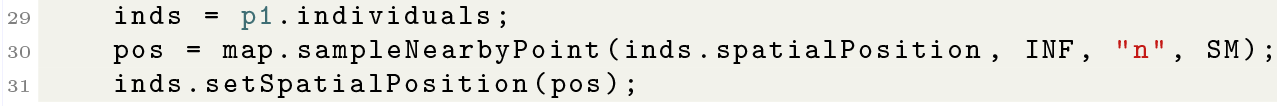

This will move each individual to a new location sampled nearby and weighted by map value: if the original location is ***x***, *ρ*() is a Gaussian kernel (specified as type `“n”`) of width `SD`, as in *σ*_*D*_, and the value of the map at location *y* is *m*(*y*), then the new location is chosen with density proportional to *m*(***y***)*ρ*((***y*** − ***x***)*/σ*_*D*_).

Most real populations are far from uniformly distributed in space, and in most cases, the underlying cause is thought to be environmental heterogeneity. Until this point, we have considered simulations of homogeneous landscapes only. Such “flat” landscapes are useful for developing models and/or theory, but incorporating aspects of real landscapes can make simulations more realistic. Spatial heterogeneity can be introduced simply by making some parameter of the model, such as fecundity or mortality (*f* or *µ* as defined in Section 2), vary across space – in which case we can visualize the parameter as a map. Such maps might represent specific environmental conditions, habitat boundaries, or abstract habitat quality.

Raster-based images provide a convenient way to introduce a map of spatial heterogeneity into a simulation framework that cannot directly read geospatial data formats. A monochrome `.png` file consists of a rectangular grid of pixels with integer values between 0 and 255. These values can be shifted and scaled to lie within a useful range. See Case Studies (Section 9) for several examples of the use of images, and Box 5 for an example using `.csv` data. High-resolution maps to use as source images are publicly available from various sources, including NASA’s Earthdata platform (2024), ESA’s Earth System Data Lab (2024), or PRISM Climate Group. Open-source tools for processing remote sensing data are also available (see Montero et al., 2023).

To use a raster-based image, we must decide how the bounds of the image map onto geographic space. In general, we match the (rectangular) image with the (rectangular) spatial area to be simulated. The image is represented as discrete pixels, but we need to obtain values of the map at arbitrary locations (not just at the center each raster pixel). This can be done by either associating each pixel’s value with the rectangle it would visually cover (as seen in Box 2), or associating each pixel’s value with the corresponding cell’s midpoint and interpolating between.

In the simple example shown in Figure 5, an image represents the altitude map of a mountain, with darker red shades indicating higher elevation. The local carrying capacity, *K*(***x***), is modified by the value of the map at ***x*** so that high-elevation locations can support more organisms. Comparing the provided altitude map (background of Figure 5A) with a summary of realized average density (Figure 5B), demonstrates that population density roughly matches the underlying elevation map.

**Figure 5:**
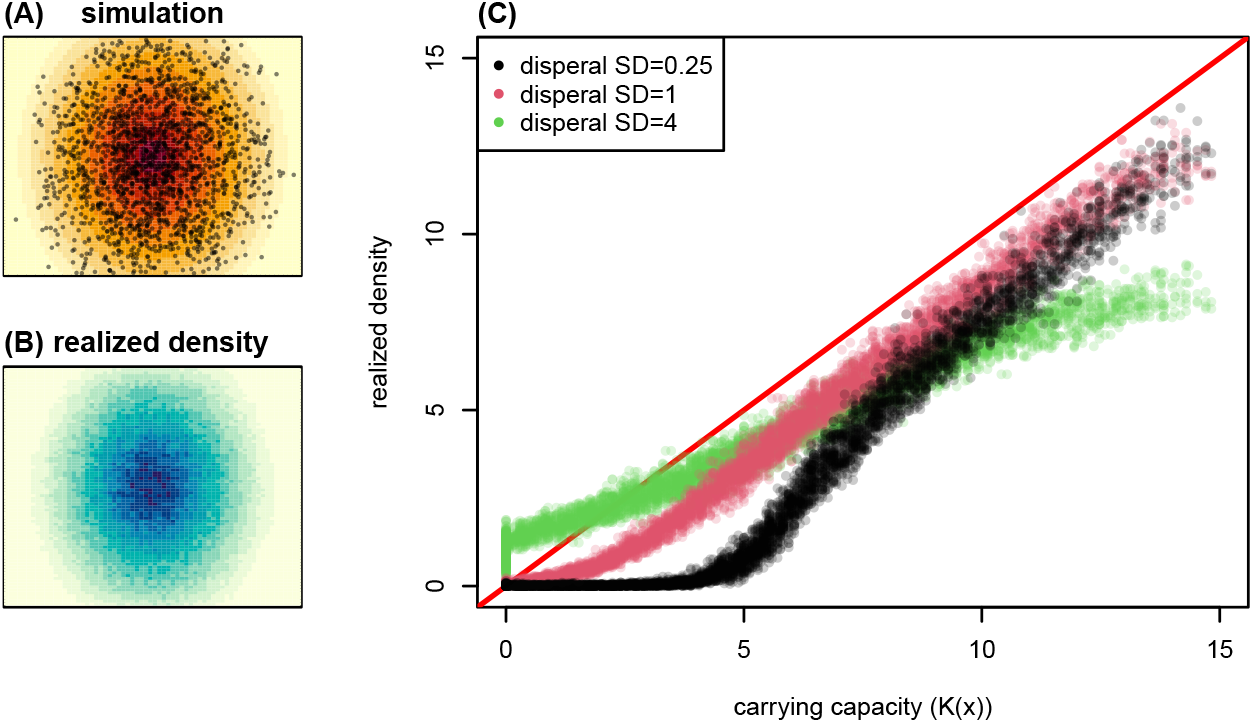
A spatial simulation using a heterogeneous map of carrying capacity generated from an image file. **(A)** A representative time step during the simulation, with each of the roughly 1500 individuals shown as points and the map of carrying capacity, *K*(***x***), shown in the background. Red hue indicates greater *K*(***x***) in the pixel. **(B)** A map of realized density, averaged over 10^4^ time steps. Blue hue indicates greater individual density in the pixel. **(C)** Comparison of realized density to carrying capacity from three different runs using different dispersal scales: each point shows the realized density in one of the pixels of the map shown in (B) plotted against the value of *K*(***x***) in the center of the pixel, for three values of *σ*_*D*_ (labeled “SD” in the legend). The dimensions of the map are 25 *×* 20, and the interaction scale is *σ*_*X*_ = 0.3. The simulation has Ricker regulation of fecundity: the expected number of offspring of an individual at location ***x*** is 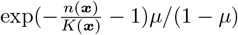,where *µ* is the (fixed) probability of survival, *n*(***x***) is the local density at ***x***, and *K*(***x***) is obtained from the value of the image at ***x***.

**Figure 6:**
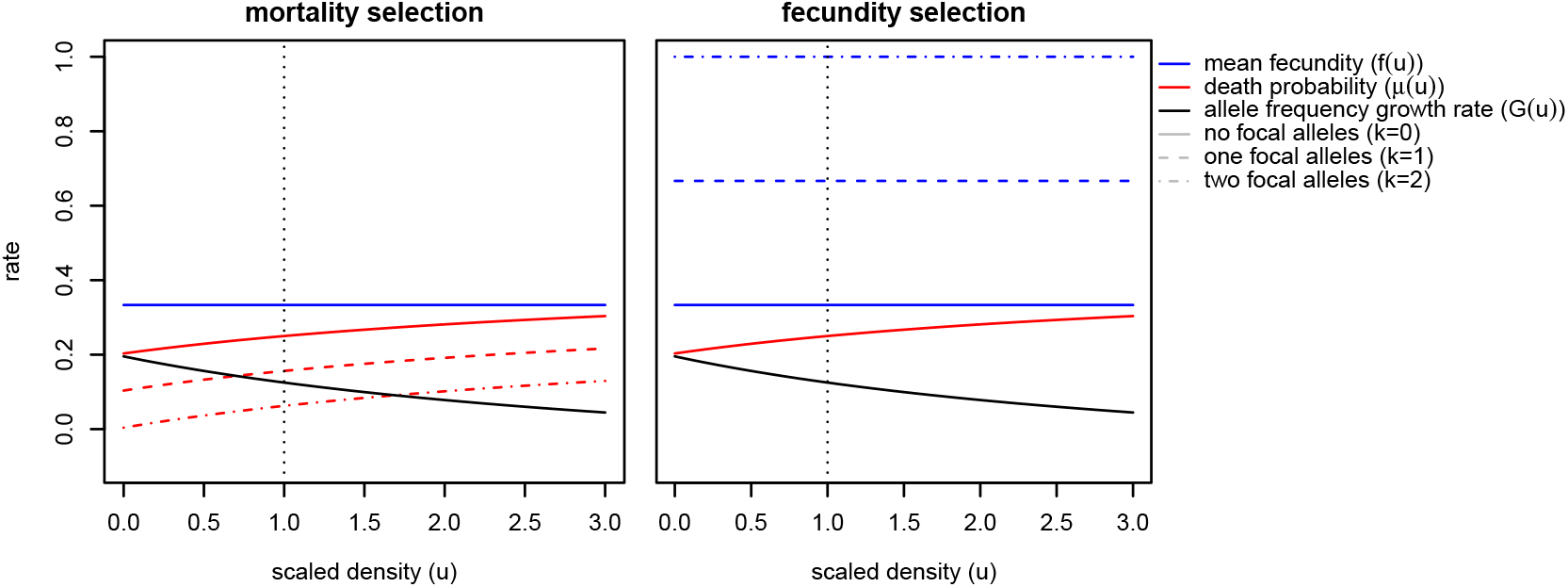
Density dependence of for two methods of selection: on **(mortality)** or **(fecundity)**, as described in equations (8) and (10) with *s* = 0.25. Line types show the mortality (red) or fecundity (blue) for individuals carrying *k* = 0, 1, or 2 copies of the beneficial allele. The mean growth rate of the beneficial allele when rare (black line; *G*(*u*) from equation (9)) is the same for both types of selection, at all densities. The vertical dotted line is at scaled density *u* = 1. Other parameters are as in Figure 1.

A more precise comparison (Figure 5C) shows that *n*(***x***), the local density at ***x*** defined in equation (1), is not exactly *K*(***x***). In most situations, the equilibrium density is below *K*(***x***). This is probably for two reasons: first, stochasticity usually reduces equilibrium density (see Section 7.1 and Appendix B.4); second, since the mountain is conical, most locations ***x*** are surrounded by more low elevation area (where *K* is lower than *K*(***x***)) than higher area (where *K* is higher than *K*(***x***)), making the net flux of migrants at ***x*** negative. The degree of deviation of *K*(***x***) from *n*(***x***) can depend on other factors such as *σ*_*D*_. When dispersal is short (*σ*_*D*_ = 0.25, black points), areas with carrying capacity below about 5 individuals per unit area are not self-sustaining. (See Appendix B.3 for more discussion.) On the other hand, with long-range dispersal (*σ*_*D*_ = 4, green points), the overall relationship between density and carrying capacity is flattened as offspring from high-fecundity areas end up across the entire range. However, most offspring are still produced near the top of the mountain, and lower elevations are maintained by source-sink dynamics.

## 7 Density dependence, life history, and stochasticity

In our initial model we used Beverton–Holt density-dependent feedback on mortality to control global population size (Section 2.2). Here we give examples that have the same Beverton–Holt form for the net effect of density on population regulation, but differ in other ways.

### Fecundity regulation

The Beverton–Holt density-dependent regulation in equation (4) has the probability of death increase with local density while fecundity stays constant. Alternatively, we can set the probability of death to a constant: *µ*(*u*) = *µ*_0_, and then solve for *f* (*u*) to obtain the Beverton–Holt form *F* (*u*) = *α*((1 + *a*)*/*(1 + *au*) − 1). (It turns out we will need the scaling factor 0 *< α <* 1 to make a model with positive birth rates and death probabilities between 0 and 1.) Then, fecundity should depend on scaled density *u* as follows:

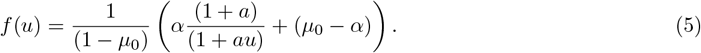

#### Box 6

**Adaptive empirical tuning for emergent parameters**

We often want a simulation to match a given estimated or observed density. However, this is not as simple as setting the value of *K* (local carrying capacity, introduced in Section 2) in the code, because population density is an *emergent* quantity – a complex consequence of births, deaths, movement, and local interactions. Fortunately, matching even emergent quantities is possible in SLiM. The code in this box dynamically adjusts parameters to make population density match a desired value. The same general technique may be applied to other quantities such as mean age or degree of clustering.

We first define a global parameter `ADJ` that will be adjusted:

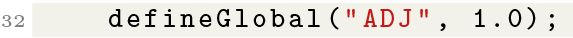

Then we modify the density regulation code from Box 1 so that `ADJ` adjusts the carrying capacity:

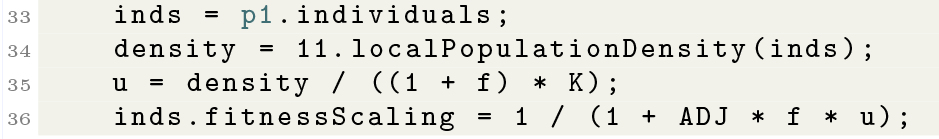

In each time step, `ADJ` is updated by a factor exp[*α*(*Y* − *K*)], where *Y* is global population density (population size divided by total area) and *α* is an update rate (*e.g*., *α* = 0.01):

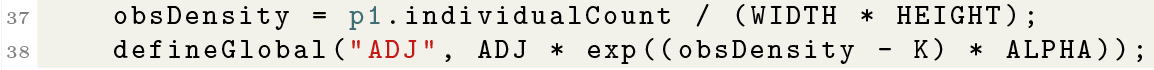

When *Y > K*, we want the density to be lowered in the next cycle, so `ADJ` is increased, decreasing carrying capacity. On the other hand, when *Y < K*, `ADJ` is decreased so that we get a higher realized global population density *Y* in the next cycle.

Importantly, this tuning should stop after an initial “burn-in” period, and the appropriate value for `ADJ`, once found, should be hard-coded into the final model. If changes occur in the simulation that affect population size (*e.g*., a reduction in habitat), this adaptive code will force the density back to the chosen value, which is generally not desirable – the population size should, in fact, change in response to such changes in the simulated conditions.

This functional form is shown in the middle panel of Figure 1.

### Compensatory regulation on juvenile and adult mortality

To stabilize the population, the effect of density on *net* reproductive rate must be negative, but the effects on individual demographic components (such as birth or death) can be positive, as long as they are compensated for by other components. For instance, the probability of survival 1 − *µ*(*u*) can *increase* with density as long as fecundity *f* (*u*) decreases faster. One way to do this is to set

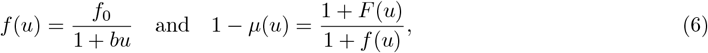

where *F* (*u*) = *α*((1 + *a*)*/*(1 + *au*) − 1); *i.e*., the Beverton–Holt form again. This produces valid survival probabilities if *f*_0_ *≥ aα*. Depending on the choice of the parameters, *f* (*u*) may increase or decrease with *u*, and is not necessarily monotone. One choice of valid parameters is shown in the rightmost panel of Figure 1. Our three examples (4), (5), and (6) (illustrated in Figure 1) might all be reasonably called “Beverton– Holt” models, although they differ substantially in the underlying mechanism of regulation. Although they have similar behavior around equilibrium density, they have quite different life history implications. Most strikingly, in the first model, mortality increases with density, in the second mortality is constant, while in the third, mortality decreases with density. There are corresponding differences in age structure among the models (as shown in Figure S5), although the dynamics of total population size are similar. More examples along these lines are given in Appendix C.

### 7.1 Stochastic effects

Despite all this theory, in practice, equilibrium density is usually not *K* – it is often lower. (See Figures 5 and S6, for instance.) This is due to various forms of stochasticity. One is random lack of mates (when *N*_*M*_ is small); another is local extinction caused by temporal population fluctuations (when *N*_*X*_ is small). These effects can be quite troublesome when setting up computational experiments across a range of parameters, especially if we want constant total population size.

Apart from those issues, the most common reasons for a significant discrepancy between the realized density and the “desired” density (set by *K*) have to do with when and how density is measured. First: *when* is density measured? In each time step, density increases after birth and decreases after death; *K* can match at most one of these times. Second: *where* is density measured? We naturally look at total population size divided by total area; however, on average individuals experience a higher density, since they themselves count towards their own local density. So, the correct comparison is of *K* to local population density averaged across the location of all individuals. Since this over-samples areas of higher density, this density will be higher than “total population size divided by area.”

All of these issues are discussed in much greater detail in Appendix B. For practical reasons it often suffices to simply be aware that population density is fundamentally an emergent property, determined in complex ways by nearly all parts of the life cycle. If a precise total population size is desired in a particular simulation, a simple solution is to adjust some parameter (*e.g*., the birth rate) until the desired value is achieved (see Box 6), but of course that will alter the dynamics of the simulation in other respects.

## 8 Natural selection

It turns out that there are many different ways to implement natural selection in an individual-based simulation that all map to the same abstract models from population genetics, yet produce distinct outcomes.

Modelers may choose to have an individual’s genotype affect its choice of mate, its preference for certain habitats, its longevity, or any number of other life history traits. These effects may differ across a landscape, leading to local adaptation. In this section we demonstrate selection that acts on birth and death, buliding on Section 2. Just as we could regulate population size in our individual-based simulation through birth rate, death rate, or both, one may achieve the same increase in “fitness” by increasing fecundity or decreasing mortality, and we will see that the selected allele thus affects equilibrium population density. The resulting models might be classified as having “hard” selection (Wallace, 1975); “soft” selection might act through increased or decreased chance of being chosen for mating (and would usually cause little to no change in equilibrium population density). For a view of the literature on these ecological-evolutionary models, see Mallet (2012) or Travis et al. (2023).

Much of population genetics selection theory is expressed in terms of the “selection coefficient” of a variant, usually denoted *s*, which is usually defined in the context of a Wright–Fisher model as the change in relative probability it confers on the individual to be “selected” to provide offspring to the next generation. The point in the life cycle when the variant confers its selective effect is known to produce differences even in nonspatial models (Bodmer, 1965; Nagylaki and Crow, 1974), and so analogies between the dynamics of such alleles and those in a Wright–Fisher model are necessarily approximate. This is important when attempting to match simulation results to theory: just because a variable that affects survival or fecundity is named *s* does not mean that using its value in expressions derived from the Wright–Fisher model correctly predicts the probability of establishment, mean frequency, or other classical quantities, even in a nonspatial simulation.

Below, we demonstrate two models in which selection acts on either mortality or fecundity. We choose our parameter *s* so that a variant’s frequency changes on average by a factor of 1 + *hs* per time step when rare, where *h* is the dominance coefficient. (Note that this may not be the best definition for use with classical formulas, which often measure change per generation.) For a historical and philosophical review of definitions and measures of selection see Endler (1986). Implementation in SLiM is described in Box 7.

### 8.1 Defining allelic growth rate

Recall from Section 2 that *f* (*u*) and *µ*(*u*) are, respectively, the per-capita mean number of offspring and probability of death per time step for an individual experiencing scaled density *u*. For selection, these vital rates should depend on the individual’s genotype. So, let *f*_*k*_(*u*) be the mean fecundity of an individual with *k* copies of a focal allele, and similarly *µ*_*k*_(*u*) the probability of death. When the focal allele is rare, most of the population use *f*_0_(*u*) and *µ*_0_(*u*), but a few individuals use *f*_1_(*u*) and *µ*_1_(*u*). So, then the per capita rate at which the number of focal alleles grows when at scaled density *u* in an outcrossing species is

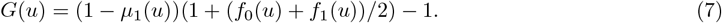

This is almost the same as the expression for *F* (*u*), the expected change in the number of *individuals* from equation (2), except that the fecundity term is (*f*_0_(*u*) + *f*_1_(*u*))*/*2; this is because the offspring of a heterozygous parent will only inherit the focal allele from that parent half the time.

### 8.2 Parameterizing selection

First we consider mortality-based selection. Suppose an additive allele (with *h* = 1*/*2) increases survival by *s/*2 per copy (*k*), so that

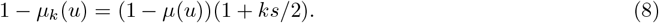

If the allele doesn’t affect fecundity (meaning *f*_*k*_(*u*) = *f* (*u*)), then we plug equation (8) into equation (7) with *k* = 1 to find that per-capita allelic growth rate in an outcrossing species is

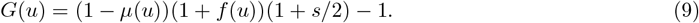

We’ve set things up here so that *s* means what we want: at the population’s equilibrium density, the allele’s growth rate when rare is *G*(1) = *s/*2.

Now suppose that the allele increases fecundity and does not affect survival (meaning *µ*_*k*_(*u*) = *µ*(*u*)). We define

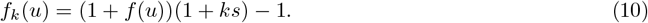

With this form for fecundity selection, the allelic growth rate when rare is the same as the rate we calculated for mortality-based selection. We can check this by plugging in our fecundity expression (10) into equation (7) with *k* = 1 and verifying that we get the same expression as equation (9). Therefore, at equilibrium, the per-capita growth rate of the number of alleles is again *s/*2. Roughly speaking, each allele increases fecundity by *s*, rather than the *s/*2 in equation (8) for mortality selection, because here the effects of fecundity are only affecting half of the parents (the offspring-bearing ones).

### 8.3 Spatial selection mechanisms in practice

Above we laid out deterministic, large-population-size theory that suggests that a variant that affects survivorship or fecundity might have similar frequency dynamics. However, in practice there are some meaningful differences. Figure 7 shows selective sweeps in spatial and nonspatial simulations. All simulations use density-dependent feedback on mortality (via equation (4)), but selection acts on either fecundity (via equation (10)) or mortality (via equation (8)). In nonspatial simulations, the global density (total number of individuals divided by total area) is substituted for local density when computing probability of survival. Some features of the sweep experiments are expected. Selective sweeps progress more slowly in spatially structured populations. The allele-frequency trajectories of *de novo* mutations under either mortality- or fecundity-based selection (Figure 7A), at first increase at the same speed in spatial and nonspatial sweeps, as predicted (*i.e*., *G*(*u*) is the same for both when the allele is rare). However, the selection mechanism introduces a few differences. As the beneficial allele fixes, equilibrium population density increases by a greater amount under fecundity-based selection (Figure 7B) than mortality-based selection. To understand why this is, consider the population after the fixation of the beneficial allele: all individual birth and death rates are now *f*_2_(*u*) and *µ*_2_(*u*). We are at a new equilibrium density *u*_*_, which occurs when net per-capita change in population size is zero. Solving equation (2) with our post-fixation vital rates (*F* (*u*_*_) = 0), the new equilibrium density 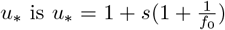 for mortality-based selection, and 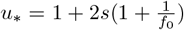 for fecundity-based selection. In other words, the increase in equilibrium density for fecundity-based selection should be roughly twice that of the increase in mortality-based selection.

**Figure 7:**
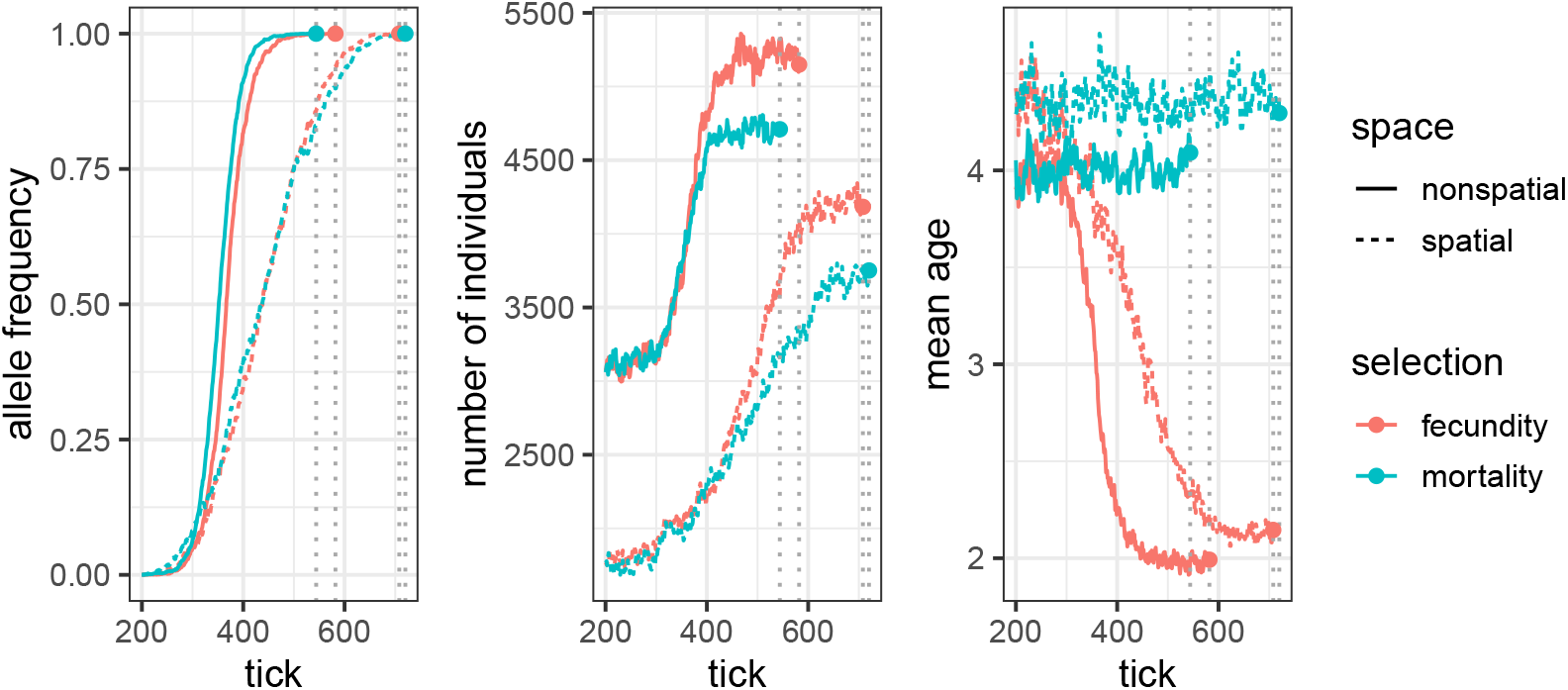
Allele frequency **(left)**, population size **(center)**, and average individual age **(right)** over time in spatial and nonspatial non-WF simulations. Curves end at the time the selected allele fixed (dotted vertical lines).

Fecundity-based selection also causes mean individual age to drop as the beneficial allele increases in frequency (Figure 7C). Population size increases as the fecundity-based sweep progresses, but because the beneficial allele does not confer protection against mortality, individuals are subject to the negative effects of increased population density.

Further differences between the spatial and nonspatial simulations are likely due to the spatial density regulation processes that cause realized population sizes in spatial simulations to be smaller than in their nonspatial counterparts. (See Section 7.1 and Appendix B.4 for discussion.)

## 9 Case Studies

How can we combine all the strategies we have seen thus far into a useful model of a living system? Here we illustrate how the spatial modeling framework we’ve introduced can be used to model complex scenarios such as (1) temporal change, (2) complex life cycles, (3) continental-scale systems, and (4) competition for resources.

We motivate each scenario with an organism. The scenarios are not meant to be complete or accurate models of the population biology of these organisms; rather, they illustrate how to apply the concepts presented in this paper to a study system. Additional methodological information for each vignette can be found in Appendix E and our code example repository https://github.com/kr-colab/spatial_sims_standard.

### Box 7

**Selection**

In nonWF SLiM models, the “fitness” property of an individual is the probability that the individual survives to the next time step. To make each mutation affect survival by a factor of `S MORT`, we simply declare:

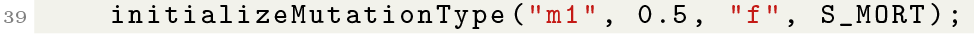

(The factor of 0.5 is the dominance coefficient: heterozygotes will have fitness multiplied by 1 + S MORT*0.5, and homozygotes by `1 + S MORT`.)

To have the same type of mutations also affect fecundity with selection coefficient `S FEC`, we need to account for genotype when setting up offspring numbers (as in Box 1):

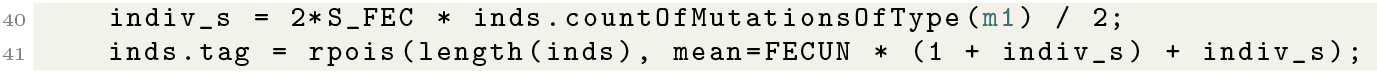

As written, the effects of any mutation of type `m1` on fecundity are additive across loci and copies; other arrangements are possible.

### 9.1 Changes over time: pikas and environmental change

Modeling spatial heterogeneity (as introduced in Section 6) makes simulations more realistic and informative, but sometimes *temporal* change in landscape variables is just as important. Here we explore how both seasonal fluctuation and globally rising temperatures affect population dynamics of an alpine organism: the pika.

Pikas (*Ochotona daurica*) are adapted to mountainous habitat at relatively high elevation, and cannot survive the extreme heat (or cold) at lower (or higher) elevations (Beever et al., 2010). Such ecology makes pikas potentially vulnerable as global temperatures are expected to increase over time.

#### Modeling approach

We incorporate three types of temporal change in temperature: (1) within-year seasonal change, (2) random, autocorrelated fluctuations between years, and (3) steady global temperature rise. Because temperature varies with elevation and elevation varies dramatically in mountainous regions, we model temperature as a function of elevation. We used a topographic map for a region of Rocky Mountain National Park in Colorado to calculate temperatures for each point in space and time during our simulation.

We connect temperature to fitness by killing offspring that are exposed to temperatures beyond the minimum (− 5^°^C) or maximum (28^°^C) sustained by pikas (Beever et al., 2010). Each tick of our simulation represents one year. To account for *within-*year seasonal variation (*i.e*., winter cold and summer heat), we narrow the viability range by the yearly variation in temperature (*s*^seas^), defining the probability of survival of a pika at location ***x*** in year *t* to be

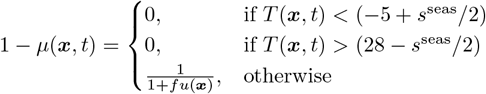

where the last term is our usual Beverton–Holt mortality regulation with local scaled density *u*(***x***) = *n*(***x***)*/K* (see equation (4)). In this equation, *T* (***x***, *t*) represents the midpoint of seasonal temperatures in year *t*, defined below, and the within-year variability *s*^seas^ broadens the temperatures experienced by the pikas around that midpoint. For example, if *T* (***x***, *t*) were − 4^°^C (inside the viability range), but seasonal temperatures varied by 4^°^C, the winter would be − 6^°^C – cold enough to kill.

Between-year temperature changes are modeled by the temperature function

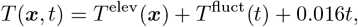

which combines elevation-related temperature change *T* ^elev^, correlated year-to-year fluctuations *T* ^fluct^, and global directional climate change.

*T* ^elev^(*x*) is calculated from the elevation map following fitted models from Collados-Lara et al. (2020). To create autocorrelated year-to-year fluctuation, we let *T* ^fluct^(0) = 0 and for *t ≥* 1,

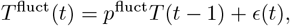

where *ϵ*(*t*) is Normally distributed noise with mean zero and standard deviation *s*^fluct^). The parameters *p*^fluct^ and *s*^fluct^ are known as persistence and shock, respectively, and determine how correlated the noise is between years. Finally, we increase the global temperature over time by adding 0.016^°^C per year (Foster and Rahmstorf, 2011).

#### Observations and extensions

The resulting simulation is a habitat suitability model in which the population’s geographic distribution moves toward higher elevation as the global temperature increases (Figure 8).

**Figure 8:**
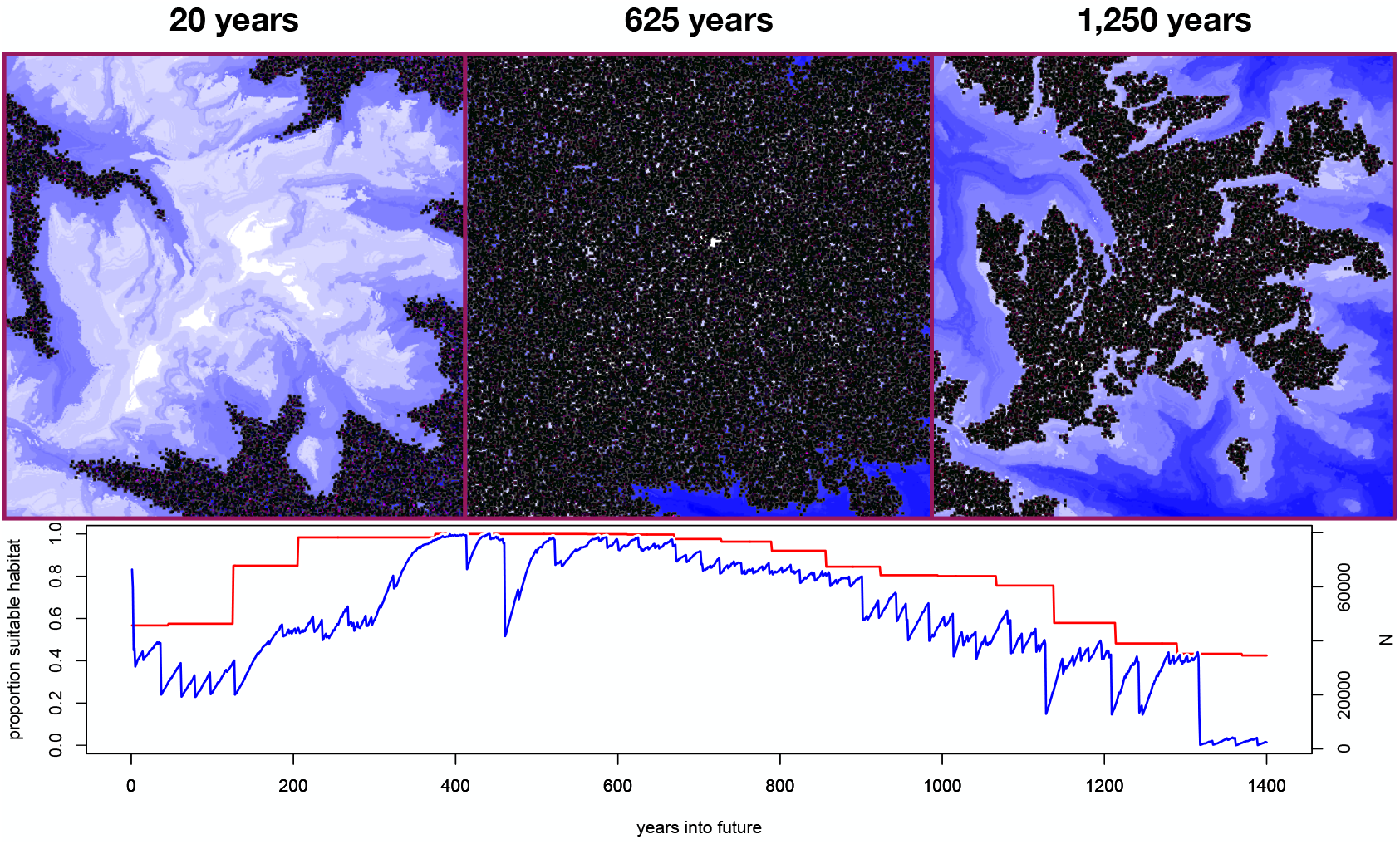
Pika simulation. **(top)** Screenshots of individual spatial positions (black) at different time points. The background image shows elevation, where blue and white correspond to lower and higher elevation, respectively. **(bottom)** The blue line shows population size over time; the red line shows the proportion of habitable space before the addition of random noise.

We found that if the magnitude of random variation around the expected annual temperature is large (*s*^fluct^ = 5^°^C) the probability of extinction increases significantly, particularly for early generations. In other words, one or a few bad years was devastating for pika populations. This result may provide a useful lesson: even if a species appears to be thriving, the long-term success of the species is not guaranteed.

Surprisingly, simulated population size increased in the intermediate-term future, since for the first few hundred years, habitat losses at lower elevation were more than compensated for by habitat gains at higher elevation. This illustrates that results from simulation on a specific map may not be generalizable.

Here, we use an empirically-informed linear trend with Gaussian noise as our global temperature change schedule. An alternative strategy for simulating temperature changes would be to pre-process multiple temperature maps reflecting different years, and continually feed the simulation new maps over time. Such maps could be generated directly from a climate model, rather than using a direct function of elevation.

### 9.2 Life cycle stages: mosquitos in Burkina Faso

So far, our models have not taken life cycle stages into account; individuals have been able to mate immediately after they are born, and their survival has not been age-dependent. However, for many organisms their patterns of mobility, competition for resources, and mating capability are age-dependent. Here, we demonstrate how to simulate a population with distinct juvenile and adult phases. Specifically, we set up a simulation of mosquitos in Burkina Faso in West Africa, inspired by North et al. (2019).

#### Modeling approach

In this model, individuals begin life as juveniles, and mature into adults after a fixed time span. Population regulation and individual behavior depends on the life stage. For adults this is a constant survival probability of 0.875 per day (time step), independent of the landscape map. The population is regulated by density-dependent survival of the *larvae*, which varies across the map.

Following the model in North et al. (2019), we set the carrying capacity of larvae based on water availability. Larval carrying capacity is only non-zero at the outlines of water features extracted from GIS data of inland water in Burkina Faso. There, the carrying capacity fluctuates seasonally to mimic rainy and dry seasons:

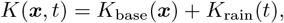

where *K*_base_(***x***) is obtained from the map of inland water, and *K*_rain_(*t*) is a sinusoidal function with a period of 365 days and a minimum value of 0.

We then use a Beverton-Holt form for the survival probability of larvae. Since there are ten age classes for larvae (and survival of each depends on the total number across all ages), parameterizing the model so that local density is (roughly) *K* involves solving a system of equations (in a matrix model) described in Appendix E.2.

A female adult mosquito mates with an adult male within the maximum mating distance, and lays eggs by sampling a location within her dispersal radius weighted by carrying capacity. Each day, adult mosquitos move by a random displacement sampled from a Gaussian distribution, whereas larvae do not move from their original location until they mature.

#### Observations and extensions

This model simulates a mosquito population with structured life history. The population size of larvae and adults fluctuate periodically, following precipitation levels with a slight lag (Figure 9). Recall that adult mortality is simply constant, so the periodic fluctuations of adults are mediated through larval carrying-capacity dynamics.

**Figure 9:**
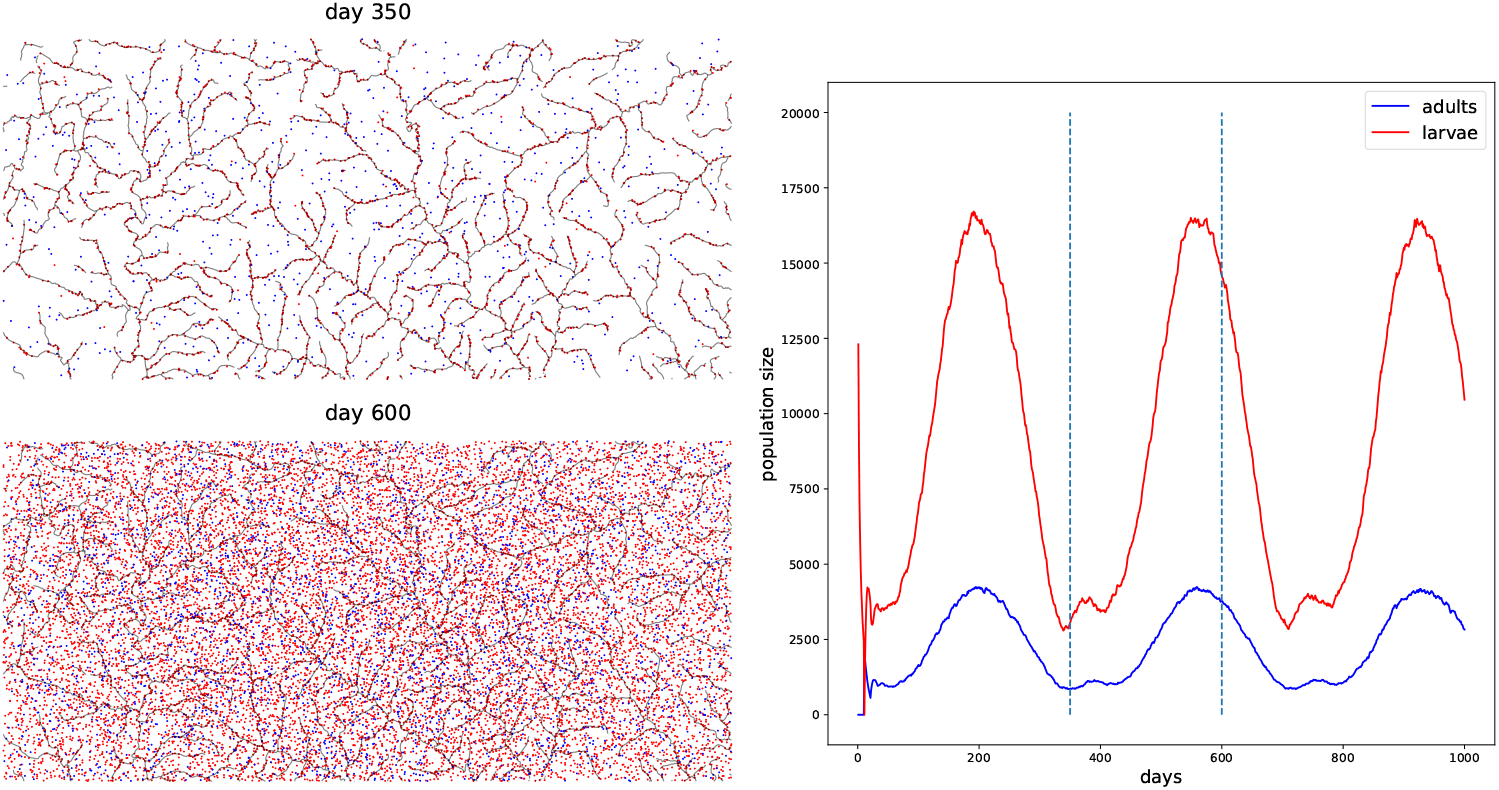
**(left)** Adult (blue) and larval (red) populations on the river map of Burkina Faso, at two time points during the year: in the dry season (day 350), larvae can only survive in bodies of water, while in the wet season (day 600), larvae can survive in many places. **(right)** Adult and larval population sizes oscillate with the seasonal cycle. Vertical lines indicate the time points plotted on the left.

There are a few immediate extensions to our simulation. Here, we did not distinguish between perennial and fluctuating water sources. Simulating water sources that appear and disappear (adding a *t* argument to *K*_base_(***x***)) could result in extinction dynamics such as those observed in the pika example above. Similarly, the amount and duration of rainfall could be location-dependent (adding an ***x*** argument to *K*_rain_(*t*)).

We set the maximum of the *K*_base_ and *K*_rain_ functions to 0.002 individuals per square meter, which is probably much lower than in nature, to keep the memory usage of our simulation low. To run a simulation with a realistic (huge) number of mosquitoes we would need to make some efficiency improvements. Fortunately, an example of the necessary techniques follows.

### 9.3 Continental-scale systems: invasion of the cane toads

There is a perception that individual-based spatial models are so slow that they are meaningfully limited in the population size that can be modeled. Our goal with this vignette is to demonstrate that this is not necessarily the case, by showing how to model large-scale populations and landscapes with relative ease and efficiency.

Cane toads (*Bufo marinus* or *Rhinella marina*) are native to Central and South America, and were intentionally introduced to the Northeast coast of Australia in 1935 as a pest-control measure. Since their introduction, cane toads have experienced explosive population growth, with hundreds of millions of individuals spreading over several million square kilometers, resulting in considerable negative economic and ecological impacts (Shine, 2010; Urban et al., 2008).

#### Modeling approach

We use information about the biology of cane toads, when available, to parameterize the model. Toad population densities in established populations have been estimated around 8000 per square kilometer (Freeland, 1986). Based on this, and then tailored heuristically over several trial runs to be computationally feasible and to produce a reasonably realistic pattern of range expansion, we settled upon a local carrying capacity of 1000 per square kilometer. Telemetry data shows cane toads can travel up to 0.2 km per day (Shine et al., 2021). In order to interpret simulation time steps roughly as years, we set the spatial scale parameters *σ*_*D*_, *σ*_*X*_, and *σ*_*M*_ to 20 km. For dispersal, we use a Student’s *t*-distribution, which provides more long-range dispersal events than a Gaussian kernel (Figure 3C).

We model survival probability as a function of precipitation due to its similar distribution to empirical occurrence data, which seems reasonable given cane toads’ known sensitivity to moisture conditions (Child et al., 2009; Cohen and Alford, 1996). Specifically, we multiplied our typical density-dependent survival probability 1 −*µ*(*u*) by 1*/*(1 + exp(− (*α* + *βP* (***x***))), where *α* and *β* control the intercept and slope of the survival curve, and *P* (***x***) is the amount of yearly precipitation at location ***x***.

The invasion began with 10,000 individuals (though the number of individuals actually released in 1935 was likely greater (Shine et al., 2020)) at locations randomly sampled from the first four years of the observed occurrence data.

Indeed, mating and interaction neighborhoods were large, which initally led to prohibitively long runtimes. To make the simulation more efficient, we followed the approach described in Box 8, modified to measure the local density of individuals per unit of *habitable* area. Mate choice was also modified as described in Box 9.

#### Observations and extensions

We were able to approach the true spatial and population scale of the cane-toad invasion, with a final census size of about 120 million individuals, nearing the estimated modern census size of Australian cane toads (Australia, 2019), after running for about 5 days and using 200 GB of RAM at maximum. We visually compared empirical occurrence data for cane toads to our simulations, with and without annual precipitation’s effect on survival (Figure 10). While there are obvious differences in densities and locations between the simulated and observed data, it is clear that modeling annual precipitation’s effect on survival greatly improves the likeness.

**Figure 10:**
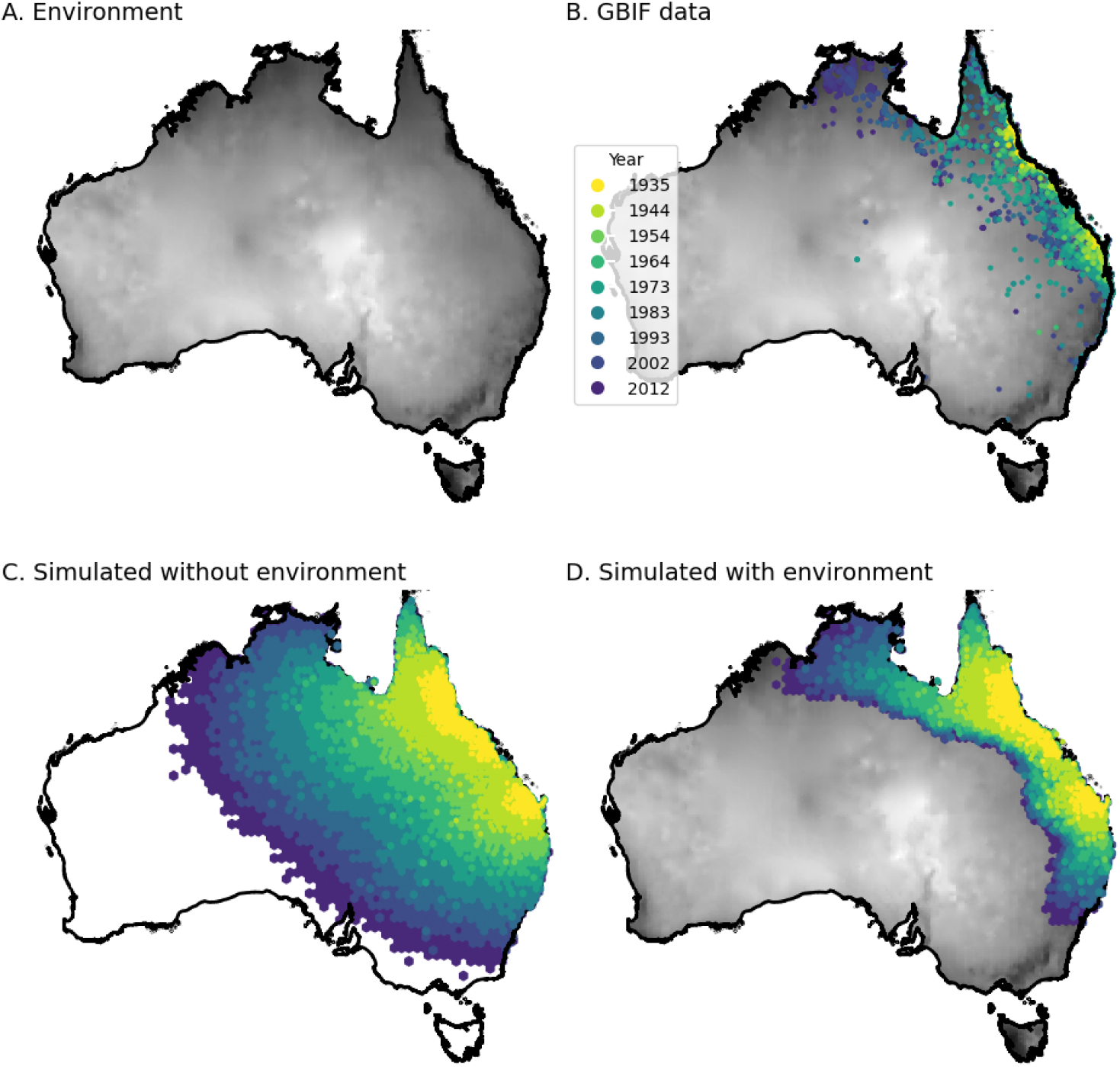
Simulating the cane toad invasion with and without and effect of annual precipitation on survival. **(A)** Map of Australia shaded by annual precipitation (kg m^−2^ year^−1^). **(B)** Observed distribution of cane toads from Global Biodiversity Information Facility (GBIF) **(C)** Simulation of cane toad invasion without effect of annual precipitation on survival. **(D)** Same as (C), with effect of annual precipitation on survival.

The approach described here is similar to classical niche modeling (Peterson, 2001), which has been used extensively to model cane-toad distributions (Shine, 2010), but is somewhat simplified by only using one environmental variable. This would be a straightforward extension: multiple environmental variables could be combined into a single habitability map for use in the model, with no impact on runtime performance. The benefit of this vignette’s approach over previous approaches is the way that it combines information from the explicit individual-based simulation and environmental data, producing a simulation at a realistic scale with respect to both landscape size and population size.

### 9.4 Resource-explicit competition: monarchs and milkweed

In the preceding examples we have regulated populations through competitive interactions between individuals: either explicitly, as in Box 1, or in a space-averaged manner, as in Box 8. Population regulation in this model is managed quite differently. This is a “resource-explicit” model, wherein the population is extrinsically regulated by the availability of an external resource (Champer et al., 2024), as outlined in Box 10.

We simulate monarch butterflies (*Danaus plexippus*) and the milkweed (*Asclepias* spp.) plants on which they lay their eggs. Though adult monarchs feed on nectar from numerous types of plants, monarch caterpillars are specialists that eat only milkweed (Oberhauser et al., 2004).

#### Modeling approach

Monarchs in the model progress through three life-cycle stages: caterpillar, pupa, and butterfly. Mortality is modeled differently in each stage. The food resource, milkweed, is directly included in the model as a second species.

During the first two weeks of their lives (time steps 0 and 1), caterpillars interact with nearby milkweed plants and accumulate resources. The amount of resources collected from a milkweed plant each week is inversely proportional to the number of competing caterpillars within a given interaction scale (*σ*_*X*_) of that plant. This means that a plant fed upon by more caterpillars will be depleted more quickly. When individuals reach their third week, they enter the pupa phase. At this time, survival is proportional to the amount of milkweed eaten as a caterpillar. (The survival probability is calculated only once when it becomes a pupa.) Surviving pupae become butterflies during their fifth week. Butterflies disperse, reproduce, and experience mortality at an age-dependent rate. Mortality is not density-dependent during the butterfly stage. Figure 11 shows a snapshot of the simulation, highlighting the dispersal and reproduction of individuals across the landscape.

**Figure 11:**
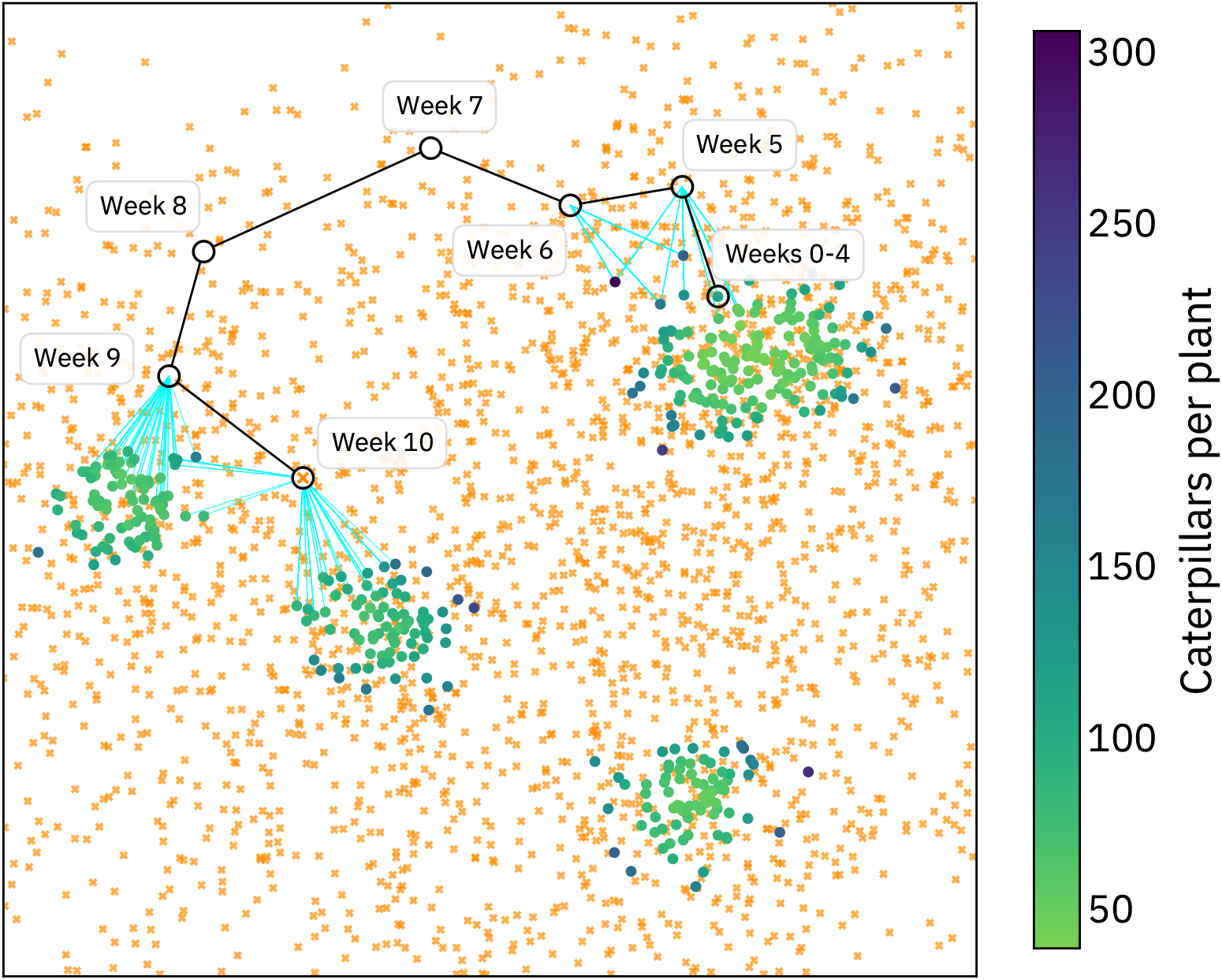
A portion of the landscape with butterflies (orange) and milkweed (green). There are about 40,000 caterpillars and 400 milkweed plants in this area. Each plant provides sufficient resources to allow an average of two caterpillars to survive to adulthood per week. A total of 2,680 butterflies are present in this area. The life history of a single individual is tracked from its origin on a particular milkweed plant, with black lines depicting dispersal, and teal lines showing where this individual laid eggs.

Reproduction also involves a resource-explicit interaction, since monarchs only lay their eggs on milkweed. To accomplish this, a spatial interaction is evaluated between adult males and milkweed (with a much longer range than the interaction between caterpillars and milkweed); each milkweed caches a list of nearby males from that interaction. Next, a similar spatial interaction is evaluated between adult females and milkweed; females are iterated through, with each female selecting a mate from the males cached at the milkweed plants within the female’s interaction range. Finally, the females randomly distribute their eggs at the plants where matings occurred.

##### Box 8

**Using maps for faster spatial interactions**

In Box 1, we estimated the local population density for each individual with equation (1), using the `localPopulationDensity()` method. For each individual, this method sums the “interaction strength” (*i.e*., kernel density) for every other individual within the provided maximum distance. So, if the total number of individuals is *T* and the typical number of neighboring individuals is *N*_*X*_ (defined in Section 3), then the complexity of this operation is *TN*_*X*_ . If each individual has a large number of neighbors, this can be quite costly. However, if the number of neighbors is large, it should work just as well to (a) create a (discretized) map of local density, and (b) look up the value of the local density experienced by each individual on that map. Map lookup is quick, so if the cost of creating the map is smaller than *TN*_*X*_, we will have a more efficient model.

We can create a map whose value at ***x*** is (approximately) given by equation (1) in two steps: (1) use `summarizeIndividuals()` to measure the number of individuals per unit area in each cell in a grid (see Box 2), and (2) `smooth()` this map using the appropriate kernel, so that the density value at a given point in the resulting map depends upon an appropriate weighted average of the individuals per unit area across nearby cells of the grid. Using the same kernel as in Box 1:

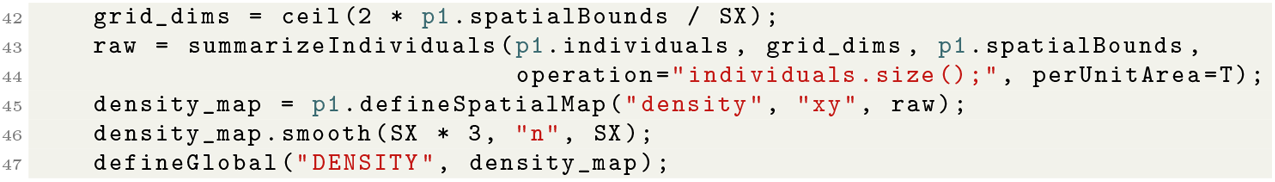

Then, we can modify the code of Box 1 to use the map instead:

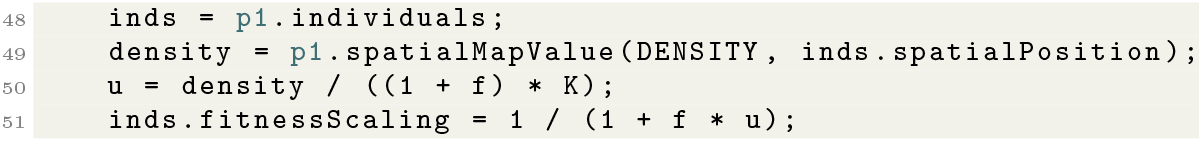

This obtains (nearly) the same value as would `localPopulationDensity()` if the resolution of the map should be finer than the scale over which the density kernel changes. In this example, that scale is `SX`, as in *σ*_*X*_, so we have ensured that the map has cells of size smaller than `SX/2`. The approximation is examined in Appendix F.

#### Observations and extensions

In addition to reflecting the life history of monarch butterflies, the resource-explicit modeling approach is highly performant. Each female monarch can lay several hundred eggs, very few of which go on to reach adulthood. As a result, the number of caterpillars in the model far exceeds the number of milkweed plants. Thus, regulating the size of the caterpillar population by evaluating a spatial interaction between caterpillars and plants is far more efficient than regulating the population by evaluating an interaction directly between caterpillars.

This approach also provides an intuitive way to investigate questions related to resource availability. For example, this model could probe the effects that milkweed habitat loss, caused by urbanization or climate change, could have on the viability of the local monarch population.

##### Box 9

**Using maps for faster mate choice**

A similar problem as in Box 8 arises when choosing mates: even though only one mate needs to be chosen, the underlying operation is of order *TN*_*M*_, where *T* is the global population size. The same map of density can be used to solve this problem as well: instead of choosing an individual with probability proportional to a kernel, it is (nearly) equivalent to: (1) choose a point in space nearby, with probability proportional to the map multiplied by the kernel, and then (2) take the individual closest to that point. Recall the number of possible mates scales as 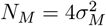 *K* from Section 3. This implies that the number of neighbors grows linearly with *K*. Thus, we can keep the number of potential mate roughly constant regardless of how large local density is by rescaling the maximum distance in the `InteractionType` used for mate choice by 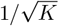 from the code in Box 4:

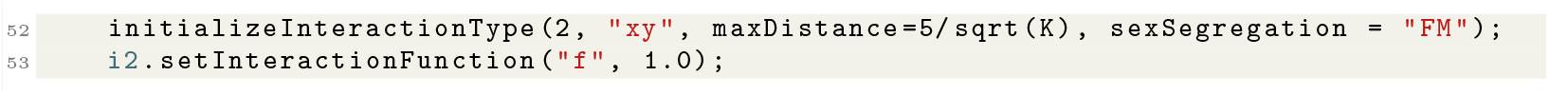

We’ve set the maximum distance in the interaction kernel to be a value that should give us around 25 neighbors for each individual; however, if density varies significantly across the landscape, this may make some individuals in low density areas fail to mate. Then, we choose the mate as follows:

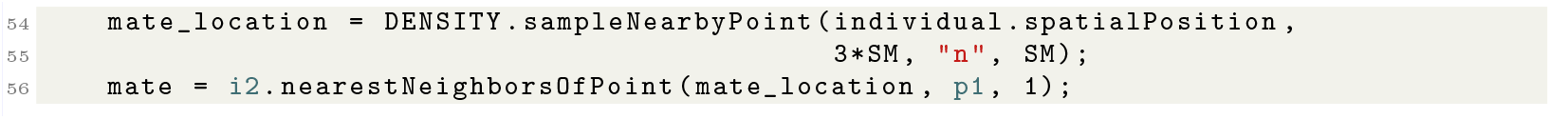

Here the specification of a Gaussian mate-choice kernel with standard deviation `SM` has moved from the definition of the `InteractionType` to the `sampleNearbyPoint` call: given a location *x*, a map with value *m*(*y*) at *y*, and a kernel *ρ*(), this returns a random point *z* sampled with probability proportional to *m*(*z*)*ρ*(*z*). We then choose the mate as the individual nearest to that point. The approximation is examined in Appendix F.

## 10 Discussion

The first spatial models intended to represent landscapes (as opposed to, say, two-deme island-mainland models) were based on partial differential equations and so did not explicitly represent individuals (Skellam, 1951; Beverton and Holt, 1957; Cantrell and Cosner, 2004). Many others have used arrays of discrete populations (*e.g*., metapopulation models, Hanski, 1997), or had individuals living on a regular grid (Epperson, 2003). Although there are use cases for these, the main reason that individual-based continuous-space models are not more common may simply be convenience, both for mathematical analysis and software programming. The continuous-space formulation we use was introduced by Pacala (1986) and Bolker and Pacala (1997), and has been used in many theoretical studies (Dieckmann and Law, 1999; Snyder and Chesson, 2004; Etheridge et al., 2024).

A more modern extension to matrix population models, integral projection models, more commonly incorporate density dependence (Ellner et al., 2016), and may even estimate functional forms – see for example Adler et al. (2010). Much of the remaining literature on density dependence only considers the form of *F* (*u*), rather than separating out the effects of density on different life history components (reviewed in Caswell, 2000; Eskola and Geritz, 2007). There are a wide variety of methods to infer *F* (*u*) from time series data (Lande et al., 2003), but statistical issues make the problem difficult (Freckleton et al., 2006). However, the “*F* (*u*)” thus inferred is not necessarily the same as ours – many of these methods assume no demographic stochasticity, and hence an infinite population size. In other words, the *F* (*u*) that these infer is a *landscape-scale* relationship, averaging the net effects of birth and death over thousands or millions of individuals. Our *F* (*u*), on the other hand, is local, and describes how a single individual is affected by having a few more or less neighbors. These are related, but need not have the same functional forms. In particular, the behavior of *F* (*u*) for very large or small values of *u* is usually much more important when applied to an individual, because the local density around an individual might be proportionally much larger or smaller than average, due to random fluctuations, than the total population size across a landscape. These can have a strong effect - for instance, increasing population density can have a positive effect on population growth (for at least low enough densities), a dynamic which can have interesting and important effects (Courchamp et al., 2008).

### Box 10

**Resource-explicit foraging and mortality**

In this box, we outline the “resource-explicit” modeling approach described more fully in Champer et al. (2024). This approach implements density-dependent population regulation that is mediated indirectly through the availability of a resource, rather than directly through competitive interactions between individuals. A simple resource-explicit simulation contains two species in a multi-species SLiM model: the focal species, in subpopulation `p1`, and a species representing the resource, in subpopulation `p2`, in which individuals represent patches of the modeled resource.

During the foraging phase, individuals collect resources from nearby patches. Each patch can each support a certain number of individuals per time step. If a patch can support 10 individuals, but there are 100 individuals nearby at a given time step, each individual will receive 10% of the amount of resource necessary to guarantee survival (but might also forage from other nearby patches). The amount of resource that each individual collects during the foraging phase is stored in the `tagF` property.

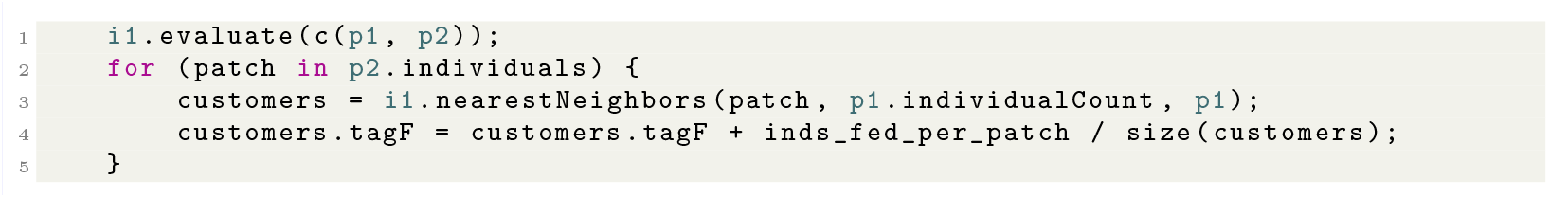

After foraging, individuals die if they have not consumed enough resources. In the code example below, this occurs when individuals reach age 2.

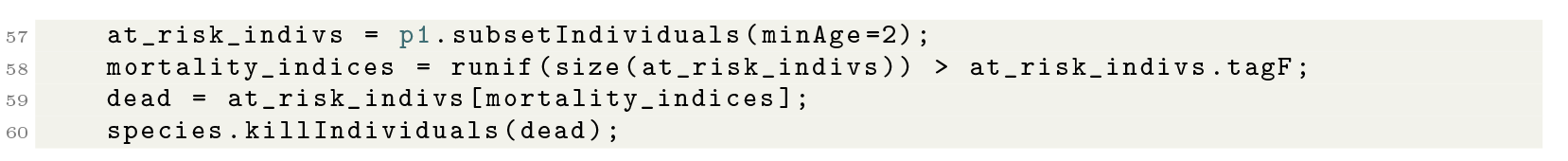

If an individual collected at least one unit of the resource, its survival to the next time step is guaranteed. Otherwise, it survives with a probability equal to the total amount it collected. See Appendix F for discussion of the relationship to the method of Box 8.

Furthermore, effects of density dependence are often mediated (at least in part) through interactions with other species, and the nature of these interactions may depend on environmental context. These interactions are often modeled as the net result of pairwise interactions, and various methods are used to estimate these potentially numerous and environmentally-dependent effects (for recent examples, see Weiss-Lehman et al., 2022; Bimler et al., 2023). It is beyond the scope of this article to review the full range of approaches and possibilities – but, note that there is no obstacle to simulation of multiple species whose interactions vary across space and/or time in SLiM (Haller and Messer, 2023).

### Discretized continuous space

There is not a hard distinction between “continuous” and “discrete” spatial models. However, in practice discretized models usually only allow density-dependent regulation to act *within* demes: in other words, vital rates of an individual can only depend on the other individuals in the same deme. This is important because the degree of demographic stochasticity depends on deme size; therefore, different discretizations may produce different amounts of stochasticity. What demographic stochasticity there is is shared among all individuals in the same deme, reducing the overall variance in the simulation. The effects of these discretization artifacts are not well-described (but see Battey et al., 2020). If demographic rates within each deme instead depended on nearby demes, then this would not be an issue. However, we are not aware of any simulators that make this choice.

### Empirical estimation of density dependence

Huge amounts of observational effort have gone into detailed estimates of the demography of particular species, especially those of conservation or management concern. However, such observations by necessity describe a snapshot of demographic rates in particular conditions (or, averaged over a particular range of conditions). Estimating the functional responses to density necessary to describe long-term dynamics is much more difficult, although methodological progress has been made, for instance, by incorporating many sources of information with “integrated population models” (Zipkin et al., 2023). A great deal of ecological work has also sought to quantify the effects of density-dependent demographic feedback. In a spatial model, this population density is usually measured locally – around the individual in question – yielding concepts such as the “competition kernel” (Bolker and Pacala, 1997) or “crowding index” (Pacala and Silander, 1985). Density-dependent feedback has a key place in the theory of coexistence between species: the Janzen-Connell hypothesis suggests that stronger intra-than inter-specific negative density-dependent effects could provide a mechanism for species coexistence (Janzen, 1970; Connell, 1971; Terborgh, 2012; Hülsmann et al., 2021). A substantial number of studies across a variety of organisms (mostly plants) have quantified these effects, both within species (*e.g*., Weiner, 1982; Silander, Jr and Pacala, 1985; Specht and Arnold, 2018; Spotswood et al., 2017) and between (*e.g*., Mack and Harper, 1977; Song et al., 2021; Zaiats et al., 2021). Estimation of density-dependent interaction strengths between many species in a community is challenging, but important for understanding community assembly (*e.g*., Weiss-Lehman et al., 2022; Bimler et al., 2023). Adding to this complexity, the effects likely often depend strongly on age (Richardson et al., 2024).

Researchers in plant ecology have made the most progress towards empirical understanding of the mechanistic underpinnings of the sort of local density dependence we require. In practice there are a great many possible ways to quantify the cumulative effect of the neighbors of a single individual (Weigelt and Jolliffe, 2003). Our formulation here treats all individuals equivalently, but in practice one could include the effects of age or size. For instance, a common method in forestry modeling defines a “neighborhood competition index” for a given tree as the sum over all neighbors of their diameter (to some power) divided by distance (to another power) (Bella, 1971; Daniels, 1976; Canham et al., 2004). Empirical studies have estimated these kernels in a variety of situations: for instance, Teller et al. (2016) and Adler et al. (2018) use spline methods to flexibly estimate the effects of total area of nearby plants on a target plant’s growth and survival.

### Empirical estimation of dispersal

The concept of a dispersal distribution (as discussed in Section 4) is perhaps most well-defined for plants, which (mostly) have only one opportunity to move during their lifetime, as a seed or other propagule. Seeds and pollen can be moved by gravity, wind (Nurminiemi et al., 1998), water (Murray, 2012), or animals (Morales and Morán López, 2022; Pons and Pausas, 2007), and complex models for these have been developed and estimated from empirical data (Neubert et al., 1995; Tufto et al., 1997; Austerlitz et al., 2004; Katul et al., 2005). Of course, many animals have relatively stable locations as well, perhaps depending on the season: for instance, Paradis et al. (1998) review estimates of post-natal and breeding dispersal in many bird species. The combination of these various processes (which often happen across many different spatial scales) has been referred to as the “total dispersal kernel” (Rogers et al., 2019), and can easily lead to “long-tailed” distributions (Cain et al., 2000; Edwards et al., 2007) in which rare, longdistance events are important. Dispersal is notoriously hard to estimate, as it often requires observing rare events, but has been done in a variety of organisms including kangaroo rats (French et al., 1968), mosquitoes (Estep et al., 2014), *Drosophila* (Dobzhansky and Wright, 1943), *Prunus* shrubs (Robledo-Arnuncio and García, 2007), pine pollen (Robledo-Arnuncio and Gil, 2005), and butterflies (Suchan et al., 2024). Dispersal can of course also depend on density (Harman et al., 2020) (*e.g*., if individuals preferentially disperse out of crowded areas). Clobert et al. (2012) review many aspects of dispersal, from parameterization and estimation to implications for ecology and evolution. See also Saastamoinen et al. (2018); Edelaar and Bolnick (2012).

## 11 Conclusion

Individual-based simulations are a powerful method for studying how demographic and population-genetic processes operate over continuous geographic space. Modelers must design rules for how individuals in the simulation interact with others nearby, and how forces such as selection operate. Individual-based simulations are well-suited to this problem, since they are very flexible, and can be tailored to a specific research system. However, flexibility can be both a blessing and a curse: it is easy to design a simulation with unstable population dynamics or unrealistic life-history traits. Similarly, stochasticity can cause a population that is intended to equilibrate to instead die out. Such problems likely reveal a flaw in our understanding of the system being modeled.

Here, we provide guidance and connections to the ecological literature for researchers interested in designing stable, efficient, and interpretable spatial simulations. Interpretability is a key advantage to spatial, individual-based simulations, since it can take substantial effort to translate the results of more abstracted models back to the domain of interest.

Realism is not a goal of our case studies, but each illustrates the degree of realism that can be obtained from SLiM without serious effort. Increasing computational efficiency and flexibility of simulation engines are bringing individual-based simulations closer to realistic models of the ecology of specific systems. Careful implementation of ecologically realistic evolutionary models will be important to many applied fields, such as understanding and predicting how climate change affects organisms’ ranges, predicting the consequences of a gene-drive release in the wild, and rescuing species close to extinction. As the scale and specificity of *in silico* models improves, individual-based simulations will become valuable tools beyond academic research for management professionals in conservation, management, and public health.

### Modeling density dependence

Setting out to write this paper, we hoped to provide a comprehensive yet simple guide to best practices in implementing density-dependent population regulation. Although we have provided one or two paths forward and elucidated many of the issues (see in particular Appendix B), careful empirical practitioners will soon encounter additional questions. What are some flexible and robust families of functional forms, and what aspects of these matter in practice? How can these be parameterized so that parameters naturally correspond to observable/interpretable quantities? How can these be fit to data? How should local habitat quality and density interact? At first, we imagined that the answers would be found in familiar names, and so the relationship between local density and fitness would be described as logistic, Beverton–Holt, Ricker, *et cetera*. However, we quickly found that these models were developed to describe population-level net changes, and so not only do not account for individual-level stochasticity, but furthermore do not separate birth from death. This is an area of active work – see, for instance, Aoyama et al. (2022) and Adler et al. (2018) for recent good examples. Full exploration of these questions was too much for this paper.

### Spatial data and niche modeling

Several of our case studies use environmental variables to specify where on the simulated landscape organisms are most likely to live. The explosion in remote-sensing data provides many potential data sources for modeling spatial heterogeneity. However, what is often needed in a model is a composite proxy for “suitability” that can be incorporated into local demographics. The process of predicting where a species might or does live is known as Ecological (or, Environmental) Niche Modeling (Booth et al., 1988; Peterson, 2001). This can be done in a variety of ways; for instance, one might model either the potential or the realized niche, and predict probabilities of occurrence or population densities (reviewed in Sillero, 2011). Environmental niche models are often used to predict suitable habitat either in other locations or other time periods (Werkowska et al., 2017; Yates et al., 2018), but resulting estimates can vary widely in quality and there are a number of statistical pitfalls (Sillero and Barbosa, 2021) that simulation testing could diagnose and simulation-based inference could potentially help avoid.

While ecological niche modeling uses observation data to predict where organisms might live, a collection of landscape-genetics tools try to use genetic relatedness to predict where organisms move. The methods generate a map of “landscape resistance” that aims to describe how easily individuals move over different parts of the landscape (McRae et al., 2008). However, resistance models rely on correlating genetic distance to an abstract notion motivated by electrical circuits (reviewed in Peterson et al., 2019; Cruzan and Hendrickson, 2020). They lack an underlying mechanistic model, so estimates can be problematic in practice (Cushman et al., 2013; Graves et al., 2013), and are expected to mislead in plausible situations such as biased dispersal (Lundgren and Ralph, 2019). Again, simulation-based inference provides a promising route forward (Smith et al., 2023, 2024), since it does not rely on explicit likelihoods or other mathematical descriptions.

### Sampling

Any simulation study that wishes to make comparisons to real data needs to also consider the sampling effort that led to those data, and realistic simulation of many sampling schemes can be daunting. In practice, sampling can strongly affect results (for an example in population genetics, see Battey et al., 2020). However, it is relatively easy to assess the robustness of results to variations in sampling scheme. Furthermore, it is often possible to “over-sample” simulations. For instance in Smith et al. (2023), many simulated datasets can be obtained from each costly spatial simulation, simply by repeating the sampling effort (however, if sufficient simulations are not done, model performance will be poor). It would be useful to develop a standard set of tools that implement various sampling schemes for simulated spatial populations.

### The future

Although the spatial simulations we present here incorporate many more aspects of real organisms’ lives than does the Wright–Fisher model, there are many things that we have not tried to explicitly model, such as seasonal migration, herding or flocking, territoriality, foraging strategies, microhabitat variation, broadcast spawning, resource storing, pollination, predation, facilitation, and other inter-species interactions. Any of these can be modeled in SLiM with more or less effort, and indeed many are described in the SLiM manual (Haller and Messer, 2024). The decision of which aspects of biology to model is in practice made by prioritizing those aspects expected to substantially affect the question being studied. We are excited to see the wide variety of simulations that researchers develop in the future, as we explore these questions and build on each other’s work.

## Acknowledgements

Many thanks to Jeremy Collings and two anonymous reviewers for useful and insightful comments on the manuscript, and to Erin Landguth for input on landscape simulation methods. This work was funded in part by NIH/NHGRI grant R01 HG012473 to PLR, ADK, BCH, and PWM, NIH/NIGMS grants R35 GM148253 to ADK, R35 GM139628 to SR and ETC, R35 GM152242 to PWM, and F32 GM146484 to CS. ETC also acknowledges support as a trainee under the Brown University Predoctoral Training Program in Biological Data Science (NIH T32 GM128596).

## Data availability

SLiM scripts suitable for re-use of all simulations used in this paper are at https://github.com/kr-colab/spatial_sims_standard (for SLiM v4.2 Haller and Messer, 2023). Scripts to produce the figures in this manuscript are available at at https://github.com/kr-colab/spatial_sims.

## Appendix A Example SLiM script

Here is a complete SLiM script for a spatial simulation with local Beverton-Holt regulation on mortality.

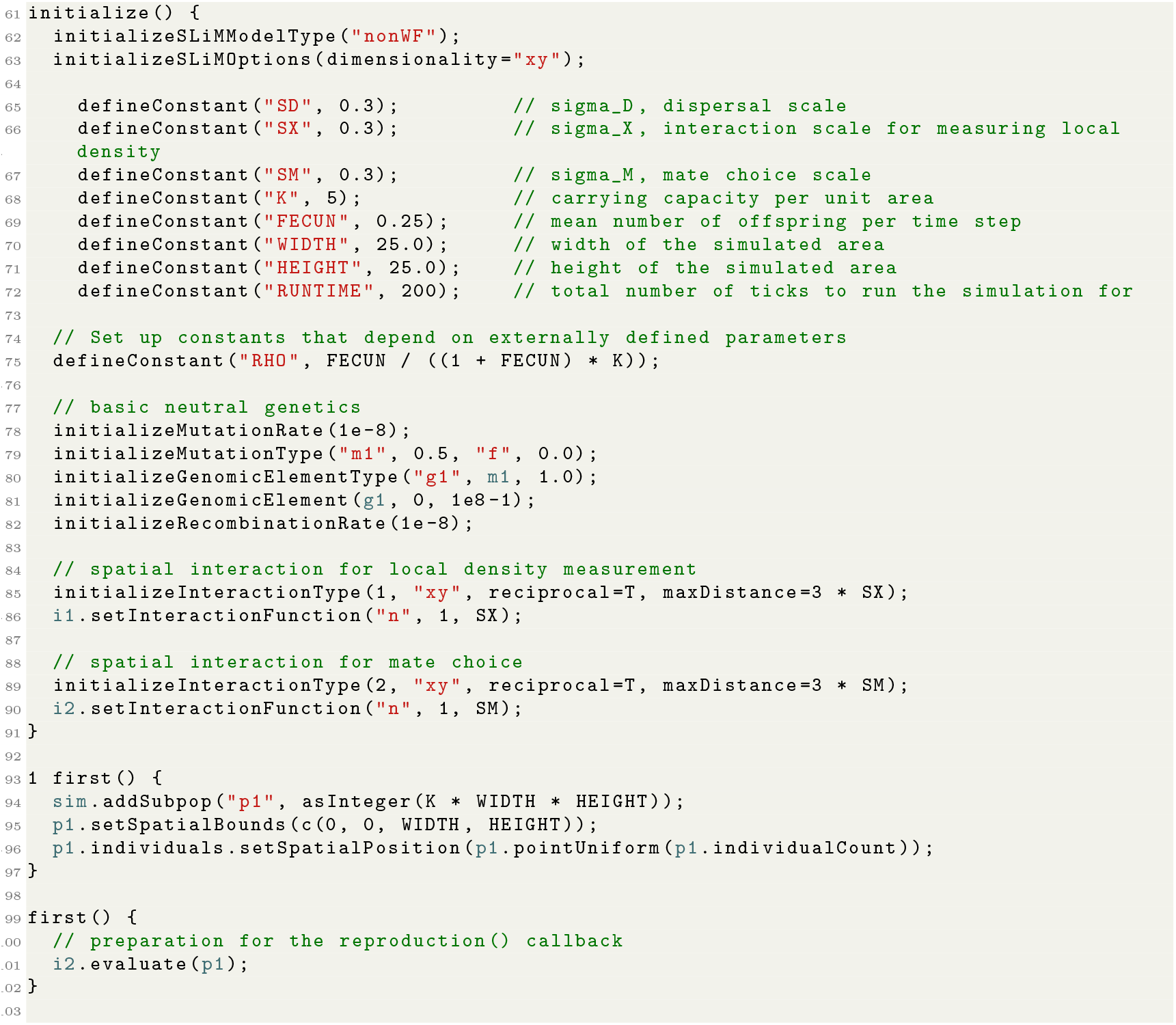
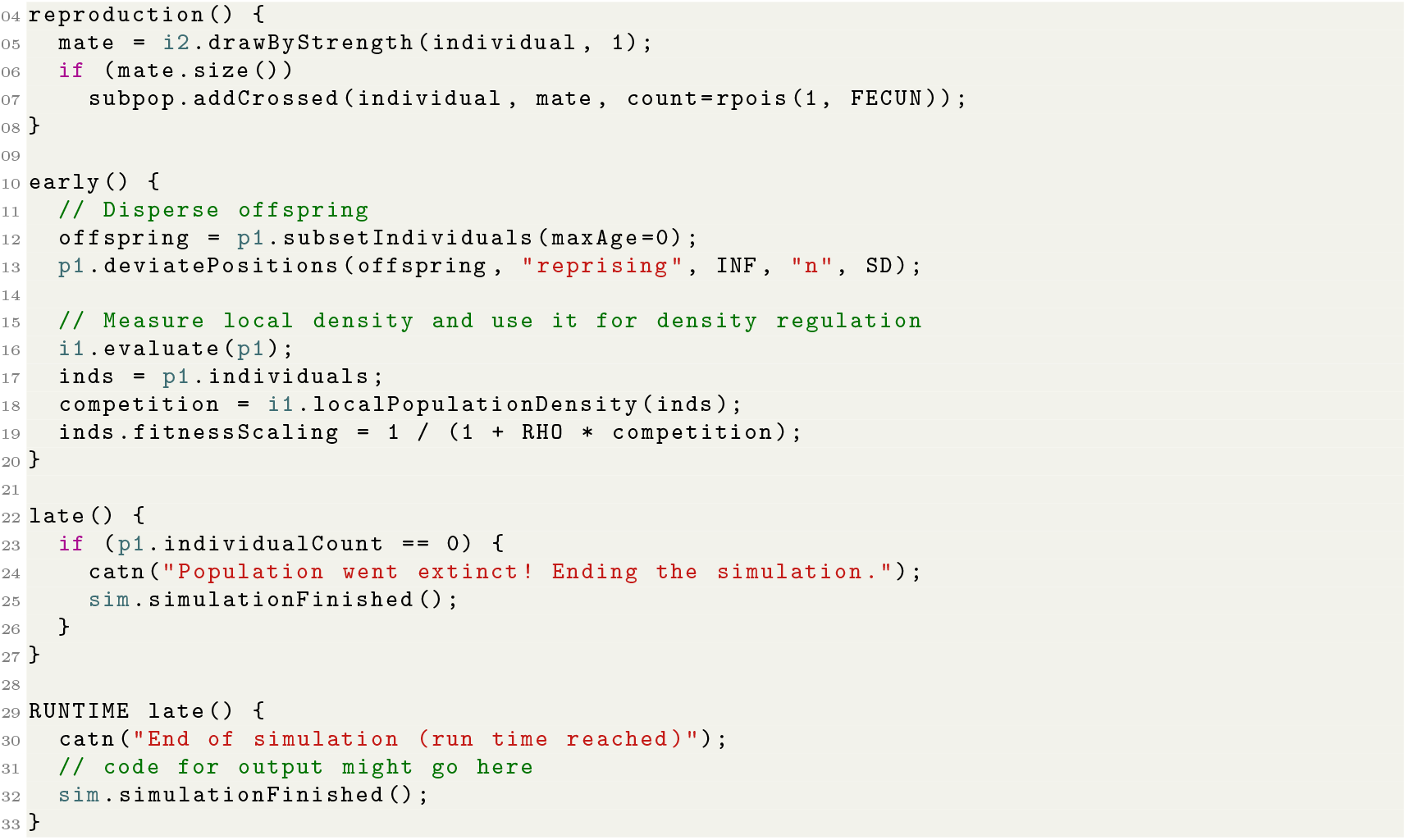

## Appendix B Pitfalls

Evenafter carefully parameterizing a simulation to equilibrate near a given population density, it is fairly easy in practice to end up with a spatial simulation that mysteriously dies out or behaves oddly in other ways. A less dramatic annoyance is that usually the realized population size is not equal to the desired density, *K*, multiplied by the total area. Indeed, developing a formula for expected total population size in terms of the simulation parameters that is better than a rough first-order approximation seems extremely difficult. This section describes the root causes of these issues and ways to diagnose them. The discussion gets into the weeds, so here is a summary of what to check if density is not what you expect (details below):

1. Visualize the simulation to check for odd dynamics or spatial patterns, such as a regular array of clumps.
2. Make sure the neighborhood sizes *N*_*X*_, *N*_*M*_, and *N*_*D*_ are not too small (if in doubt, observe the effects of increasing *σ*_*X*_, *σ*_*M*_ and/or *σ*_*D*_).
3. Look at the mean density *experienced by individuals*, not the total density across the landscape. If you want instead to set the total population size, you’ll need a post-hoc adjustment to *K* as in Box 6.
4. Make sure the *stage* you’re measuring density in agrees with the theoretical calculations (*i.e*., between birth and death or between death and birth).
5. Consider stochasticity: density varies randomly across the landscape, making the realized mean density differ from *K*.

Although it is natural to expect that the realized density of a simulation will be exactly the specified value of *K*, it is important to remember in practice that having a density different from *K* is not necessarily a problem: instead, it may reflect the natural biological consequences of the chosen model.

### B.1 Why are there weird regular clumps?

At its most extreme, a dispersal scale much smaller than the interaction scale can lead to strange, regular arrays of clumps. (Clumps may appear for many other reasons, but here we’re talking about a regular, hexagonal grid of clumps.) Examples are shown in Figure S8. Although such regular patterns formed by this mechanism rare in nature, they are easy to accidentally produce in simulation (and are one reason it is important to visualize the simulation, as in Box 2). For discussion of this strange phenomenon, see Sasaki (1997), Etheridge et al. (2024), or the “Spatial competition and spatial mate choice in a nonWF model” section of the SLiM manual (Haller and Messer, 2024). These are probably an indication that the dispersal or interaction scale are not well-chosen, but may indicate something more interesting.

### B.2 Why does my simulation run so slowly?

The runtime of an individual-based simulation is at least proportional to the total number of individuals. However, it is common for the runtime of spatial simulations to grow *more* than linearly in the number of individuals, because of spatial dynamics that involve a large number of pairwise comparisons or interactions (such as spatial mate choice and spatial competition). Performance problems resulting from this can often be diagnosed by looking for *large* neighborhood sizes: if *N*_*X*_ (the interaction neighborhood size) or *N*_*M*_ (the mating neighborhood size) are large, one may encounter slow run times. Happily, there are solutions that can often be applied.

Perhaps the most obvious solutions are to directly reduce the number of pairwise interactions. One way is to shrink the neighborhood sizes of the model, by reducing *σ*_*M*_ and/or *σ*_*X*_ . However, that will often noticeably change the behavior of the model and sacrifice biological realism. Another way is to shrink the neighborhood size is to cut off the spatial kernel at a shorter distance, with little loss of exactitude; for a Gaussian spatial kernel, for example, cutting off at two standard deviations rather than three can reduce runtime with (perhaps) little change in dynamics, since interacting individuals 2–3 standard deviations from the focal individual interacted with that individual quite weakly anyway. However, this cannot cut runtime by more than about half, so for most models with large neighborhood sizes, another strategy is needed.

A second option is to use a “resource node” approach, as demonstrated in the monarchs example (Section 9.4), which effectively mediates the many possible individual-individual interactions with a smaller number of interactions between each individual and a nearby node (representing a localized amount of resources). The approach is discussed more fully in Champer et al. (2024).

A third option is to use spatial map operations to approximate the pairwise interactions more efficiently. Effectively, a map of the population’s density can be computed in each tick of the model and then used to look up the density near each individual as a summary of all of the pairwise interactions it receives (Box 8). It turns out the same method can be used to efficiently pick nearby mates as well (Box 9). This option is in fact equivalent to using a regularly tiled grid of resource nodes, and can be proven to be a good approximation: see Appendix F.

Figure S1 shows that using the map-based approximation methods described in Boxes 8 and 9 makes it much easier to scale simulations to much higher neighborhood sizes. A naive implementation of pairwise interactions would result in runtimes that are quadratic in total population size, and hence totally infeasible for all but very small populations. Standard pairwise interactions in SLiM use efficient data structures (*k*-d trees) and a maximum distance cutoff (here, of 3*σ*), but still compute all pairwise interactions out to the maximum distance, and so are quadratic instead in neighborhood size (*N*_*X*_ or *N*_*M*_), shown as solid lines in Figure S1. Models using spatial map-based methods or a regularly tiled grid of resource nodes, on the other hand, scale linearly with neighborhood size. These faster methods are approximate, but correspond closely, especially at high densities, as shown in Figure S2.

Each option has pros and cons, and may alter the behavior of the model. If the natural dynamics of the species are mediated through discrete locations (e.g., feeding or locations or mating sites), then adapting the resource node method is probably the most natural method. (In fact, it has been suggested that uncommon insects gather in discrete locations to find mates for not dissimilar reasons (Alcock, 1987).) If not, it may be more natural to use map-based methods. Finally, it may be necessary (especially for development purposes!) to simply model a smaller landscape.

**Figure S1:**
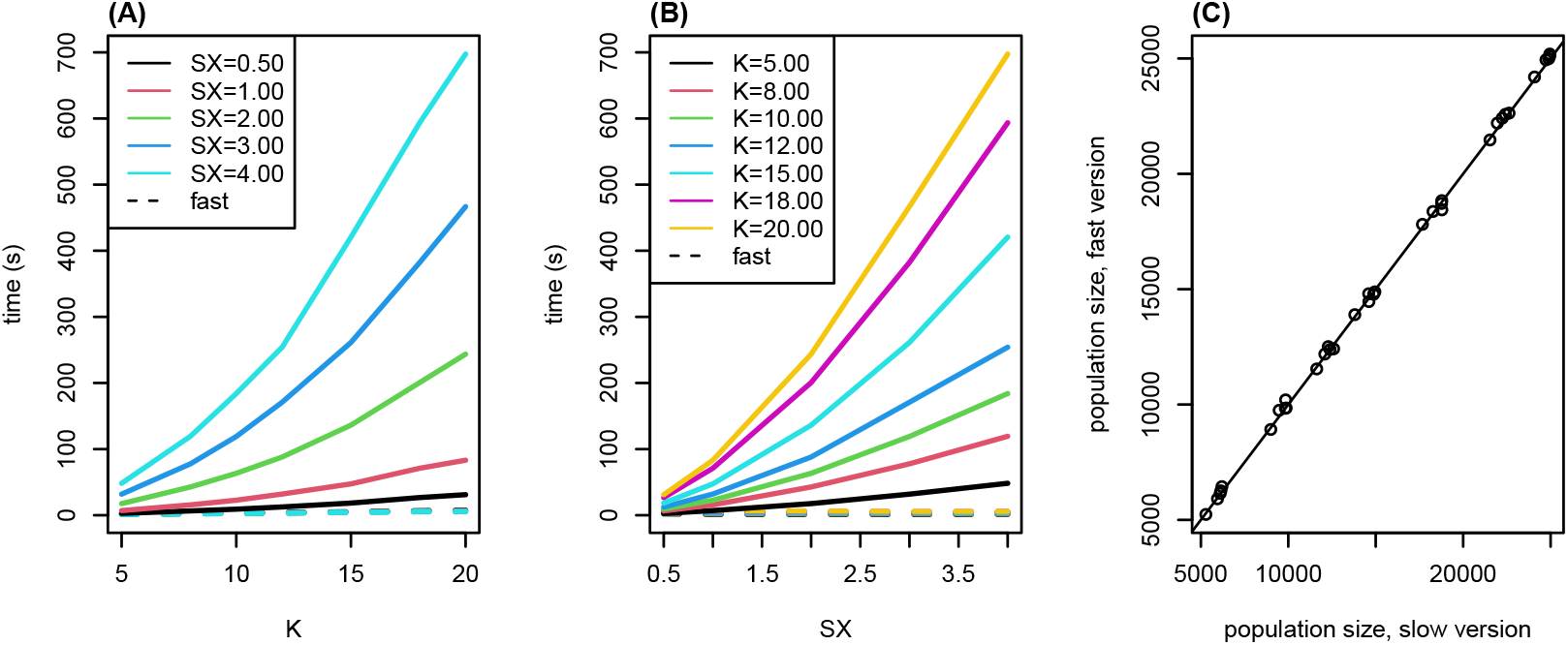
Runtimes for a “minimal” model with local mate choice and density-dependent Beverton–Holt feedback on mortality, plotted against **(A)** carrying capacity (*K*) and **(B)** interaction scale (*σ*_*X*_, written `SX`). Runtimes are shown for models that do both mate choice and local density computations (solid lines) using (pairwise) interactions, and (dotted lines, all overlapping) using spatial maps, as described in Boxes 8 and 9. Also, **(C)** final population sizes after 100 time steps for the same combinations of *K* and *σ*_*X*_ shown in (A) and (B), for otherwise equivalent models that use either pairwise interactions (horizontal axis, “slow version”), or spatial map methods (vertical axis, “fast version”).

### B.3 Why does my simulation die out?

There are a variety of reasons why a simulation might die out (or have far fewer individuals than you expect). For instance, this can happen if *σ*_*M*_, *σ*_*X*_, or *σ*_*D*_ are too small. In all cases, “too small” can be diagnosed by looking at the relevant neighborhood sizes: for example, if 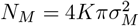 is small (less than about 1), there may be a problem related to *σ*_*M*_ . This problem manifests as individuals being unable to reproduce because they cannot find a mate. This is particularly likely to happen if the mating scale has been made smaller to reduce runtime (see the previous section for discussion). Solutions might be either to increase *σ*_*M*_, to allow selfing, or to increase the fecundity of those individuals that do reproduce, depending upon the biology of the system being modeled. The result also depends on the mating scheme, as shown in Figure S6.

The reasons that small *σ*_*X*_ can be a problem are more subtle. Since *σ*_*X*_ determines the range over which density is computed, and each focal individual itself counts towards its local population density, then if *σ*_*X*_ is sufficiently small even a single isolated individual can have “local density” greater than the carrying capacity. This effect is demonstrated in Figure S7, in which the population dies out for small *σ*_*X*_ . The effect also appears in Figure S6, in which selfing simulations die out at low *K* (and hence low *N*_*X*_) – since they self, they are not dying out due to small *N*_*M*_ . Conceptually, this happens if the simulated individuals cannot range over a large enough area to obtain sufficient resources for survival, even in the absence of competition.

Since the density of a single individual calculated by equation (1) is 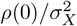,then this will occur if 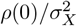 is close to or greater than *K, i.e*., roughly if 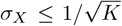.This could be avoided by excluding the focal individual from the calculation of local density; however, this leads to the opposite problem: densities can get much larger than *K*. This happens because if 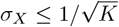,a single neighbor within scale *σ*_*X*_ will lead to a “local density” of more than *K* – but further away neighbors are unaffected. If local density increases mortality, a cartoon version of the situation is that an individual with a neighbor within distance *σ*_*X*_ is killed, but neighbors further away than this are ignored; as a result, the density equilibrates to around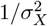, rather than *K*. (The above discussion is in two dimensions; in a one-dimensional model the equilibrium density would be around 1*/σ*_*X*_ .)

**Figure S2:**
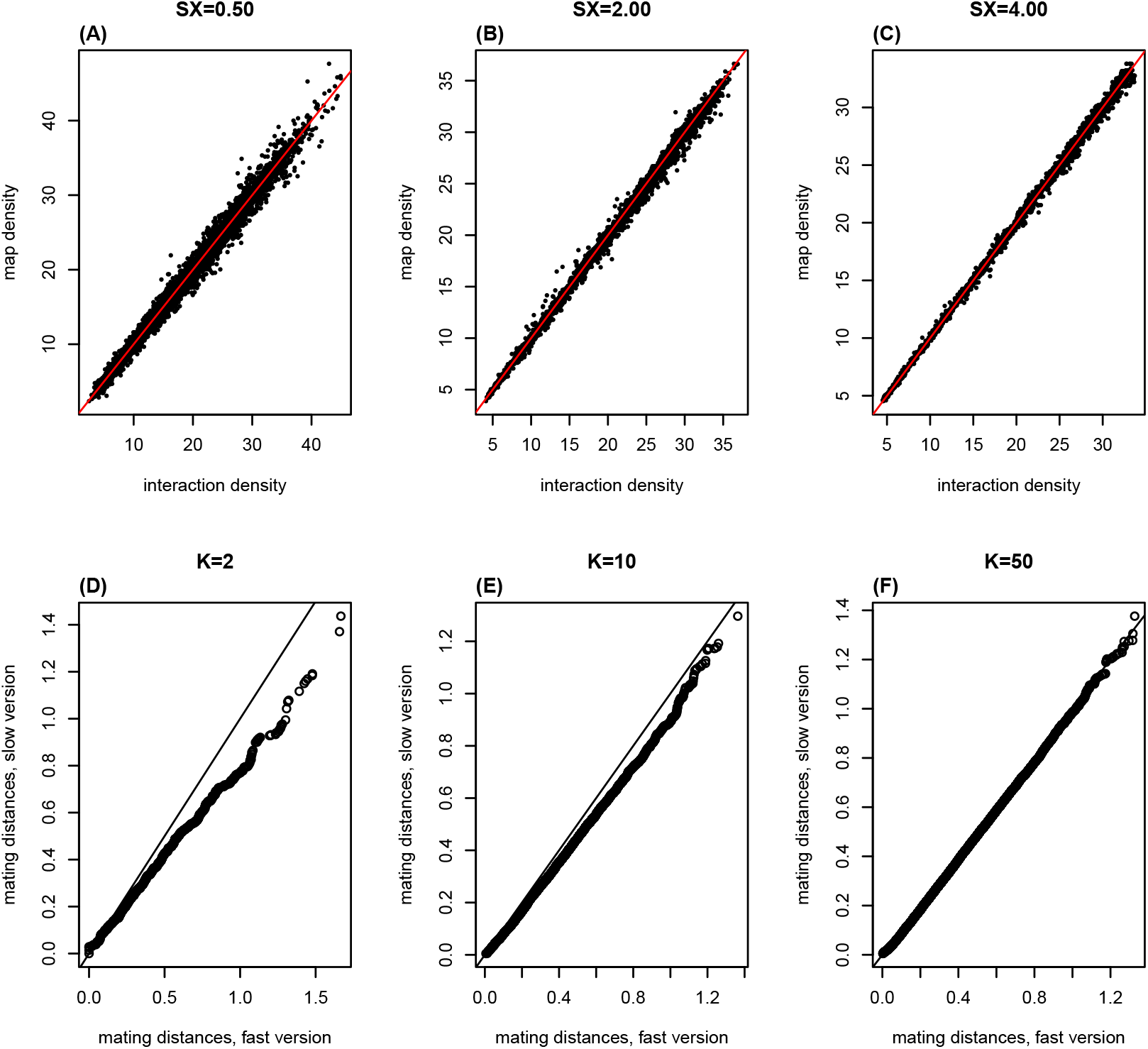
**(A-C)** Local density around each of 10^4^ individuals, measured both using pairwise interactions (equation (1)) and interpolation on a spatial map (Box 8). Each panel shows 10^4^ individuals randomly sampled from a separate simulation of the type described in Figure S1, but with *K* increasing linearly over 20 time steps, and different values of *σ*_*X*_ (labeled `SX`). **(D-F)** Q-Q plots comparing realized distributions of mating distances between simulations as above but either using individual-based mate choice (horizontal axis, “slow version”) or using map-based mate choice as in Box 9 (vertical axis, “fast version”), at three values of density (*K*). Shown are the quantiles of the distances to the most recent mate for roughly 30,000 individuals in each simulation; at lower densities, the “fast version” tends to have slightly longer mating distances.

The population can die out if *σ*_*D*_ (and/or *σ*_*V*_) is too small for similar reasons: conceptually, if offspring do not disperse far enough from their parents and local competition is strong, then families reduce their own fitness. For a simple example, suppose that the probability that a new offspring with local density *n* survives is *e*^−*n/K*^ and *σ*_*D*_ is much less than *σ*_*X*_ . Then, a group of *m* siblings form a clump of *m* + 1 individuals with their parent, with density at least 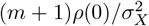; so, the expected number of surviving offspring is smaller – at most 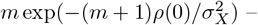 and the smaller number should be used in calculations of net reproductive output. However, if new individuals move sufficiently far (with *σ*_*V*_) before local density effects, then the effect may be avoided.

The effects of changing *σ*_*X*_, *σ*_*D*_, and removing the focal individual from density calculations are shown in an example in Figure S9.

### B.4 Why is the realized density not equal to *K*?

We’ve carefully set things up so that the equilibrium density in a neutral, spatially homogeneous simulation “should be” equal to *K*. If the realized density is very different (*e.g*., zero) then the problem is probably one of the pitfalls described above. But even if *N*_*X*_, *N*_*M*_, and *N*_*V*_ are not small, realized density still often differs from *K* by 20% or 30%. Perhaps the simplest reason is “edge effects,” but we assume the range is large enough these are unimportant (and, in practice SLiM computes local population density in such a way that local density is unaffected by edges). Another simple reason could be that density differs at different points in the life cycle – see below for more discussion of this.

First, we need to consider: *which* realized population density should we compare to *K*? The first answer that might spring to mind is “number of individuals divided by total area;” however, what matters for equilibrium is the density *experienced by individuals*. In other words, to see theoretical predictions playing out, we should measure local population density for each individual, and average that across individuals – that is after all the density that matters to the dynamics. This is seen in Figure S7, where mean density around individuals is shown in blue and number of individuals divided by area is shown in red, as well as in Figure S6, where mean density around individuals is shown in dotted lines and number of individuals divided by area in solid. Conceptually, if at equilibrium the simulation is very patchy (so individuals tend to be bunched up), then the mean density experienced by individuals could be much higher than the number of individuals divided by total area. In fact, the mean density around individuals is almost always be higher than the number of individuals divided by total area.

This line of reasoning leads to the second point: across individuals, local population density is a distribution, not a single value. It turns out that this stochasticity can also be important. Conceptually, the decrease in net reproduction of individuals with higher than average density may not be balanced by those individuals with lower density; how this happens depends on the shape of *F* (*u*) and the shape of the distribution of densities. Below, we work through both these issues in more detail.

#### B.4.1 Mean density around individuals

Why is the density experienced by individuals higher than the number of individuals divided by total area? Concretely, suppose that *u*_*i*_ = *n*(*x*_*i*_)*/K* is the scaled local density for individual *i*; the expected change in population size is zero if ∑ _*i*_ *F* (*u*_*i*_) = 0. Equivalently, if *U* is the scaled local density for a randomly chosen individual, equilibrium occurs if 𝔼[*F* (*U*)] = 0. Suppose instead that we look at the distribution of local densities across *space* instead of across individuals. Heuristically, suppose that we divide the landscape up into many small regions, each of area *ϵ*, and let *v*_*j*_ be the scaled density in region *j* (*i.e*., *n*(*y*_*j*_)*/K* for some point *y*_*j*_ in region *j*, and the regions are small enough that density is constant within each). The number of individuals in region *j* is *ϵKv*_*j*_, and so the net contribution to the next step’s population size from region *j* is *ϵKv*_*j*_*F* (*v*_*j*_). This equilibrium occurs if ∑ _*j*_ *v*_*j*_*F* (*v*_*j*_) = 0. Equivalently, if we let *V* denote the local density around a uniformly chosen point on the landscape, then equilibrium occurs if 𝔼[*V F* (*V*)] = 0. In fact, the relationship between *U* and *V* is that *U* is a *size-biased* draw from *V* ; the relationship between the two is that 𝔼[*f* (*U*)] = 𝔼[*V f* (*V*)]*/*𝔼[*V*] for *any* function *f* . In particular, 𝔼[*U*] = 𝔼[*V* ^2^]*/*𝔼[*V*] *>* 𝔼[*V*] (by Jensen’s inequality), and 𝔼[*V*] is just the total number of individuals divided by the total area (except for some edge effects).

Sharp-eyed readers of this and the next section will notice that we are sweeping something under the rug: when we measure density using equation (1), we do not include the focal individual. Taking this into account properly when defining *V* is much less clean, so for illustrative purposes we have omitted this. In fact, if individual locations are independent and uniformly distributed, then mean density around individuals (measured without the focal individual!) is equal to the number of individuals divided by total area. Nonetheless, we think the calculations are informative.

#### B.4.2 Stochasticity

Now we can immediately see how stochasticity interacts with nonlinearity in density dependence to increase or decrease equilibrium density. First suppose that *F* is convex and decreasing, *i.e*., *F* ^*″*^ (*u*) *>* 0 and *F* ^*′*^ (*u*) ≤ 0 for all *u*. Then by Jensen’s inequality, 𝔼[*F* (*U*)] *> F* (𝔼[*U*]), and since at equilibrium, 𝔼[*F* (*U*)] = 0, we have that *F* (𝔼[*U*]) *<* 0. Since we’ve assumed that *F* (1) = 0 and *F* is decreasing, this implies that 𝔼[*U*] *>* 1, *i.e*., a convex *F increases* the equilibrium mean density experienced by individuals above *K*. By the same argument if *F* is concave, 𝔼[*U*] *<* 1.

We can make the same argument for the total density, *V* : if *G*(*v*) = *vF* (*v*) is convex, then 𝔼[*V*] *>* 1, while if *G* is concave then 𝔼[*V*] *<* 1. This at first seems odd: if *F* (*u*) is convex and *uF* (*u*) is concave, then the mean density experienced by individuals is *higher* than *K*, while the total density is *lower* than *K*. However, this is perfectly possible, and in fact seen in Figure S7.

A Taylor expansion lets us estimate more precisely the deviation of equilibrium size from *K*. Taylor expanding *F* (*u*) about *u* = 1, we get that

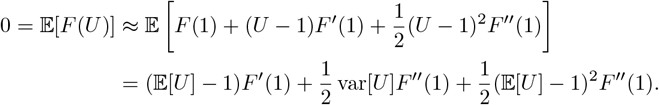

If the deviation is small (*i.e*., |𝔼[*U*] −1« |*F* ^*′*^ (*U*)*/F* ^*″*^ (*U*) |, (𝔼[*U*] −1)^2^ «var[*U*]), the second order term is negligible. Thus, we can write:

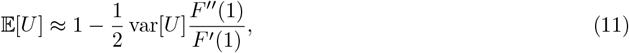

when 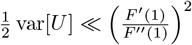,Since *F* ^*′*^ (1) is negative, agrees with the argument above. Figure S10 shows that this prediction bears out well in practice (in two nonspatial models), as long as the population does not go extinct.

Similarly,

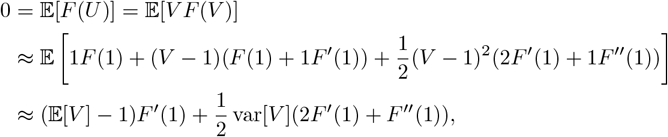

and hence

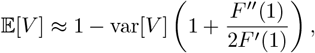

which is smaller than 𝔼[*U*], and further away from 1, except in extreme circumstances.

What determines the strength of stochasticity? Since this has to do with random variation in “experienced” density across the landscape, stochasticity goes down as interaction neighborhood size *N*_*X*_ increases. In other words, if *σ*_*X*_ is larger, then we measure density averaging over larger areas, which is therefore less variable. As in equation (11), stochasticity affects equilibrium by a factor proportional to var[*U*]. If *Y* is the number of individuals within distance *σ*_*X*_ of a random individual, then 𝔼[*Y*] is around *N*_*X*_, and if noise is

Poisson then also var[*Y*] *≈ N*_*X*_ . Since *U* is obtained by dividing *n*(*x*) (from equation (1)) by *K*, and *n*(*x*) is roughly 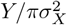,we expect var[*U*] to be of order 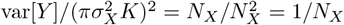.So, we expect the deviation of realized density from *K* to be of order 1*/N*_*X*_ . Again, this is seen in Figure S7: the form of density dependence has a convex *F* (*u*), and so for smaller values of *σ*_*X*_, the value of 𝔼[*U*] (blue line) is above *K*. Conversely, the function *uF* (*u*) is concave, and so 𝔼[*V*] (red line) is increasing, but is well below *K* for other reasons.

The difference between the mean density around individuals (𝔼[*U*]) and the mean density by area (𝔼[*V*], or total population size divided by total area) is well-known: the ratio 𝔼[*U*]*/*𝔼[*V*] is a scale-dependent measure of clustering known as *mean crowding* (Lloyd, 1967) that increases the more clustered individuals are on the scale used to measure local density.

#### B.4.3 Density measurement timing

*The other thing to consider is: when* is density being measured? In each time step there are some births and some deaths; we follow SLiM in taking births first in the time step, but since the two alternate, this choice seems arbitrary. However, having density effects or movement occur in one or the other stage can affect the model (e.g., Taylor, 2010). Following SLiM as we do, the most common “population size” is at the end of the time step, *i.e*., after deaths (or equivalently, before births). However, population size may also be measured (and used!) between births and deaths.

Concretely, suppose that *N*_*t*_ is the population size at the start of time step *t*, and 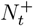 is the population size after births in time step *t*. Above, we did calculations like this: if the mean fecundity in time step *t* is *f*_*t*_, and the mean probability of survival is 1 − *µ*_*t*_, then

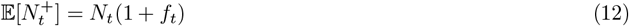

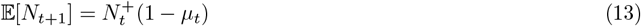

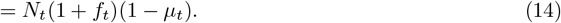

*Which* density is used to determine *f*_*t*_ and *µ*_*t*_? Naturally, *f*_*t*_ can’t depend on the number of births, so it will use *N*_*t*_ (*i.e*., the density at the start of the time step). However, should the density dependence for *µ*_*t*_ use *N*_*t*_ or 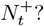 If the newborn individuals contribute to density, then *µ*_*t*_ should depend on 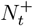, *i.e*., density computed using the offspring as well. However, this will be larger than the density at the start of the time step by a factor of 1 + *f*, and so the equation for the local net per capita reproductive rate analogous to equation (2) is

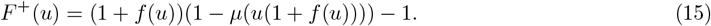

As before, equilibrium would be around density *n*_*_ solving *F* ^+^(*n*_*_*/K*) = 0, and so to arrange as before for the equilibrium density to be around *K* we’d like to set up the functional forms so that *F* ^+^(1) = 0. One way to do this is to start with functional forms for *f* (*u*) and *µ*(*u*) so that *F* (*u*) = (1 + *f* (*u*))(1 −*µ*(*u*)) − 1 satisfies *F* (1) = 0 (*i.e*., formulated for measuring density between death and birth), and then define the survival probability to be 1 −*µ*^+^(*u*) = 1 −*µ*(*u/*(1 + *f* (1))). Then *F* ^+^(*u*) defined using *µ*^+^(*u*) satisfies *F* ^+^(1). This is the approach taken by Battey et al. (2020). Another approach is to compute a density *map*, as in Box 8, at the start of each time step, and use that map to determine density for juveniles as well.

## Appendix C Parameterization of density dependence

If one wants to use a familiar phenomenological model as the basis for density dependence, there are several popular choices for the function form of *F* (*u*). Here, we summarize these, parameterized so that *F* (1) = 0 (and so will have an equilibrium near *n*_*_ = *K*):

- *F* (*n*) ∝ 1 − *n*, (Discrete-logistic model)
- 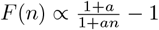, (Beverton–Holt model)
- 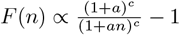,Hassell model)
- *F* (*n*) ∝ *e*^*r*(1−*n*)^ − 1. (Ricker model)

(Here, ∝ indicates that each can be scaled by a constant, reflecting an overall time scaling.)

Using each one still requires a number of choices. Next, we work through in more detail the steps involved in arranging birth and death rates so that the net birth rate, *F*, has a particular functional form, and give a number of examples that help to show the issues involved. Suppose here that each time step has birth followed by death; death applies in the same way to individuals just born as those previously alive; the mean fecundity of an individual with local density *N* is *f* (*N/K*); and the probability of death of an individual with local density *N* is *µ*(*N/K*).

Roughly, the net change in population size when at scaled density *u* = *N/K* is *F* (*u*) = *f* (*u*)(1− *µ*(*u*)) −*µ*(*u*). However, this is not not necessarily right, since it depends when the densities are measured: if the density for mortality is measured after birth, then the density passed to *µ* will be different than that passed to *f* . So, we’ll always define *F* (*u*) to be the mean per-capita change in population size across one step when starting at scaled population size *u*. However, which point in the time step (*i.e*., after birth and before death or vice-versa) is the reference point will depend on the situation. Our goal is to figure out how to arrive at a given functional form for *F* in various scenarios.

### Fecundity regulation

If death probability is constant: *µ*(*u*) = *µ*_0_, then taking *u* to be the scaled population density before birth, simply

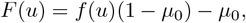

and so

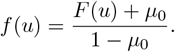

Since 0 ≤ *f <* ∞, for this to make sense we need *µ*_0_ *>* 0 and *F ≥* −*µ*_0_.

### Beverton–Holt fecundity regulation

With *F* (*u*) = *α*((1 + *a*)*/*(1 + *an*) − 1), this is

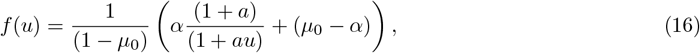

and we need *α* ≤ *µ*_0_.

### Ricker fecundity regulation

With *F* (*u*) = *α*(*e*^*r*(1−*u*)^ − 1), this is

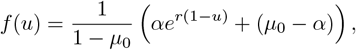

and we again need *α* ≤ *µ*_0_.

### Mortality regulation

Suppose instead that fecundity is constant: *f* (*u*) = *f*_0_, and that we measure density for mortality *after* birth (so it includes the new births). Then

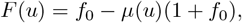

and so the survival probability is

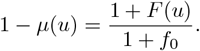

Since we must have 0 ≤ *µ* ≤ 1, we require that −1 ≤ *F* ≤ *f*_0_. Note that in this model (regardless of the form of *F*), the death probability at the stationary point (*u* = 1, since *F* (1) = 0) is *µ*(1) = 1*/*(1 + *f*_0_); and so the mean lifetime is 1 + *f*_0_. Also note that this produces an equilibrium density of *K before death* (as opposed to before birth, in the previous models); the density will be lower after death.

### Beverton–Holt mortality regulation

With *F* (*u*) = *α*((1 + *a*)*/*(1 + *au*) − 1), this is

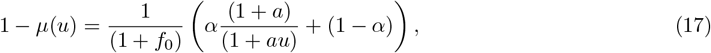

and we need *αa* ≤ *f*_0_.

### Ricker mortality regulation

With *F* (*u*) = *α*(*e*^*r*(1−*u*)^ − 1), this is

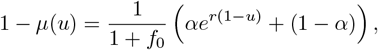

and we need *α*(*e*^*r*^ − 1) ≤ *f*_0_.

### Mortality and fecundity

Suppose now we’d like both mortality and fecundity to change with density. So that there’s only one density, let’s say again that is measured after birth and before death. If the density at this time is *N*, then the mean number of individuals in the next time step is

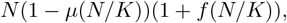

so that the mean net change is

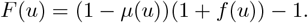

Given a desired functional form for *F* (*u*) and *µ*(*u*) we would then define

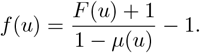

Note that for fecundity to remain finite, we need *µ*(*u*) to be bounded away from zero. On the other hand, if we have *f* (*u*) then

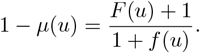

### Mixed Beverton–Holt

Suppose we set *f* (*u*) = *f*_0_*/*(1 + *bu*) and would like *F* (*u*) = *α*((1 + *a*)*/*(1 + *au*) − 1) for some constants *b* and *α*. Then, we would set

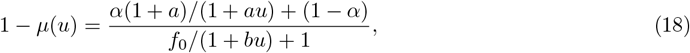

and we need *αa < f*_0_.

### Mixed Ricker

Now suppose that we set 1 − *µ*(*u*) = (1 − *µ*_0_)*e*^−*su*^ + *µ*_∞_(1 − *e*^−*su*^) and would like *F* (*u*) = *α*(*e*^*r*(1−*u*)^ − 1). (The extra parameters are not unnecessary complications: we will need *α <* 1 and *µ*_∞_ *>* 0 for the following to work out.) Then, we would set

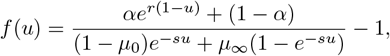

and we need *s* ≤ *r* for this to remain positive. This has *f* (0) = (*α*(*e*^*r*^ − 1) + 1)*/*(1 − *µ*_0_) − 1, which is positive if *r >* 0 (already a requirement). Also, *f* (∞) = (1 − *α*)*/µ*_∞_ − 1, so we also need 1 − *α > µ*_∞_.

### Hassell with mortality regulation

Let’s set up the Hassell, which is *F* (*u*) = *b*((1 + *a*)*/*(1 + *au*))^*c*^ − 1). Note that *F* (0) = *b*((1 + *a*)^*c*^ − 1), *F* (∞) = −*b*, and *F* ^*′*^(1) = *cab*(1 + *a*)^*c*^. Setting fecundity to be constant at *f*_0_ and plugging in to the expression for mortality regulation above,

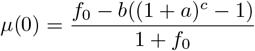

and

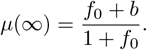

So, if we’d like to fix *µ*(0) = *µ*_0_ and *µ*_∞_ = *µ*(∞), then these determine *a* and *b*:

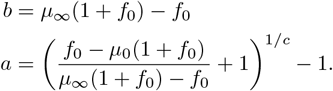

This leaves us with a parameterization in terms of the mean fecundity, *f*_0_, the death rate at low density, *µ*_0_, the death rate at high density, *µ*_∞_, and the exponent *ϵ* that controls how steep the curve is between. (We could reparameterize *ϵ* so we have a parameter that is exactly *F* ^*′*^(1), but this is less compelling.)

### Ricker parameterization with fecundity regulation

If *µ*(*u*) = *µ*_0_ and we want *F* (*u*) = *C*(*e*^*r*(1−*u*)^ − 1), then

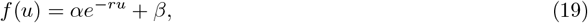

where letting *α* = *Ce*^*r*^*/*(1 − *µ*_0_) and *β* = (*µ*_0_ − *C*)*/*(1 − *µ*_0_). So, *f* (0) = *α* + *β* and *f* (∞) = *β*. If we set *C* = *µ*_0_ then this is *β* = 0 and *α* = *µ*_0_*e*^*r*^*/*(1 − *µ*_0_), and so *f* (*u*) = *µ*_0_*e*^−*r*(*u*−1)^*/*(1 − *µ*_0_).

### Regulation by juvenile mortality

Now suppose that adult death rate and fecundity are constant (so, *f* (*u*) = *f* and *µ*(*u*) = *µ*), but that the probability of survival of *juveniles* is density-dependent. (Perhaps density dependence only affects the species during seedling recruitment.) So, if we call *r*(*u*) the probability of survival in the first year at scaled density *u*, the net per capita reproduction function is *F* (*u*) = *r*(*u*)*f − µ*. To make this proportional to the Beverton–Holt form, we can set

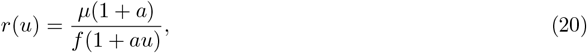

which results in *F* (*u*) = *µ*((1 + *a*)*/*(1 + *au*) − 1) – *i.e*., the Beverton–Holt form, scaled by *µ*.

### Density regulation of both juvenile and adult mortality

Now suppose that the probability of survival of juveniles (*i.e*., to their first year) is *r*(*n/K*), not 1 − *µ*(*n/K*). In this case,

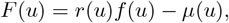

and so if we fix *f* (*u*) = *f* then we have, for instance,

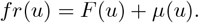

Suppose we want *F* (*u*) = (1 + *a*)*/*(1 + *au*) − 1 and *µ*(*u*) = *bu/*(1 + *bu*); then we would have

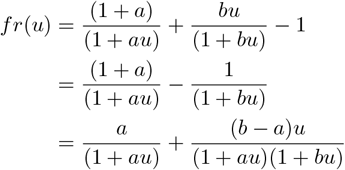

With this definition, *r*(0) = *a/f* and *r*(∞) = (*b* −*a*)*/*((*b* + *a*)*f*), so we need *a* ≤max(*b, f*) (and some other conditions). Note that *r*(∞) *> r*(0) (*i.e*., *increasing* recruitment with density) if *b* −*a > a*(*a* + *b*): for instance, take *f* = 1 and *a* = 1*/*4 and *b* = 3*/*4.

## Appendix D Parameterization and sampling for dispersal kernels

When sampling a new random vector for displacement or dispersal, it is simplest to think in Cartesian coordinates: to draw the displacement as (*X, Y*) where *X* and *Y* are independent draws from some distribution. This works well if *X* and *Y* are Gaussian, but plotting the resulting bivariate distribution quickly shows oddities: dispersers will tend to fall on either around the axes or around the diagonals, depending on the distribution chosen. In fact, displacements in orthogonal directions are independent *only* for the Gaussian distribution. To obtain a rotationally symmetric dispersal kernel, it helps to think in polar/spherical coordinates. This also brings up an issue of terminology: what do we call a given rotationally symmetric bivariate distribution? Natural choices are to name it after the shape of either (a) the profile, *X*, or (b) the distance, 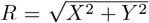.These agree only in one dimension. We work through some examples below to make the underlying issues clear.

### Gaussian (and Rayleigh)

Imagine displacements along the *x* and *y* axes are sampled from the same Normal distribution with zero mean and variance of *σ*^2^. The displacement in *x, y* then has the multivariate normal density

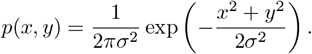

In polar coordinates,

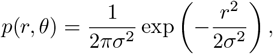

*i.e*., the density only depends on *r*, and so the kernel is radially symmetric.

To find the distribution of *R*, we can find the cumulative distribution, ℙ(*r < R*), and differentiate. The CDF can be found by integrating *p*(*x, y*) over rings of circumference 2*πr* and infinitesimal width of *dr*:

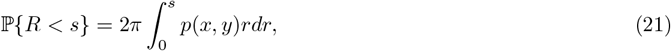

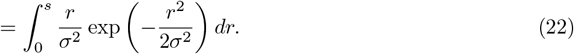

Thus, the PDF of 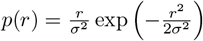 which is a Rayleigh distribution. In summary,

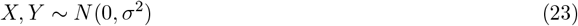

is equivalent to

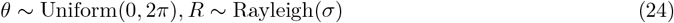

By the same argument, in three dimensions the angular part would be uniformly distributed on the sphere, and *R* has density proportional to *r*^2^ exp(*r*^2^*/*2*σ*^2^).

### Student’s *t*

We can do something similar with the Student’s *t* distribution, but it will be clear that we need to be careful with generalizing it to a higher dimension.

First, what is the Student’s *t* distribution? In one dimension, it is the distribution with density

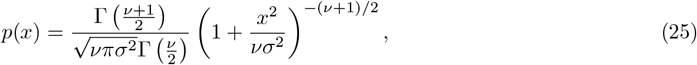

where ν is the “degrees of freedom” and *σ* is a scale parameter. If ν = 1, it is a Cauchy distribution, and if ν → ∞, the distribution converges to a standard Normal distribution. The Student’s *t* distribution is a heavy tailed distribution: all moments of order ν or higher are not defined.

Now suppose we want to “use the Student’s *t*” in two dimensions. A **wrong** way to do this is to sample *x* and *y* independently from the same *t* distribution. If you do that, the joint distribution *p*(*x, y*) is *p*(*x*)*p*(*y*) due to independence, and it looks like

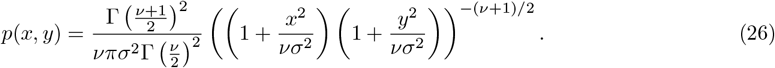

If we plug in *x* = *r* cos *θ* and *y* = *r* sin *θ* as we did before with the Normal distribution, we realize that *p*(*x, y*) depend on both *r* and *θ*! In other words, the distribution of *R* depends on the angle, *θ*: long distances will be more common in some directions than others.

It turns out that although there are many ways to generalize the *t* distribution to more than one dimension, there is not a single standard way (Kotz and Nadarajah, 2004). Here are three possibilities for how we might choose a radially symmetric bivariate kernel *p*(*x, y*):

1. The distribution of the distance, *R*, is Student’s *t*.
2. The distribution of the distance along an arbitrary axis, *X*, is Student’s *t*.
3. The shape of the kernel taken along a line through the origin is the Student’s *t* density.

These conditions are, equivalently, that (1) *p*(*r, θ*) ∝ *r*^−1^(1 + *r*^2^)^−(ν+1)*/*2^ (where *p*(*r, θ*) is *p*(*x, y*) is radial coordinates); (2) ∫ *p*(*x, y*)*dy* ∝ (1 + *x*^2^)^−(ν+1)*/*2^; and (3) *p*(*x*, 0) ∝ (1 + *x*^2^)^−(ν+1)*/*2^. If we chose the first option, then we’d be compelled to call the bivariate Gaussian the “bivariate Rayleigh” distribution, so for consistency with the Gaussian, we’ll discard that option. Although option (2) is perhaps more elegant, figuring out what the actual density is for an arbitrary kernel shape is more involved, so we have chosen to go with option (3) (both here and in SLiM).

So: to make a radially symmetric Student’s *t* kernel, we’d like a bivariate kernel *p*(*x, y*) proportional to 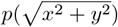,where the second *p*() is the univariate Student’s *t*. With the normalization factor, for ν *>* 1 this is

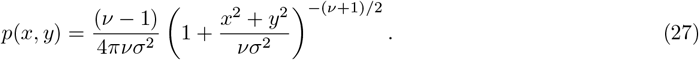

The density of *R*, the displacement distance, is equal to 2*πrp*(*r*, 0) .

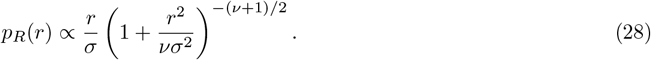

Note that if ν = 1 (*i.e*., the Cauchy distribution), the integral 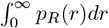 does not converge. This means ν = 1 gives an invalid probability distribution, and so in two dimensional space, the number of degrees of freedom should be greater than 1. (More generally, in *d* dimensions we’ll need ν *> d* − 1.)

### General kernels

To generalize this, begin with a univariate probability distribution with density *f* (*x*). Then, we define another distribution for *r* by *p*_*R*_(*r*) ∝ *r*^*d*−1^*f* (*r*) if in *d* dimensions. This is equivalent in two dimensions to defining a joint distribution of *x, y* by 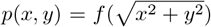. To sample a random variable (*X, Y*) in two dimension, we sample *R* from *p*_*R*_(*r*), sample *θ* from the uniform distribution on (0, 2*π*), and set *X* = *R* cos *θ* and *Y* = *R* sin *θ*. (More generally, we would choose the radial component to be uniform on the *d*-sphere; the easiest way to do that is to let 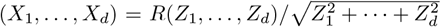,where *Z*_1_, …, *Z*_*d*_ are independent standard Normal.) This is how the function `pointDeviated()` in SLiM returns a displaced locations from various rotationally symmetric kernel.

#### D.1 Sampling from kernels with a covariance matrix

The multivariate Normal is not necessarily rotationally symmetric: it allows a general covariance matrix; if the covariance matrix is not a multiple of the identity, then the contours of the density are ellipses, not circles. How might we introduce covariance matrices to multivariate dispersal? The most convenient way to do this is to use *scale mixtures* of Normals, *i.e*., just multiply a given multivariate Normal distribution (with some covariance matrix) by a random scaling, choosing the distribution of the random scaling appropriately.

For instance, if we’d like a fat-tailed dispersal kernel whose level sets are ellipses rotated by an angle *θ* counterclockwise, we can write:

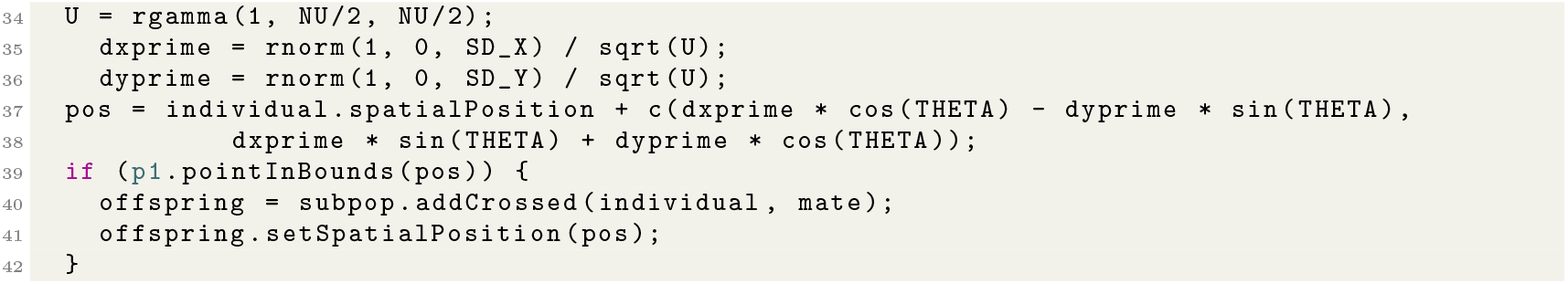

Note that we use `pointInBounds()` to check boundary condition, and so the boundary is absorbing. Here we’ve generated a *scale mixture of Normals*, by dividing our (correlated) multivariate Gaussian by 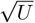, where *U* is Gamma(ν*/*2,ν*/*2) distributed. This is in fact another (and arguably better) common definition of the “multivariate Student’s *t*” (Kotz and Nadarajah, 2004).

## Appendix E Additional methodological details for case studies

### E.1 Temporal change: pikas

For computational efficiency, we focus on a 266 km^2^ region of Rocky Mountain National Park (RMNP) in Colorado (40.40363^°^N to 40.53856^°^N; 105.7326^°^W to 105.5977^°^W). A map in WGS84 projection was obtained using the `elevatr` package in R.

By simulating a restricted area of the species range, we could overestimate the probability of extinction by missing larger spatial scale population dynamics. In addition, the resulting genetic variation will certainly be affected by modeling a smaller, narrowly distributed population.

To regulate the population we at first aim for a uniform density of 250 individuals per km^2^ throughout the habitat. This value is informed by the number of pika scat piles observed by Erb et al. (2014) in sites in the Rocky Mountains. For the expected lifetime we use 3.25 years, as reported by Smith (1974). Thus, we set *K* = 250 and fecundity equal to 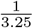.

We did not find a published value for competitive interaction distance for pikas, nor for mating distance. Therefore, we decided to use the same value for the spatial scales of competition, parent-offspring dispersal, and mating (*i.e*., *σ*_*I*_ = *σ*_*D*_ = *σ*_*M*_). Our assumption will introduce error if the competitive interaction scale in pikas is different than the dispersal scale, or if the mating scale varies from the other two values, which are both likely to be true. There is no adult movement in the pika model.

This simplifying assumption might be accurate, as pikas are territorial (Smith, 1974). To choose a value for *σ*_*X*_ (and the other, shared interaction scales) we use as a starting point the value of 300 m from Smith (1974) which was the maximum reported distance traveled by juveniles. Assuming that parent-offspring dispersal is Gaussian distributed in each dimension, we calculated a *σ*_*X*_ such that three standard deviations from the mean, Euclidean distance is 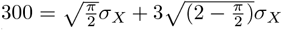 (using formulas for mean and standard deviation of the Rayleigh distribution).

### E.2 Complex life cycles: mosquitoes

In our model of mosquitoes, we assume that only the juveniles’ viability is affected by local density, while adult population size is regulated only through a constant mortality. Due to this detail, if we want to control the juvenile density to match carrying capacity, we need to modify the density control function (such as Beverton–Holt model) to reflect the life cycle.

To start our derivation, let’s say density of juveniles with age *i* is *a*_*i*_ with *i* = 1, …, *m* − 1 where *m* is maturation age. Let’s also define density of adults as *a*_*m*_. The adults have a fixed survival probability, 1 − *µ*_*a*_. We use a variation of Beverton–Holt model where survival probability of juvenile population is a local population density factor, 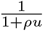, multiplied by the baseline survival probability 1 − *µ*_*j*_, where *µ*_*j*_ is the baseline mortality of juveniles. (here *u* is population density of juveniles, *i.e*., 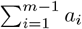).Our goal is to find *ρ*.

Then we get a system of *m* equations for **a** (left-hand-side is the *a* in the next time step, but there is no time dependence by definition of equilibrium.):

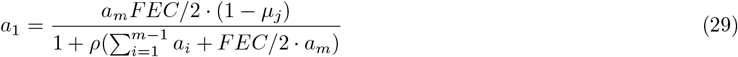

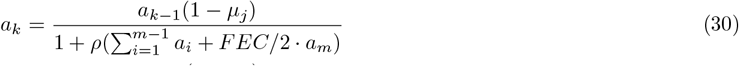

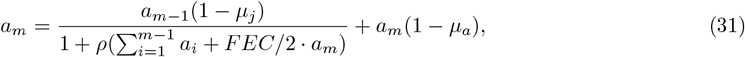

where *k* = 2, …, *m* − 1. Notice that we have a factor of 2 for fecundity, FEC, because only female adults (assumed to be half of total adult population) produces FEC new individuals. *ρ* controls where the equilibrium density is for juveniles, and it is an unknown for now. We want *ρ* to make 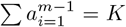,where *K* is carrying capacity of larvae.

Due to recursive relation between *a*_*i*_’s, we can simplify the system of *m* + 1 equations for *m* + 1 unknowns (*a*_*i*_’s and *ρ*) to two equations with two unknowns, *a*_*m*_ and *ρ*.

To make it a little clear, let’s define,

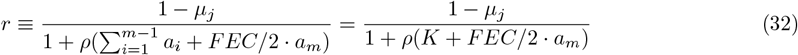

Now we start from the first equation of *a*_1_,

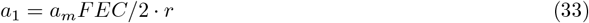

and plug it into the next one to find *a*_2_.

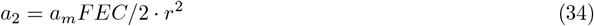

and keep going to *a*_*m*−1_. We see that *a*_*i*_ = *a*_*m*_*FEC/*2 *· r*^*m*^, a nice geometric series. (This makes sense because we tend to see exponential distributed age-structure in simulations.) This is nice because the sum can be simplified very nicely

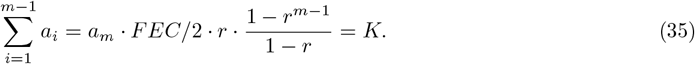

This is one of two equations we will need. The second one comes from plugging in *a*_*m*−1_ to the original equation for *a*_*m*_.

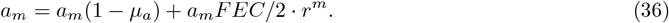

Dividing both sides by *a*_*m*_, and rearranging terms, we get

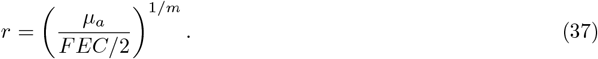

Plugging this into the first equation, we get *a*_*m*_! And we can find *ρ* by plugging it into the equation we used to define *r*. Finally, we get *a*_*m*_ and *ρ* as a function of *a*_*m*_ that we have in our simulation model:

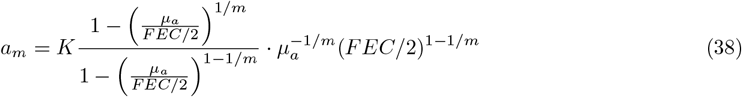

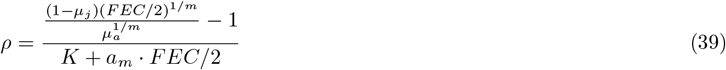

In Figure S11 we compare the expected adults to juveniles ratio from the equation above to the simulated value. Eventhough we didn’t consider other factors like spatial aspects (finding mates, migrations, heterogeneity of the river map) and seasonal fluctuation, the simulated ratio stay pretty close to theoretical expectation. In addition, I also plot the average local population density that juveniles measure around themselves through an interaction kernel with width *SX* = 20.0 in Figure S12. The density closely follows the rain factor which is a sinusoidal function added everywhere on the map to set the baseline carrying capacity, except for when the rain factor is close to zero. During the “dry season,” we see increase of density because we programmed the adult females to disperse offspring to the locations where carrying capacity is high within the maximum dispersal distance.

### E.3 Continental-scale systems: cane toads

Note that there has already been considerable effort to simulate the Australian cane toad invasion (Kearney et al., 2008), including simulations that incorporate genetic information to infer many biological parameters (Estoup et al., 2010).

Toads were simulated to have a juvenile state of 1 year. Individuals that were at least one year old were allowed to disperse once each year over their entire lifetime. Toads were randomly assigned to be male or female. Offspring initial locations were set to their mother’s locations.

Previous studies have shown that the spread rate and distribution of cane toads has been influenced by environmental heterogeneity (Urban et al., 2008). We initially used a homogeneous landscape to model the cane toad invasion, which resulted in glaring discrepancies between the simulated and observed distributions. In efforts to increase the likeness of simulated distributions to the observed data, we incorporated environmental heterogeneity across space.

The kernels controlling competition and mate choice were kept as Gaussians.

The pipeline for downloading the complete data is available (https://github.com/kr-colab/spatial_sims/blob/main/silas/range_expansion/pipelines/get_data.smk), and is described here briefly. We downloaded cane toad occurrence data was from the Global Biodiversity Information Facility (https://doi.org/10.15468/dl.8pukaa), bioclimatic variables from WorldClim (https://www.worldclim.org/data/bioclim.html (Fick and Hijmans, 2017)), and a shapefile for the geographic perimeter of Australia from the Australian Bureau of Statistics (https://www.abs.gov.au/). We converted the locations from latitude and longitude values to km using using GeoPandas (Jordahl et al., 2020) (see notebook/vignette for further details), and set the origin of the map to the earliest location in the occurrence data.

#### Extensions

Our approach could be extended to use multiple environmental maps along with a multivariate fitness function over each environmental dimension.

If desired, the simulations could be extended to include the ability to use and compare genetic information between observed and simulated data. SLiM’s ability to model explicit genomes could also allow for more complexity and realism, such as a heritable component for dispersal ability to model assortative mating for dispersal ability, as well as gene surfing, where deleterious alleles are maintained at the range edge (Miller et al., 2020; Shine et al., 2021). Additional complexity could be incorporated into the life history traits as well. For example, cannibalism has been observed in cane toads (DeVore et al., 2021). This and other modifications could be made related to changes age related competition, fecundity, mortality, and establishment.

### E.4 Resource competition: monarchs

The milkweed is only minimally simulated within this model. A number of patch centroids are randomly spread across an area of the model, and a number of milkweed plants are scattered around each centroid according to a Gaussian distribution. During each year of the model, after the monarchs have migrated south for the winter, the locations of the patch centroids and plants are re-randomized.

## Appendix F Map-based approximations to density

In this section, we provide a formal argument showing that the approximation scheme of Box 8 or of a uniform tiling of resource nodes converges to the local density for a fine enough grid (for an empirical demonstration of this convergence, see Champer et al. (2024)). Conceptually, this works because the approximation effectively computes density as if all points were at the node of the region they are in. Since all the regions are small, this does not change things much.

Suppose that the positions of individuals on the landscape are recorded as a collection of points 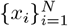.

It will be helpful for notation to represent the state of the population as a point measure, 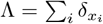,For simplicity in this section, suppose that distances are measured in units of the interaction scale, i.e., *σ*_*X*_ = 1.

The density we would like to compute from equation (1) is then

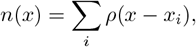

the convolution of Λ with the kernel *ρ*. Now suppose that we placed a discrete set of nodes on the landscape at locations *{y*_*j*_*}*, and for each *j* let *A*_*j*_ denote the portion of space that is closer to *y*_*j*_ than to any other node.

In other words, *{A*_*j*_*}* is the Voronoi tesselation associated with *{y*_*j*_*}*; and suppose we assign the boundaries between regions in some sensible way. Suppose that the diameters of all *A*_*j*_ are less than *ϵ*; we will show that using these nodes we can approximate *n*(*x*) well to within an error that is proportional *ϵ* – so, finer meshes of nodes will make better approximations.

Suppose that we’re evaluating density at the location of node *y*_*j*_. The approximation outlined in Box 8 seeks to approximate *n*(*y*_*j*_) by

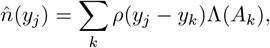

where Λ(*A*_*k*_) is the number of individuals within the region *A*_*k*_, which we may write as 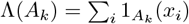.So, we can write

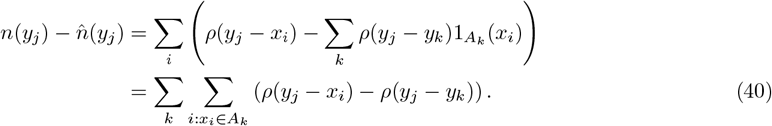

Now note that by the intermediate value theorem and the fundamental theorem of calculus,

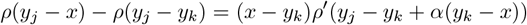

for some 0 ≤ *α* ≤ 1. Now, *ρ*(*x*) → 0 as *x* → ∞, so for any δ we may pick *R* so that

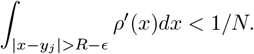

Write *N*_*R*_ = #*{i* : |*x*_*i*_ − *y*_*j*_| ≤ *R}* for the number of points closer than *R*. Furthermore, suppose the derivative of *ρ* is bounded above by *C, i.e*., *ρ* ^*′*^ (*x*) ≤ *C*. If |*x* − *y*_*k*_| ≤ *ϵ*, then

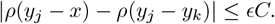

Since for *x*_*i*_ ∈ *A*_*k*_, by definition |*x*_*i*_ − *y*_*k*_| ≤ *ϵ*, plugging this into equation (40), and splitting the sum into regions with nodes further than *R* away and from *y*_*j*_ and not,

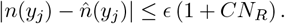

Since *C* and *N*_*R*_ are fixed, this goes to zero as *ϵ* → 0. This shows that 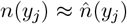;since 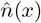 is defined for arbitrary *x* by interpolation, 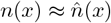 as well.

## Supplementary figures

**Figure S3:**
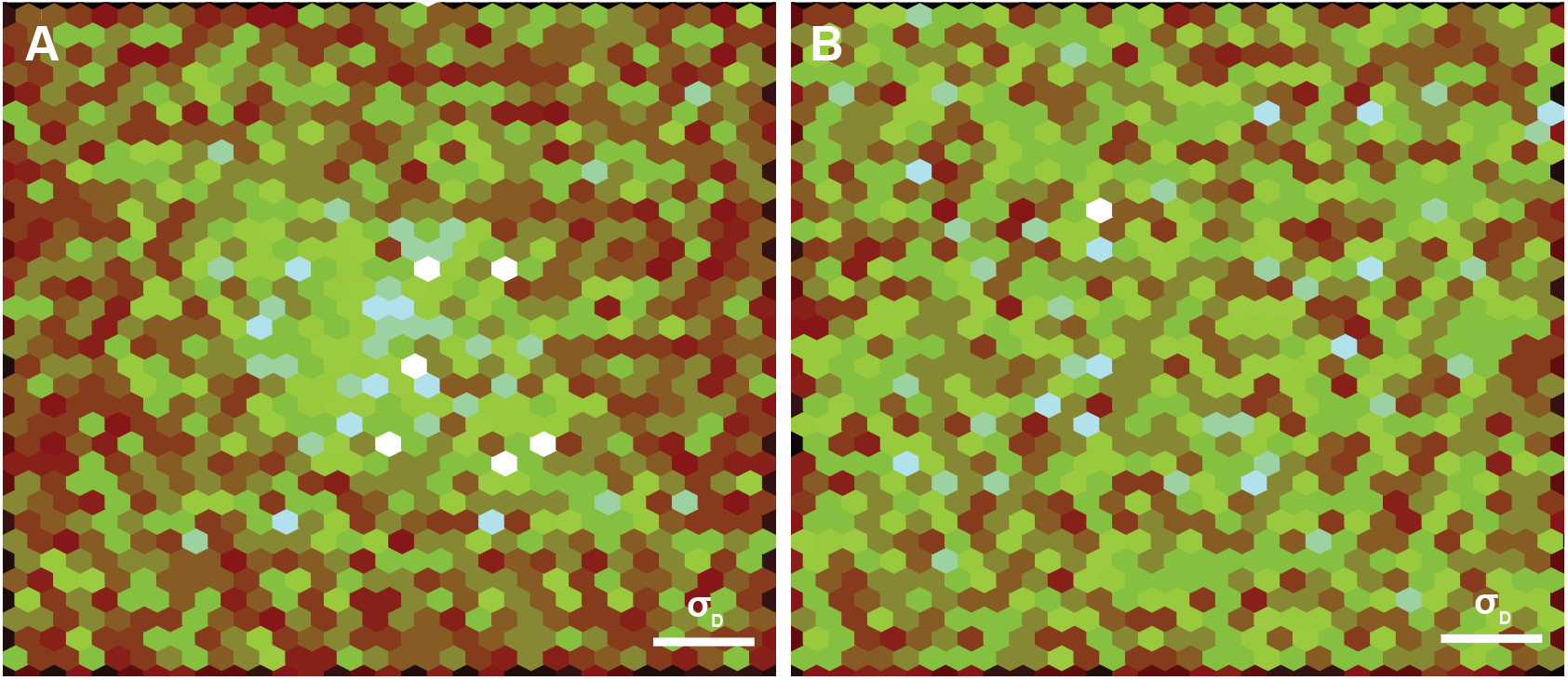
Fatter-tailed dispersal reduces clumping. Fry plots illustrate the distance between each pair of individuals at a given tick. The dense region of points near the origin in the Gaussian panel indicates many pairs of individuals are separated by small distances. Points shown are pairwise distances ≤ 1 from an arbitrary time step from one replicate. Here, *σ*_*D*_ = 0.3. **(A)** Gaussian dispersal kernel. **(B)** Student’s *t* dispersal kernel. Figure 3CD quantify clumping visualized here.

**Figure S4:**
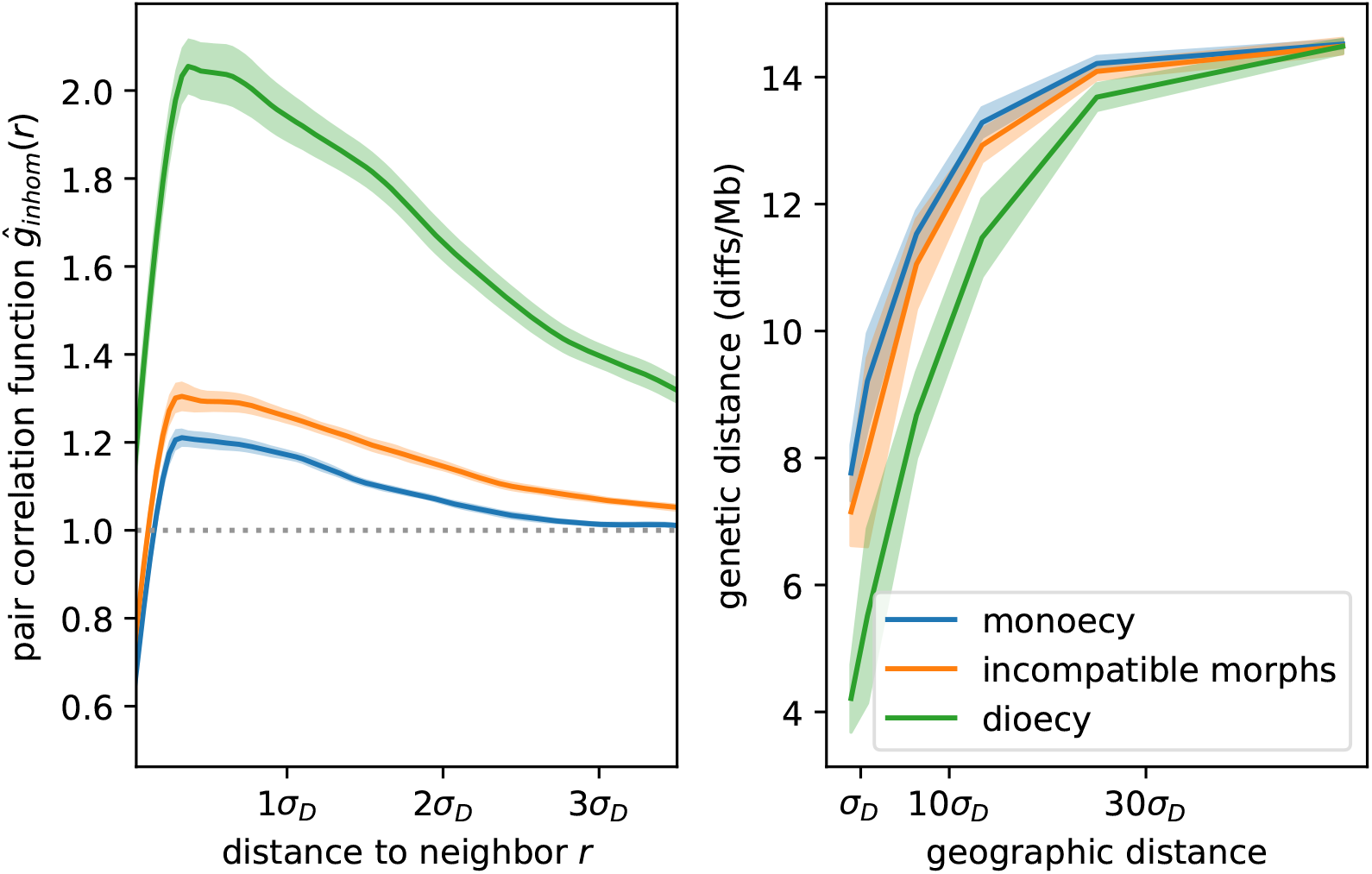
Mating type affects magnitude and spatial scale of clumping. **(left)** Pair correlation functions show density of pairs of individuals found a particular distance apart, relative to distances expected under a Poisson process (1.0; grey dotted line). Curves show the average across 50 independent time steps. Mating types are as described in Section 5. **(right)** Mate limitation increases genetic isolation-by-distance. Plots show mean genetic distance between pairs of individuals at increasing geographic distance, averaged across ten independent replicates. Figure 3AB visualize clumping quantified here for monoecious and dioecious scenarios, respectively.

**Figure S5:**
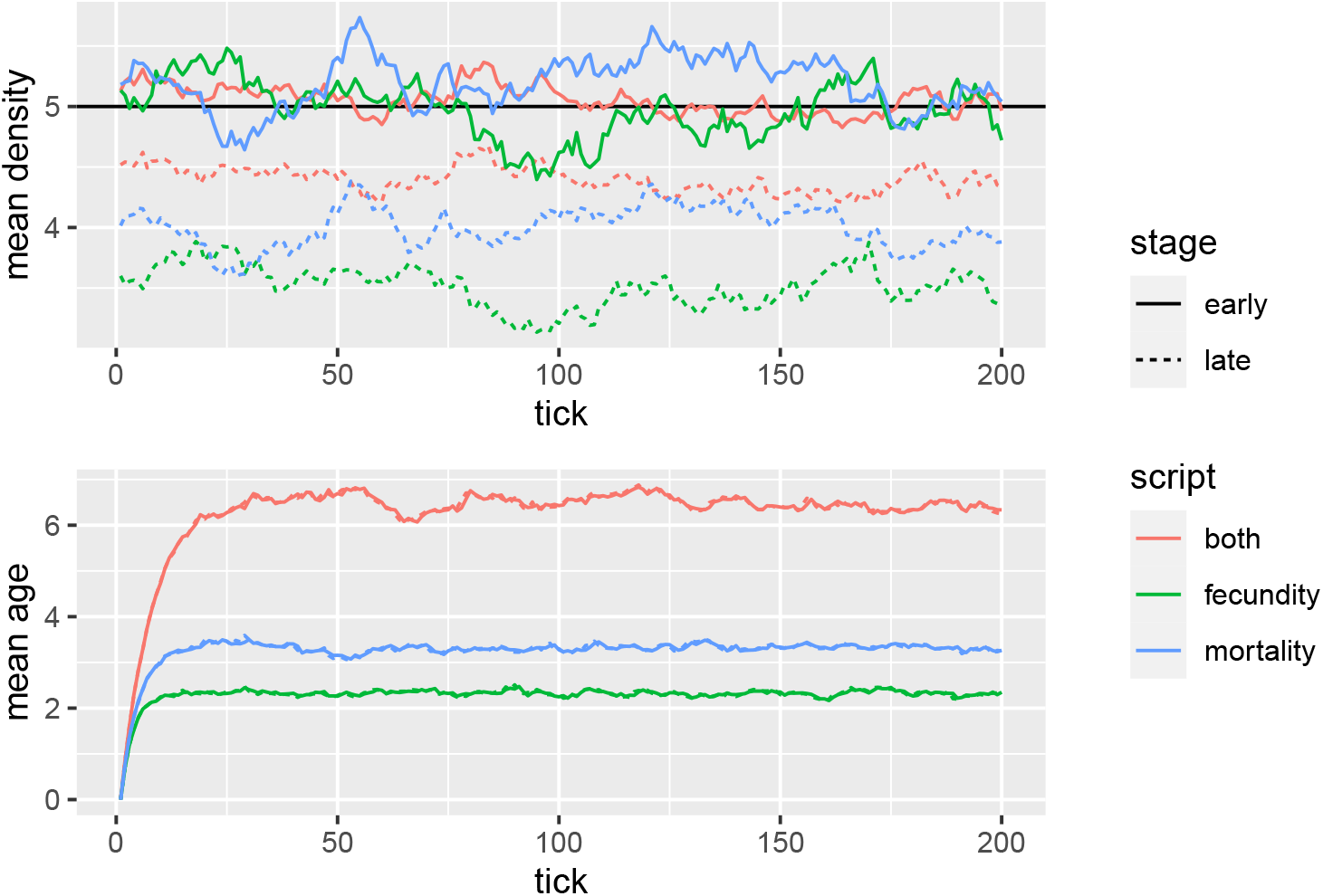
Traces from spatial simulations of the three “Beverton–Holt” models shown in Figure 1. **(Top)** Average local density for all individuals, measured both between birth and death (“early”) and between death and birth (“late”), and **(bottom)** mean ages across 200 time steps. Horizontal line shows the value of *K* = 5; local population density determines population regulation between birth and death for all models. Other parameters: *σ*_*D*_ = *σ*_*X*_ = 1.2, and range was a 25 *×* 25 square area.

**Figure S6:**
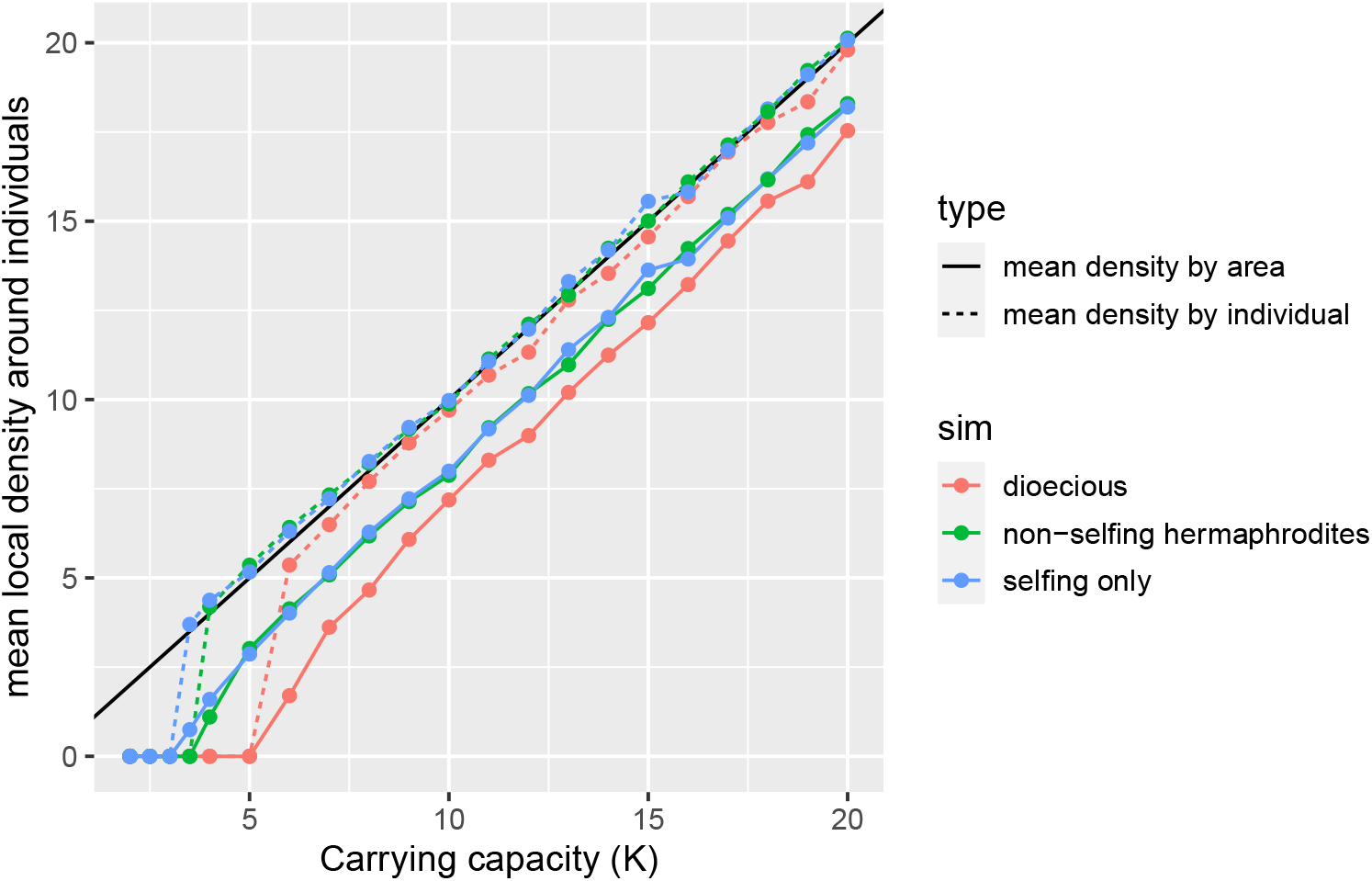
Average realized population density at equilibrium of three different models, plotted against the “carrying capacity” parameter, *K*. Each model uses Beverton–Holt regulation of mortality with the same parameters (*σ*_*D*_ = *σ*_*I*_ = *σ*_*M*_ = 0.3). and differ only in that “*dioecious*” individuals are one of two sexes: only females reproduce and only if they mate with a male; “*non-selfing hermaphrodites* “ also must mate with another individual, but all individuals can reproduce; and “*selfing*” individuals can all reproduce and have no need for mating. Furthermore, mean fecundity is *f* = 0.5 for the dioecious simulations and *f* = 0.25 for the others. Dotted lines show mean local density experienced by individuals; solid lines show total population size divided by total area. All simulations die out at low density, but those requiring mating die out at higher *K*: in each case, when *N*_*M*_ is around 1.6.

**Figure S7:**
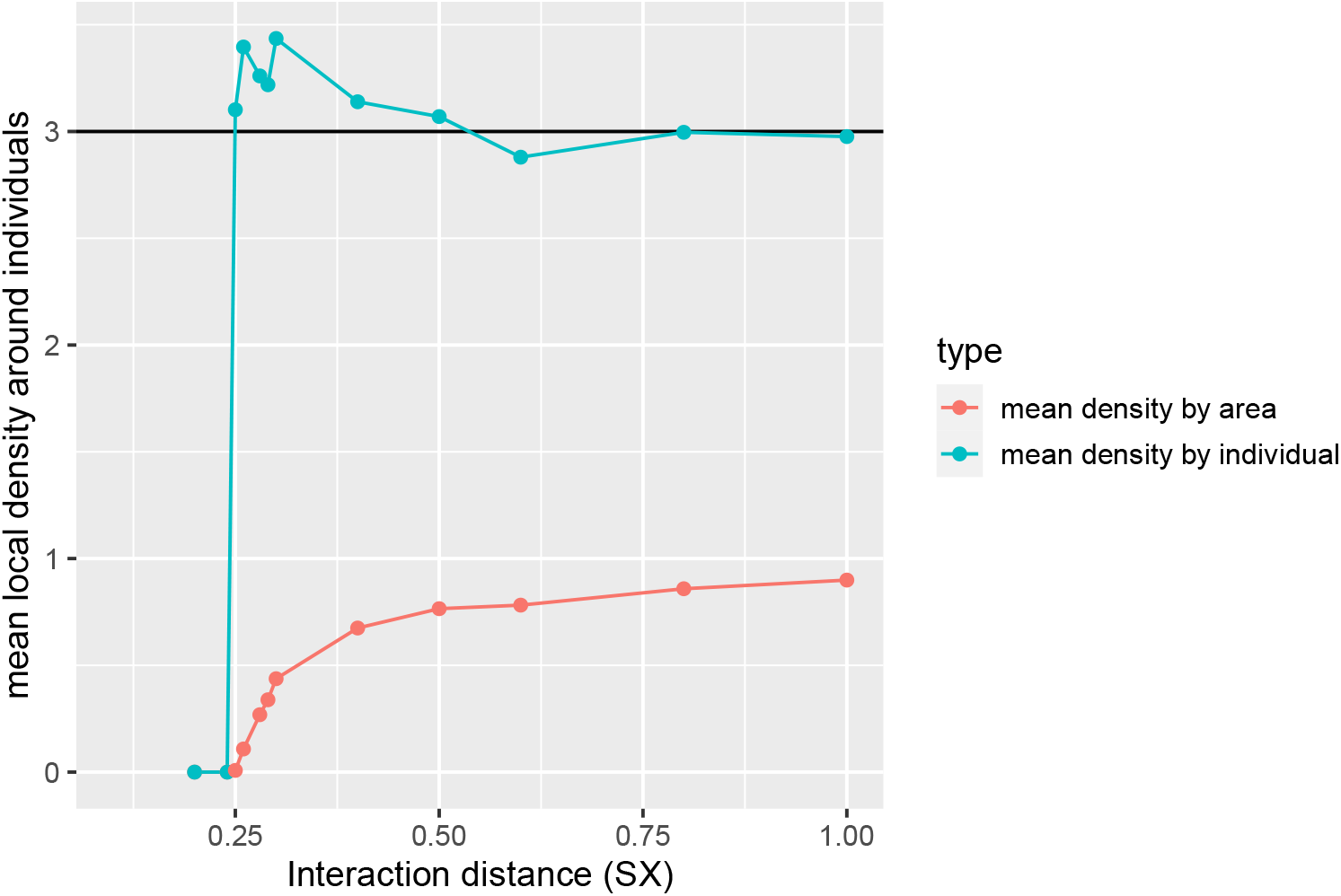
Mean density at equilbrium in simulations with different values of *σ*_*X*_ (`SX` in the figure). The blue line shows mean density around individuals (*i.e*., the average of local density calculated with equation (1) for all individuals), the red line shows total population size divided by area (a 25 *×* 25 box), and the horizontal line is at *K* = 3. The simulation uses Beverton–Holt control of mortality with *σ*_*D*_ = 0.5, reproduction entirely through selfing, and no adult movement. The population dies out if *σ*_*X*_ is below 0.25, while for 0.25 ≤ *σ*_*X*_ ≤ 0.5, total density (*i.e*., density averaged by area) increases, while mean density experienced by individuals is larger than *K* and decreases.

**Figure S8:**
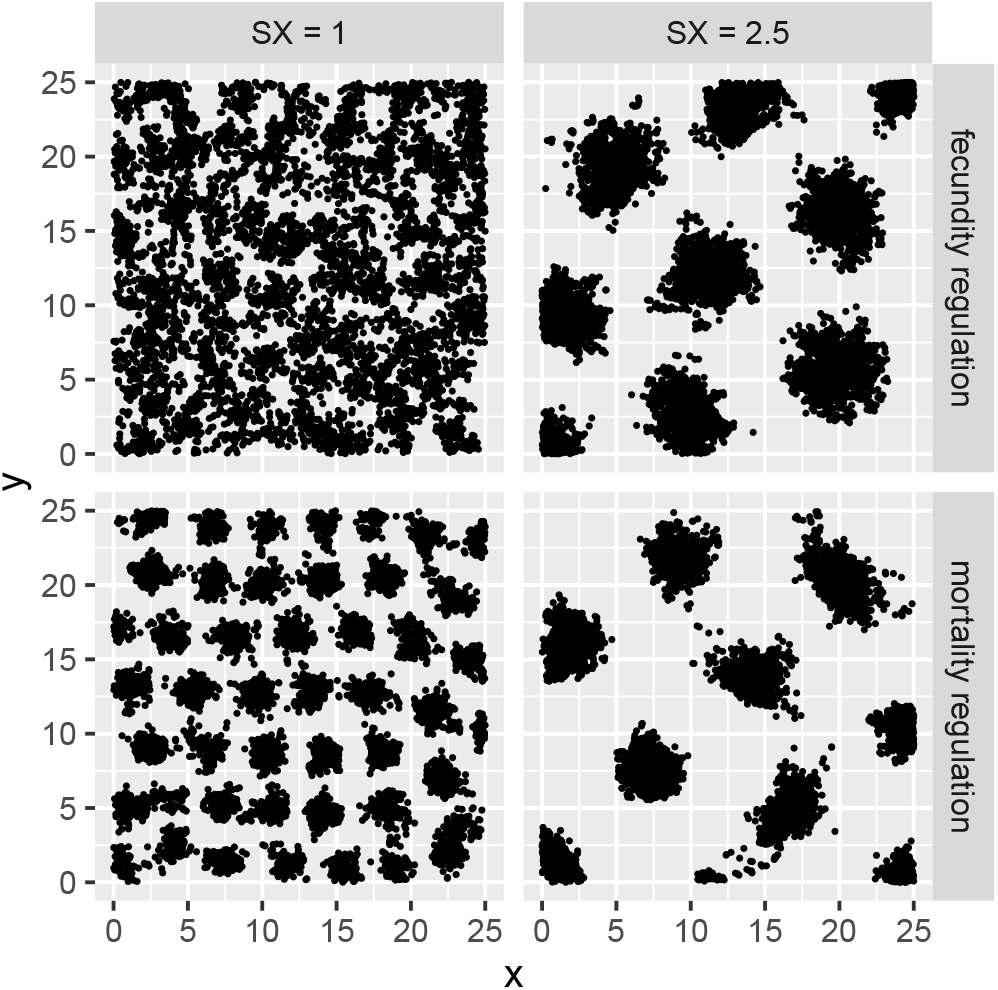
Examples of simulations exhibiting clumping, using *σ*_*D*_ = 0.2, two different values of *σ*_*X*_ (labeled `SX`), and two different types of density-dependent feedback (either mortality or fecundity regulation, as in Figure S5).

**Figure S9:**
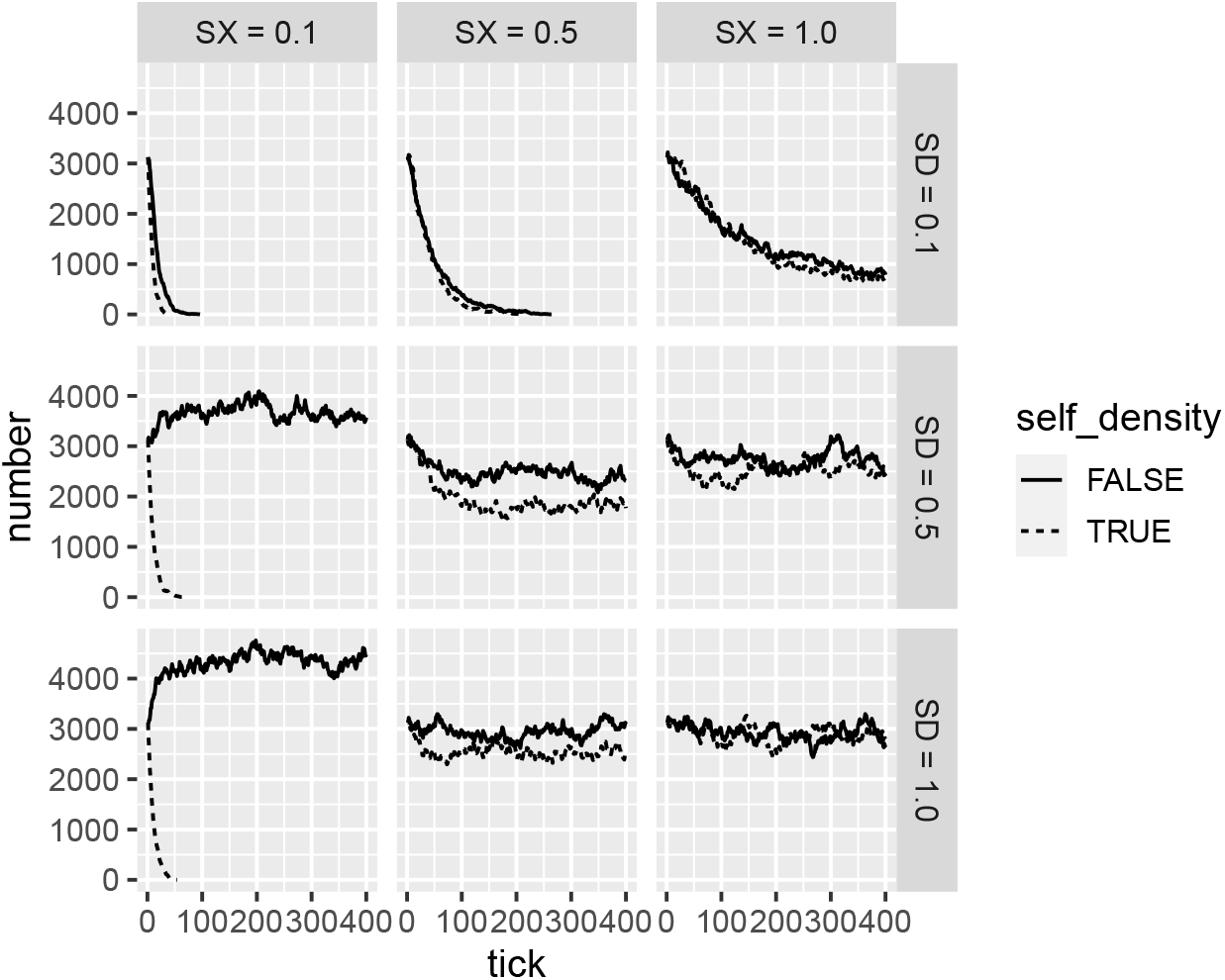
Population sizes through time for simulations with various values of dispersal scale (*σ*_*D*_, here `SD`) and interaction scale (*σ*_*X*_, here `SX`), with or without inclusion of the focal individual in local density calculations. Each simulation had Beverton–Holt density-dependent feedback on mortality (as in Figure S5), and was run with *K* = 5 on a 25 *×*25 square area, and were started with 5 *×*25 = 3125 individuals (so, lines that are roughly flat are fluctuating around a total density of *K* = 5). Solid lines compute “local density” for control of mortality of each individual using equation (1), while dotted lines do the same except excluding the focal individual.

**Figure S10:**
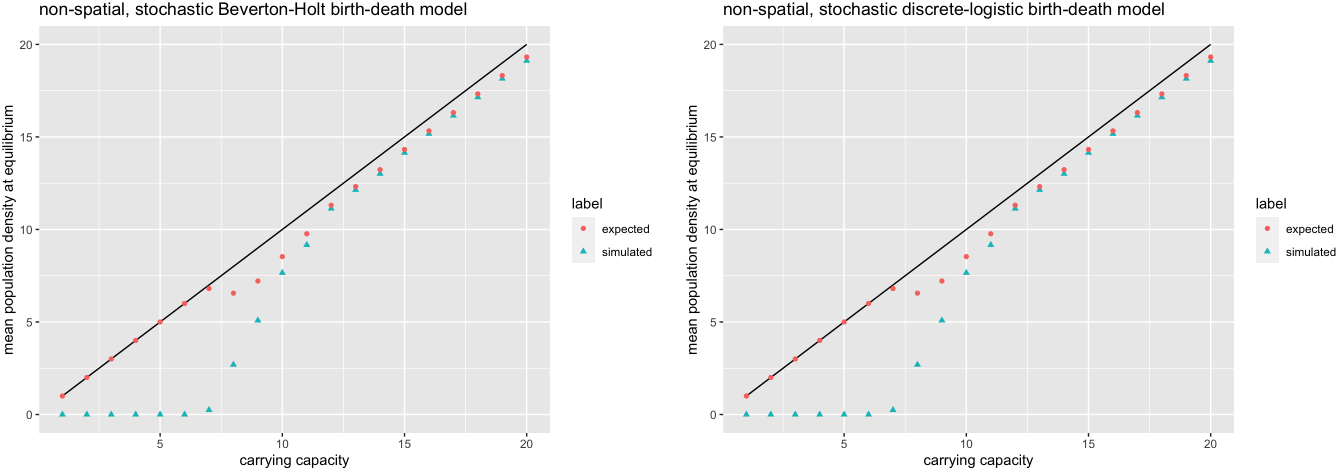
Expected population density obtained using equation (11) and actual average population density in nonspatial models with **(left)** Beverton–Holt regulation of mortality for which “expected” is 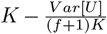,and **(left)** discrete logistic regulation of mortality, for which “expected” is 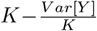.Nonspatial simulations were run in R. Solid line is where density is equal to carrying capacity, which is true for deterministic model.

**Figure S11:**
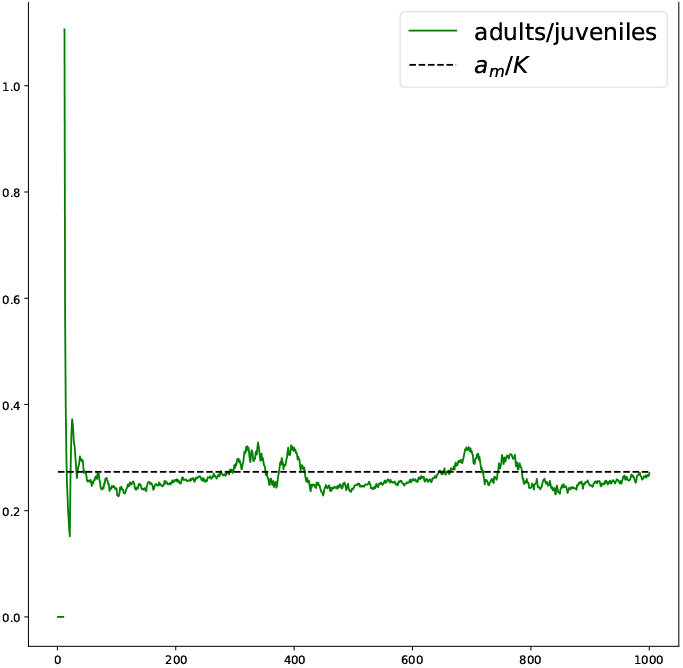
Ratio between adult and juvenile mosquito counts.

**Figure S12:**
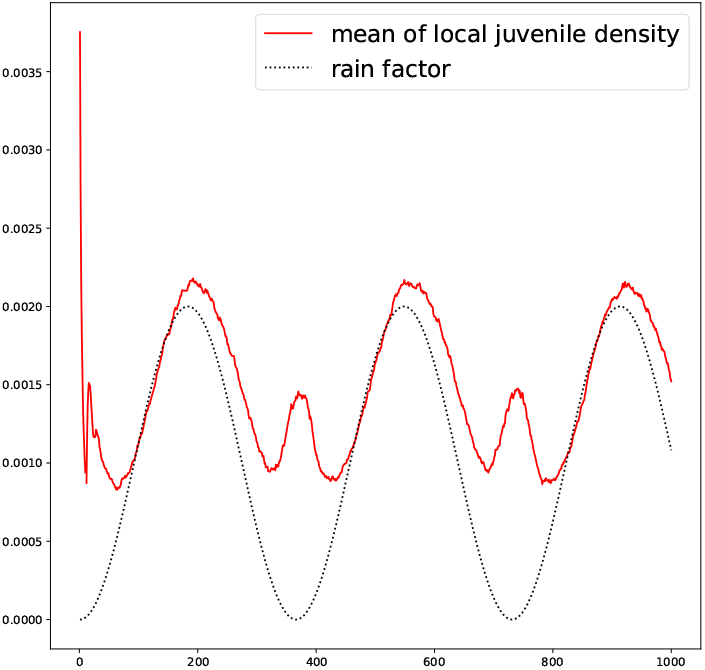
Local population density of juvenile mosquitoes plotted against rain factor, which is minimum carrying capacity at each time point.

